# scMicrobe PTA: Near Complete Genomes from Single Bacterial Cells

**DOI:** 10.1101/2024.01.30.577819

**Authors:** Robert M Bowers, Veronica Gonzalez-Pena, Kartika Wardhani, Danielle Goudeau, Matthew James Blow, Daniel Udwary, David Klein, Albert C Vill, Ilana L Brito, Tanja Woyke, Rex Malmstrom, Charles Gawad

## Abstract

Microbial genomes produced by single-cell amplification are largely incomplete. Here, we show that primary template amplification (PTA), a novel single-cell amplification technique, generated nearly complete genomes from three bacterial isolate species. Furthermore, taxonomically diverse genomes recovered from aquatic and soil microbiomes using PTA had a median completeness of 81%, whereas genomes from standard amplification approaches were usually <30% complete. PTA-derived genomes also included more associated viruses and biosynthetic gene clusters.

## MAIN TEXT

Difficulties in cultivating most bacterial and archaeal species presents a barrier to exploring the genetic make-up of the Earth’s microbiomes. To access the genomes of most microorganisms, culture-independent methods such as shotgun metagenomic sequencing ^1–3^ and single-cell sequencing ^4–8^ can be employed. While metagenomics has led to unprecedented insights into the metabolic potential of uncultured microorganisms ^9–12^, the approach has some limitations. For example, it is difficult to connect mobile genetic elements such as plasmids and phages to metagenome-assembled genomes (MAGs) ^13^. Generating MAGs from heterogeneous or low abundance populations is also challenging ^14,15^. Single-cell sequencing, in contrast, does not share these same limitations ^5^, and the approach has provided insights into microbial dark matter ^4,7^, experimentally linked phages to their hosts ^16,17^, and dissected natural populations ^13,18,19^. However, multiple displacement amplification (MDA) – the predominant single-cell genome amplification method ^20^ – is limited by the poor uniformity and completeness of the genomes it produces ^21^. Single-cell amplified genomes (SAGs) typically have genome completeness <=40% ^4^.

Different variations on genome amplification chemistry ^22–24^ and sample processing strategies ^25–30^ have improved genome recovery in some situations, but an approach for consistently generating complete or nearly complete genomes from single microbial cells is still lacking. We recently developed primary template-directed amplification (PTA), which significantly improves amplified genome uniformity and variant calling in single human cells ^31^. Here, we investigated whether PTA could also improve the quality of genomes recovered from single bacterial cells.

To benchmark PTA performance against the genome amplification chemistries commonly used in microbiome studies, we first sequenced the genomes of three bacterial isolate species: *Escherichia coli* (Gram-), *Pseudomonas putida* (Gram-), and *Bacillus subtilis* (Gram+). Individual cells were sorted into 96-well plates using fluorescence activated cell sorting (FACS), and replicate plates were subjected to genome amplification using PTA, MDA, and WGA-X, a modified version of MDA that uses a more thermostable variant of phi29 polymerase ^23^ (Supplementary Fig. 1). Sequencing reads were mapped to reference genomes to measure coverage uniformity, and later assembled *de novo* using SPAdes ^32^. All libraries were sub-sampled to 1M reads prior to these analyses to ensure comparable sequencing effort among SAGs.

In every case, genome coverages from PTA reactions were significantly more uniform than MDA and WGA-X reactions based on Lorenz curves and Gini coefficients (Fig. 1; p<0.01 one way ANOVA and Tukey HSD for *E. coli* and *P. putida*; p<0.01 one way t-test for *B. subtilis*). In addition, PTA amplification resulted in significantly greater genome completeness than did MDA and WGA-X for all three species (Fig. 1C; p<0.01 one way ANOVA and Tukey HSD for *E. coli* and *P. putida*; p<0.01 one way t-test for *B. subtilis*). For example, *B. subtilis* and *E. coli SAGs* assembled de novo had an average completeness of 94% and 91%, respectively, whereas genomes generated by MDA recovered only 60% and 62% on average. *P. putida* SAGs were less complete for all chemistries, but genomes generated by PTA were nearly 2-fold more complete than those generated by MDA and WGAX. *P. putida* genome completeness improved to 91% after increasing the number of input reads to an average of 4M. PTA also showed similar fidelity to MDA and WGA-X when copying the genomes, e.g., no significant difference in genome mismatch rates per 100 kilobases among amplification chemistries (Fig. 1C; p > 0.05 one-way ANOVA). Overall, these results mirror the superior performance of PTA versus MDA and other genome amplification strategies observed previously using human cells ^31^.

**Figure 1.**
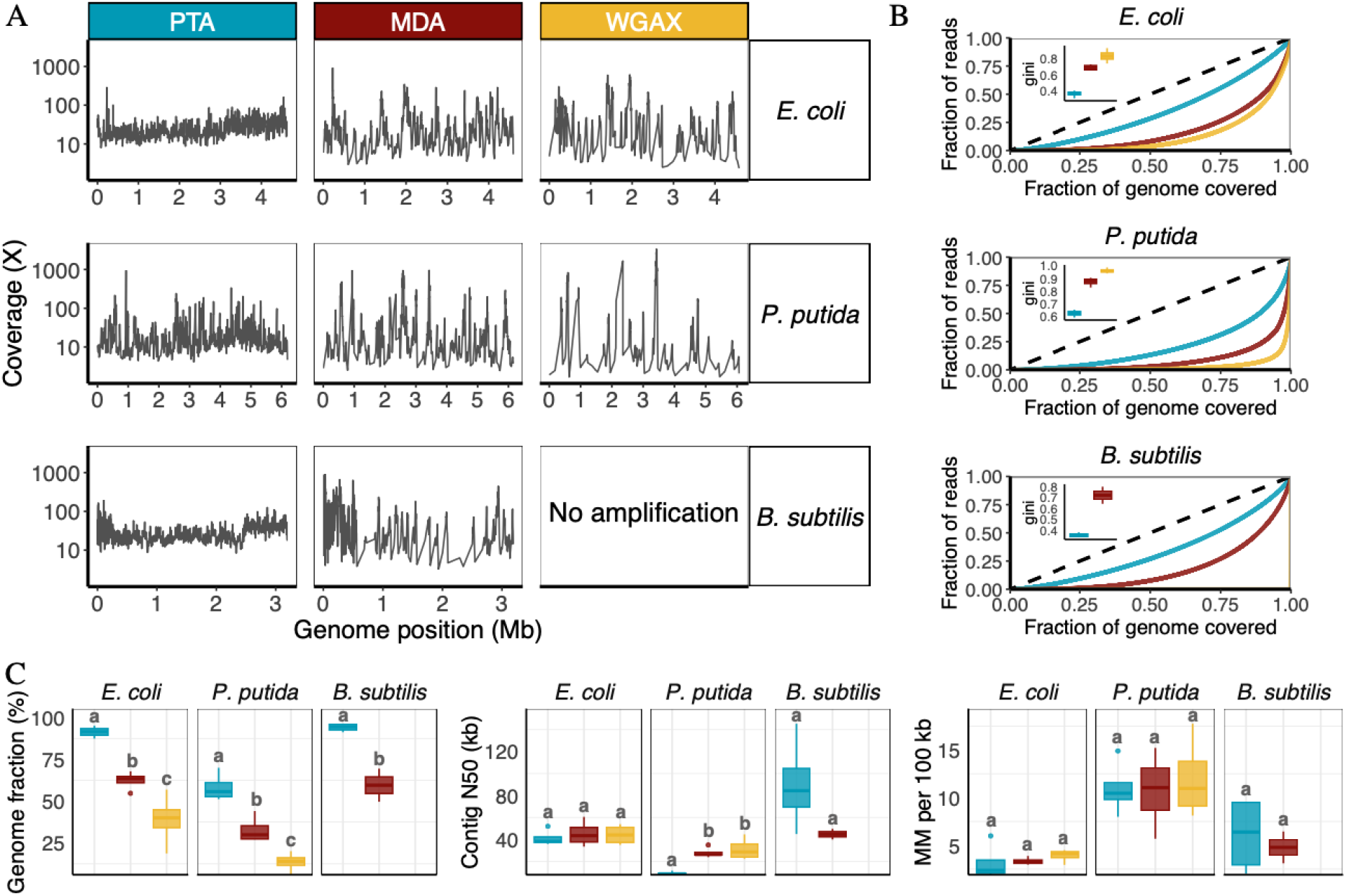
Genome quality of E. coli, P. putida, and B. subtilis SAGs amplified using PTA (blue), MDA (red), and WGA-X (yellow). **A)** Genome coverage of 500 bp windows from one representative replicate of each species amplified with each chemistry. WGA-X amplification reactions of *B. subtilis* failed and were excluded from further consideration. Refer to Supplementary Fig. 2 for genome coverage plots of all replicates. **B)** Uniformity of genome coverage illustrated by Lorenz Curves and Gini Coefficients. The dotted line represents the expected pattern of perfect uniform coverage, and solid lines illustrate the observed coverage for representative cells. **C)** Key summary statistics of *de novo* genome assemblies including completeness, contig N50, and the number of mismatches (MM) per 100 Kb. The letters a, b, and c above the boxplots denote significance at the alpha 0.05 level. Sample sizes are n = 4 for all species and chemistries except for MDA amplified *B. subtilis*, which had n = 2. The boxplot dots represent outliers that are beyond the 1.5-fold the interquartile range. Additional summary statistics are reported in Supplementary Fig. 3 and Supplementary Tables S1 and S2.

After performing these benchmarking experiments with bacterial isolates, we sought to determine if the improved performance of PTA could be extended to environmental samples. To accomplish this, we utilized the same comparison strategy to amplify and sequence single cells recovered by FACS from aquatic and soil samples (Supplementary Fig. 4). We again found that PTA resulted in significantly greater genome completeness than MDA and WGA-X (Fig 2A and Supplementary Table S3; p<0.01 one way ANOVA and Tukey HSD). For example, PTA reactions from aquatic samples had median genome completeness of 83%, while completeness from MDA and WGA-X reactions had medians of 17% and 11%, respectively (Fig. 2A and Supplementary Table S4). Deeper sequencing of MDA and WGA-X libraries to approximately 20M reads increased median completeness estimates to 30% and 23%, respectively (Supplementary Table S5), but these genomes were still far less complete than those derived from PTA reactions (p<0.01 one way ANOVA and Tukey HSD). Similar patterns were observed from a smaller soil microbiome dataset where PTA produced genomes with much greater completeness than MDA and WGA-X (Fig. 2A; p<0.01 one way ANOVA and Tukey HSD). Additionally, a larger fraction of PTA genomes recovered from the aquatic system had virus and biosynthetic gene clusters (BGC) sequences, and a larger fraction of PTA genomes from soil had plasmid sequences (Fig. 2A; p<0.05 Fisher’s exact test). Finally, phylogenetic analysis revealed successful PTA reactions on cells belonging to 20 families spread across 6 phyla (Fig. 2B), suggesting that PTA is amenable to a wide variety of microorganisms and produces substantially more near-complete genomes than standard amplification chemistries used in microbiome studies (Fig. 2, Supplementary Fig. 5).

**Figure 2.**
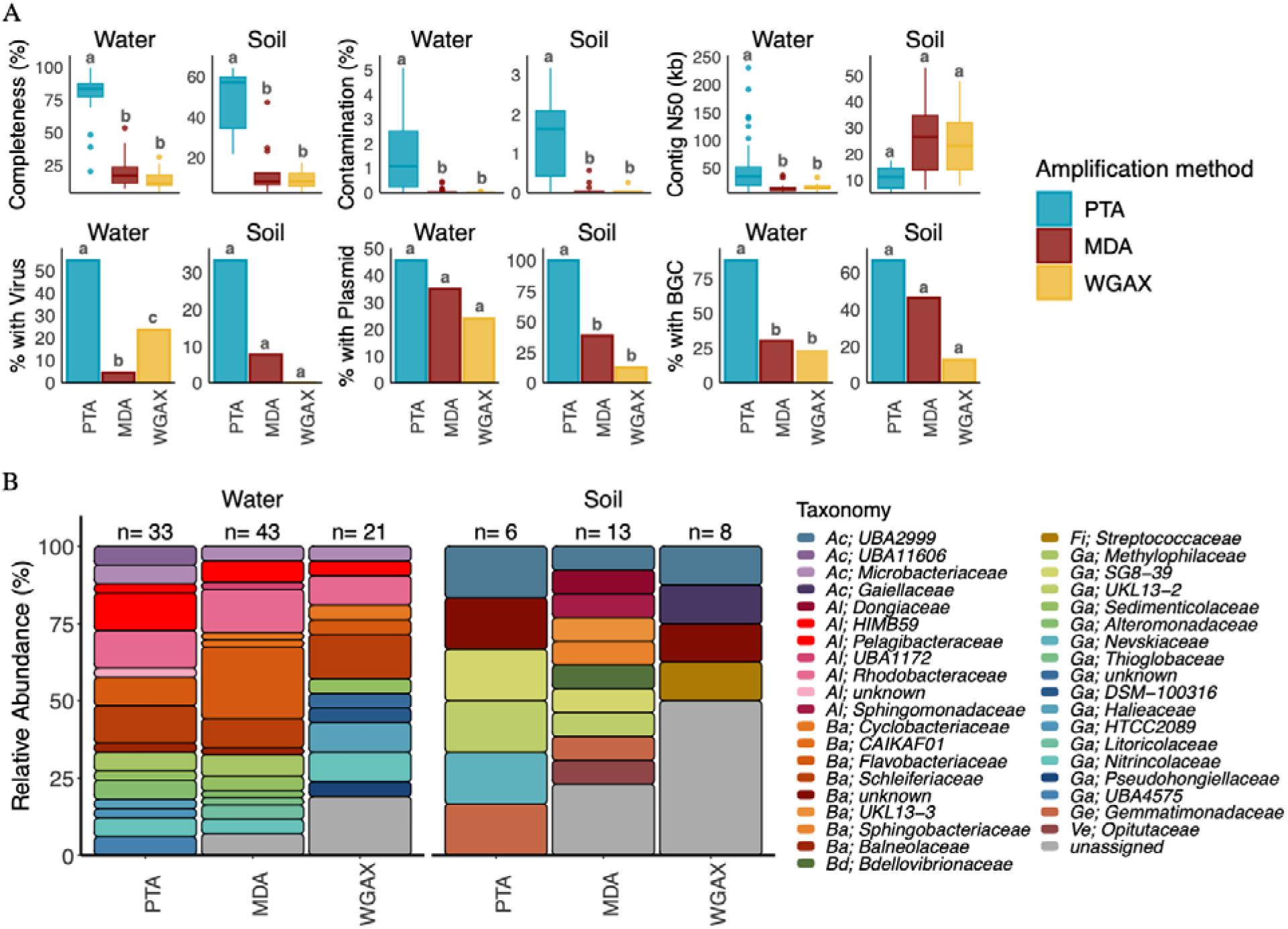
Comparison of SAGs from aquatic and soil microbiomes amplified with PTA (blue), MDA (red), and WGA-X (yellow). **A)** Estimated genome completeness and contamination, contig N50, and the percentage of SAGs containing at least one predicted plasmid (> 5 kb), virus (> 5 kb), or BGC. The letters a, b, and c denote significance at the alpha 0.05 level. **B)** Family level taxonomic assignment of SAGs assembled from <=20 Mio reads. Phylum / Class abbreviations are as follows: Ac: Acidobacteria, Al: Alphaproteobacteria, Ba: Bacteroidota, Bd: Bdellovibrionota, Fi: Firmicutes, Ga: Gammaproteobacteria, Ge: Gemmatimonadota, Ve: Verrucomicrobiota.

For single-cell genomes, overall genome quality is measured by a combination of completeness and contamination, with “high-quality” genomes defined as having >90% completeness and <5% contamination ^33^. Environmental genomes generated by MDA and WGA-X had median estimated contamination levels of < 0.1%, whereas PTA genomes had a median of 1.5% after applying an informatic decontamination procedure. We observed the same contaminating sequences across many SAGs and in the no-template control reactions amplified by PTA, suggesting the PTA reagents contained trace levels of contaminating DNA. Single-cell whole genome amplification chemistries use short random primers to amplify a few femtograms of DNA, so even trace amounts of contaminating DNA can appear in assemblies. To decrease contaminating DNA, MDA and WGA-X reagents underwent secondary treatment with UV prior to genome amplification34 while PTA reagents had initial decontamination done during manufacturing but not secondary UV treatment, which may explain the slightly higher contamination levels observed. It is also possible that PTA is detecting contaminating DNA that is not captured with other methods. Nevertheless, PTA was the only chemistry to produce high quality SAGs from the environmental samples (Supplementary Fig. 5).

In summary, we present scMicrobe PTA, the application of PTA to greatly improve genome recovery of single bacterial cells growing in culture as well as those directly sorted from environmental microbiomes. These results set the stage for a renaissance in single-cell-based environmental genomics by offering a more comprehensive insight into the population structure of the microbial dark matter that accounts for a large fraction of the Earth’s biomass.

## METHODS

### Sample Collection and Processing

Fresh cultures of *Escherichia coli* MG1655, *Pseudomonas putida* KT2440, and *Bacillus subtilis* pDR244 were grown overnight in LB at 37 °C, then used immediately for cell sorting as described below. An aquatic sample was collected from the surface waters of Mountain View Slough (latitude 37.432400, longitude -122.086632). The sample was vortexed for 15 seconds to release cells attached to sediment, filtered using a 15 um cell strainer (pluriStrainer from pluriSelect, Germany) to remove large particles, and stored in 25% glycerol at -80C until sorting. A soil microbiome sample was collected at Lawrence Berkeley National Laboratory (latitude 37.877382, longitude -122.250410). The soil sample was vortexed for 15 seconds to release cells attached to sediment, then centrifuged at 500g for 5 minutes to pellet large particles. The supernatant was used immediately for cell sorting.

### Fluorescence Activated Cell Sorting (FACS)

Immediately before cell sorting, environmental bacteria and bacterial isolates were filtered through a 35 μm cell strainer to remove large debris and cell clusters and diluted to approximately 10^6^ cells/ml in filter-sterilized 1X PBS containing 1X SYBR-Green DNA stain (Thermofisher, USA). Individual cells were sorted using an Influx FACS machine (BD Biosciences) into LoBind 96-well plates (Eppendorf, Germany) containing either 3 μL of BioSkryb SL1-B Solution for PTA reactions or 1.2ul of TE for MDA and WGA-X reactions. Plates were treated for 10 minutes in a UV crosslinker before sorting to remove any contaminating DNA. Cells were discriminated based on a combination of forward scatter characteristics and SYBR Green fluorescence. A single-cell sort mask with extra droplet discrimination was used to ensure only one cell was sorted into each well.

### Whole Genome Amplification

PTA was performed using the ResolveDNA Bacteria kit (BioSkryb Genomics, USA) with a few changes. Briefly, 3uL of SL-B reagent (BioSkryb Genomics, USA) was deposited in each well of a LoBind twin.tec PCR plate (Eppendorf, Germany) prior to sorting. Plates containing sorted cells were film-sealed, briefly spun, mixed in a Thermomixer C (Eppendorf, Germany) at 1400 rpm for 1 minute, and briefly spined again. The plates were then incubated at room temperature for 30 minutes and stored at -80 °C until ready to use. PTA DNA amplification was carried as per BioSkryb Genomics protocol for 12 hours at 30 °C, followed by 3 minutes at 65 °C to stop the reaction (ResolveDNA Bacteria Protocol PN100294). Amplified DNA was cleaned using SeraMagSelect beads at a 2X beads to sample ratio (Cytiva Life Sciences, USA).

MDA was performed using Phi29 DNA Polymerase (Watchmaker Genomics, USA) as described previously ^5^ with 20uL reaction volumes to match PTA reaction volumes. In addition, a subset of libraries received an additional Ready-Lyse (LGC Biosearch Technologies) lysozyme treatment of 50U/ul for 15 minutes prior to alkaline lysis (Supplementary Tables S5 and S6). Similarly, 20uL WGA-X ^23^ reactions were performed with EquiPhi29™ DNA Polymerase (Thermo Fisher).

### Library Preparation and Genome Sequencing

Sequencing libraries were prepared from 10 -100 ng input DNA using the Nextera DNA flex library prep (Illumina, USA) using IDT for Illumina – Nextera DNA UD Indexes Sets A-D (Illumina, USA). Fragmentation times and amplification cycles were performed according to the ranges recommended by the manufacturer. Amplification reactions were cleaned using SPRI beads (Beckman Coulter, USA) at a 2X beads-to-sample ratio. Library concentrations and sizes were analyzed by TapeStation 2200 using D1000 ScreenTapes (Agilent, USA), and library concentration was determined using a Qubit fluorometer with DNA High Sensitivity reagents (Thermofisher, USA). Bacterial isolates and a subset of the aquatic environmental cells were sequenced on the NextSeq 2000 (Illumina), while the remainder libraries from aquatic and soil bacteria were sequenced on the Novaseq 6000 (Illumina) (Supplementary Tables 5 and 6). All libraries were sequenced using a 2X150bp read format.

### Read Processing and Genome Assembly

Sequencing reads were filtered for quality using the rqc.filter2.sh script from BBTools Version 39.01 (https://bbtools.jgi.doe.gov) with following parameters: rna=f trimfragadapter=t qtrim=r trimq=6 maxns=1 maq=10 minlen=49 mlf=0.33 phix=t removehuman=t removedog=t removecat=t removemouse=t khist=t removemicrobes=t sketch kapa=t clumpify=t rqcfilterdata=/clusterfs/jgi/groups/gentech/genome_analysis/ref/RQCFilterData barcodefilter=f trimpolyg=5

To generate assemblies from high and low levels of sequencing effort, each library was first subsampled to a maximum of 20M and 1M quality filtered reads. Each subsampled library version was then normalized using bbtools.bbnorm with parameters: bits=32 min=2 target=100 pigz unpigz ow=t. This normalization reduces the massive redundancy of reads from highly covered genome regions. Error correction was done on the normalized fastq using bbtools.tadpole with parameters: mode=correct pigz unpigz ow=t. Normalized reads were assembled using SPAdes v3.15.3 ^32^ using parameters: --phred-offset 33 -t 16 -m 64 --sc -k 25,55,95.

Assembled contigs were trimmed to remove 200bp from the the beginning and ending of each contig, and contigs < 2,000bp were removed.

### Genome Quality Assessment and Taxonomic Classification

The quality of SAGs derived from isolates was determined using QUAST version 5.2.0 ^35^. Because sequencing effort varied substantially among bacterial isolate SAGs, assemblies made with 1M reads were compared so that all replicates had equivalent sequencing depths. Genome coverage levels were determined by mapping each of the isolate SAGs against its corresponding reference genome: *E. coli* (IMG taxon ID: 2600254969), *P. putida* (IMG taxon ID: 2667527229) and *B. subtilis* (IMG taxon ID: 643886132). The bbmap parameters used in the analysis were bbmap.sh -Xmx100g fast=t 32bit=t. The resulting bam files were passed bedtools (v2.31.0) ^36^ to generate coverage files using the genomecov function. Lorenz curves and Gini coefficients were calculated from genomecov files using the R package gglorenz (v0.0.2). The Gini coefficient quantifies the observed deviation from perfect uniformity for each replicate cell, with smaller coefficients indicating more uniform coverage ^37^.

Environmental SAG assemblies were screened for contamination using a stepwise approach. First, we removed any human contigs. Next, we applied MAGpurify (https://github.com/snayfach/MAGpurify) in two sequential stages to remove contaminant contigs based on GC content and phylogenetic markers (stage 1) and tetranucleotide signatures (stage 2). Following the MAGpurify cleanup, we mapped reads generated from negative control reactions that lacked sorted cells and removed contigs with coverage > 5X. Finally, we ran megablast against the NCBI non-redundant database and removed contigs with top hits to a set of organisms consistently found in the negative control reactions. Informatic decontamination reduced median contamination estimates for PTA SAGs from roughly 3% to 1.5% in genome versions assembled from 1M reads. Decontamination had little to no impact on MDA and WGA-X SAGs whose contamination levels were <0.1% before treatment. Following contaminant removal, the quality of the environmental SAGs was assessed with CheckM2 (v1.0.1) ^38^.

Statistical analysis of proportional results such as Gini coefficients, genome completeness, and genome contamination were performed on arcsine transformed data.

Taxonomic assignments of environmental SAGs were made with GTDB-tk (v2.3.2) ^39^. SAGs derived from 20M reads were used, when available, for taxonomic analysis because GTDB-tk struggled to make assignments to the less complete MDA and WGA-X genomes generated with 1M reads.

### Identification of Viruses, Plasmids and Biosynthetic Gene Clusters

Putative virus and plasmid contigs were identified by screening genomes with geNomad ^40^ using the end-to-end analysis parameter. Only hits greater than 5 kb were included in downstream analyses. Biosynthetic gene clusters were predicted using the JGI Secondary Metabolites Collaboratory pipeline which primarily uses antiSMASH v7.0 for prediction ^41^.

## DATA AVAILABILITY STATEMENT

Raw sequencing reads were deposited in NCBI’s SRA (https://www.ncbi.nlm.nih.gov/sra), and annotated assemblies of environmental SAGs based on 1M reads were deposited in the JGI’s Integrated Microbial Genomes and Microbiomes database (https://img.jgi.doe.gov/). Bioproject, biosample, and IMG genome ID’s can be found in Supplementary Tables S5 and S6.

## AUTHOR CONTRIBUTIONS

CG, RRM, TW, IB and VG-P designed the study. VGP, DG, and KW performed the experiments. RMB, MB, DWU, CG, IB and DK analyzed the data. CG, VGP, RRM, RMB, and TW wrote the manuscript. All authors read and approved the final manuscript.

## ACKNOWLEDGEMENTS

*B. subtilis* (pDR244) was kindly provided by Dr. Ilana Brito. The work conducted by the U.S. Department of Energy Joint Genome Institute (https://ror.org/04xm1d337), a DOE Office of Science User Facility, was supported by the Office of Science of the U.S. Department of Energy operated under Contract No. DE-AC02-05CH11231.

---

**Supplemental Figure 1.**
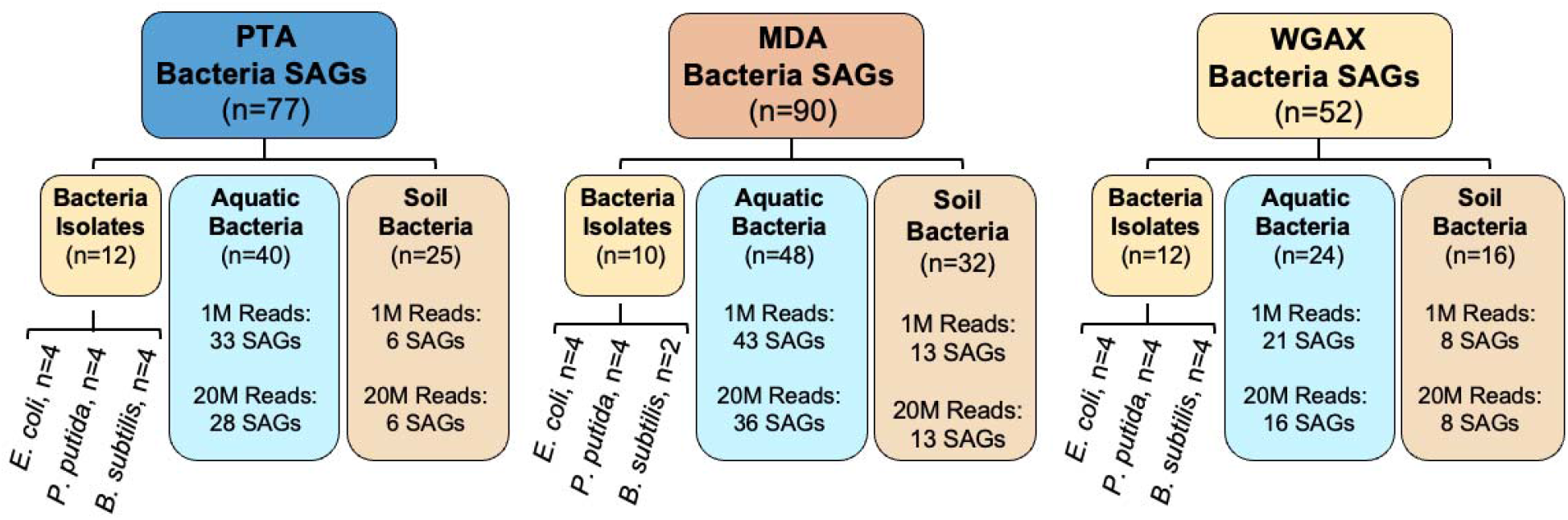
Number of SAGs amplified using each chemistry. The dataset consists of single cells derived from bacterial isolates and environmental samples from aquatic and soil samples. To generate *de novo* assemblies from low and high levels of sequencing effort, each library was first subsampled to a maximum of 1M and, where possible, 20M quality filtered reads.

**Supplementary Figure 2.**
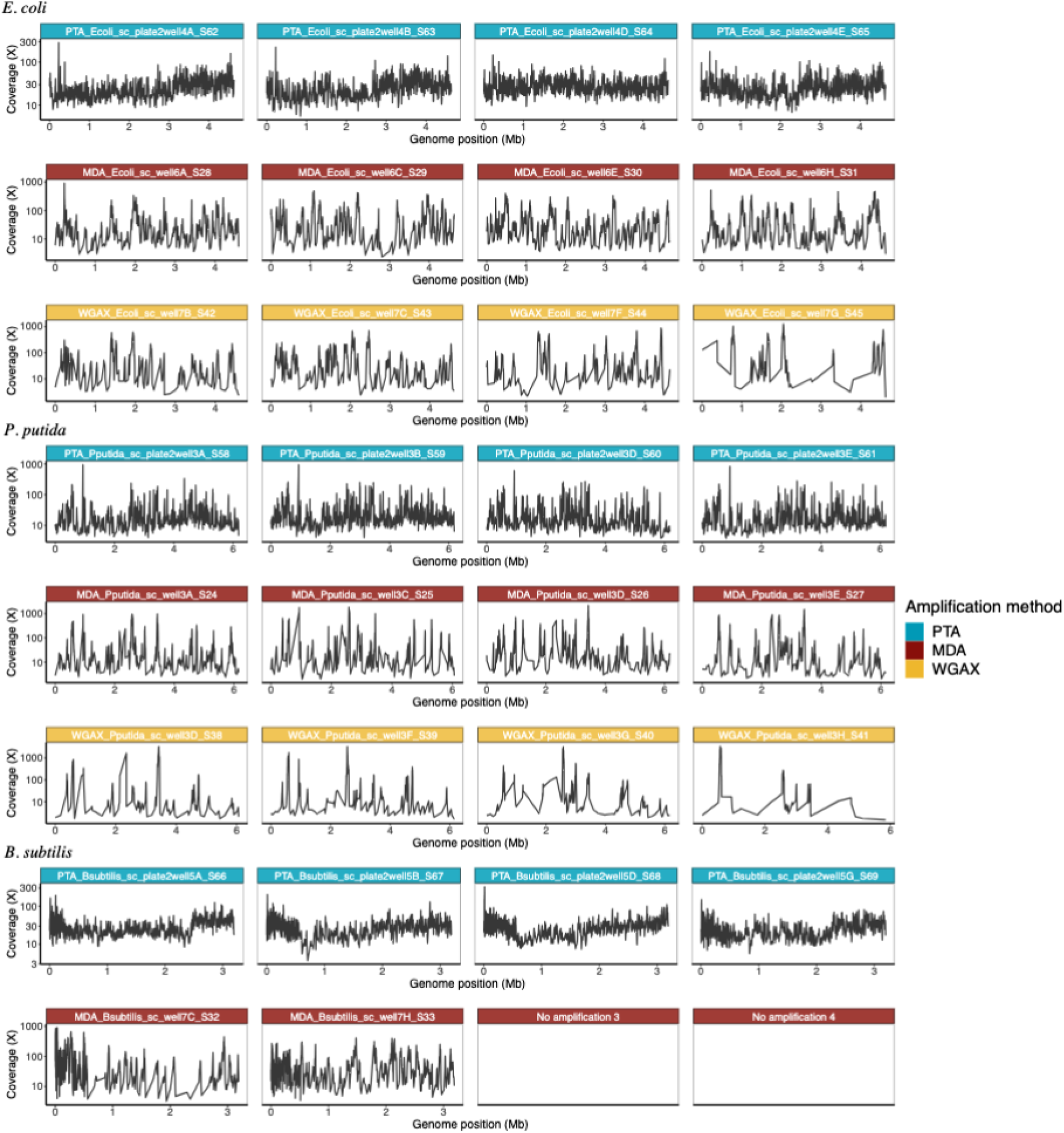
Genome coverage of 500 bp windows of all replicates from each species amplified with each chemistry. WGA-X amplification reactions of *B. subtilis* failed and were excluded, and only two MDA amplifications of *B. subtilis* were successful.

**Supplementary Figure 3.**
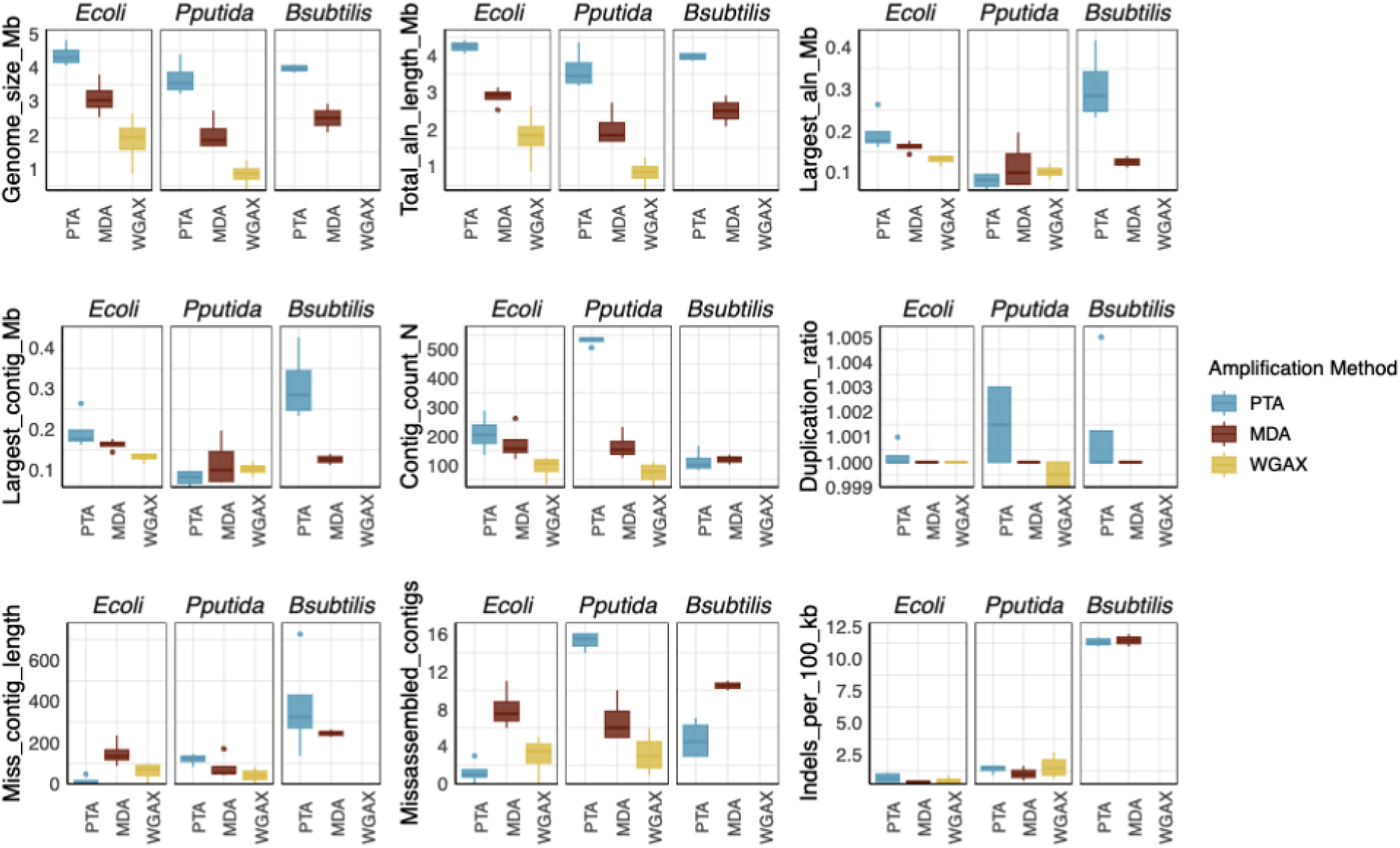
Additional genome quality parameters not present in Fig. 1 of isolate single-cell genomes. Boxplots display the minimum, 25th percentile, median, 75th percentile and maximum values. The dots represent outliers that are beyond 1.5 ^*^ interquartile range.

**Supplemental Figure 4.**
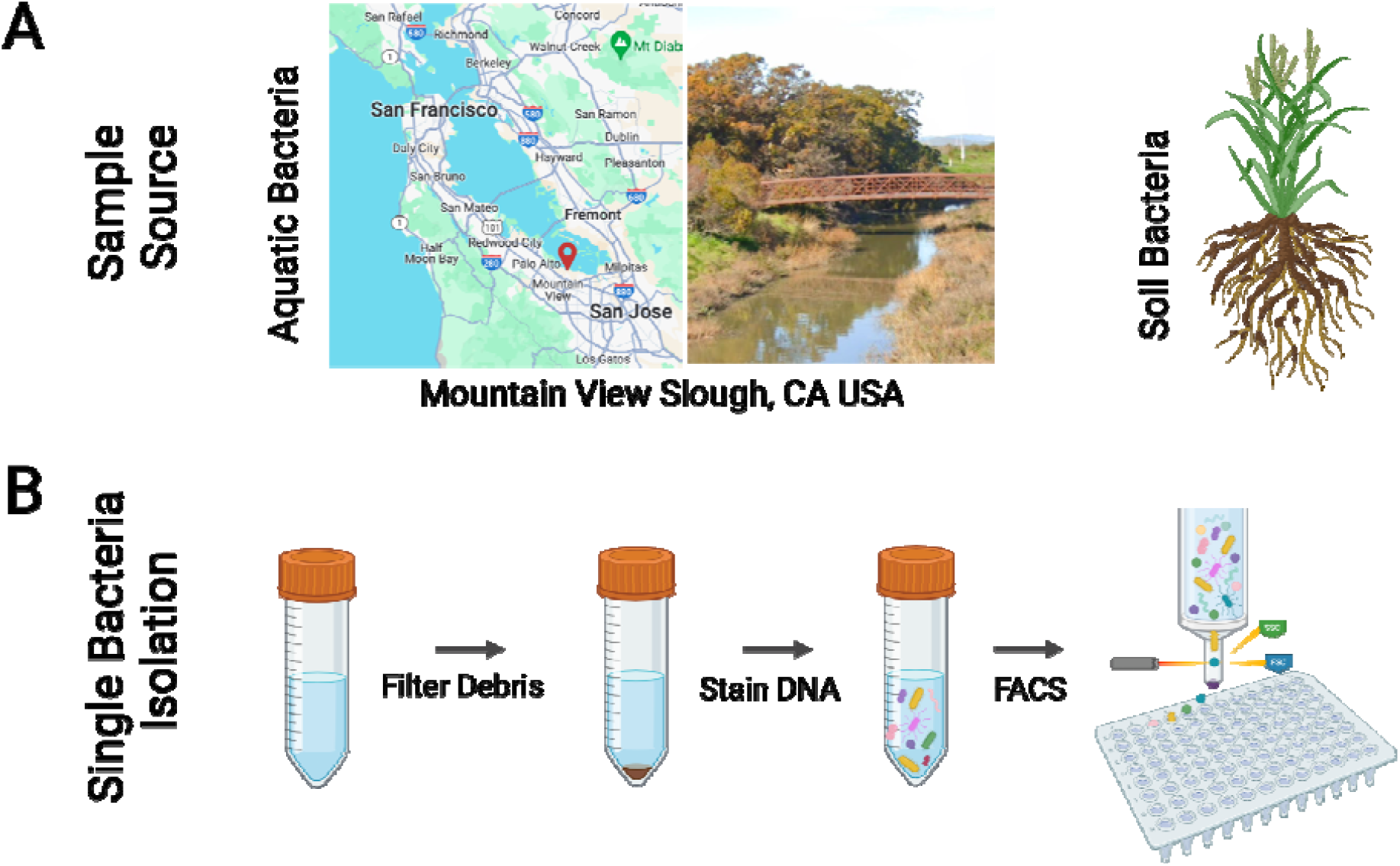
Sampling and processing of aquatic and soil samples. **A)** An aquatic sample was collected from the surface waters of Mountain View Slough and a soil microbiome sample was collected from the roots of a plant at Lawrence Berkeley National Laboratory. **B)** Samples were filtered to remove large particles and cells were stained with SYBR Green prior to FACS sorting of single bacteria.

**Supplementary Figure 5.**
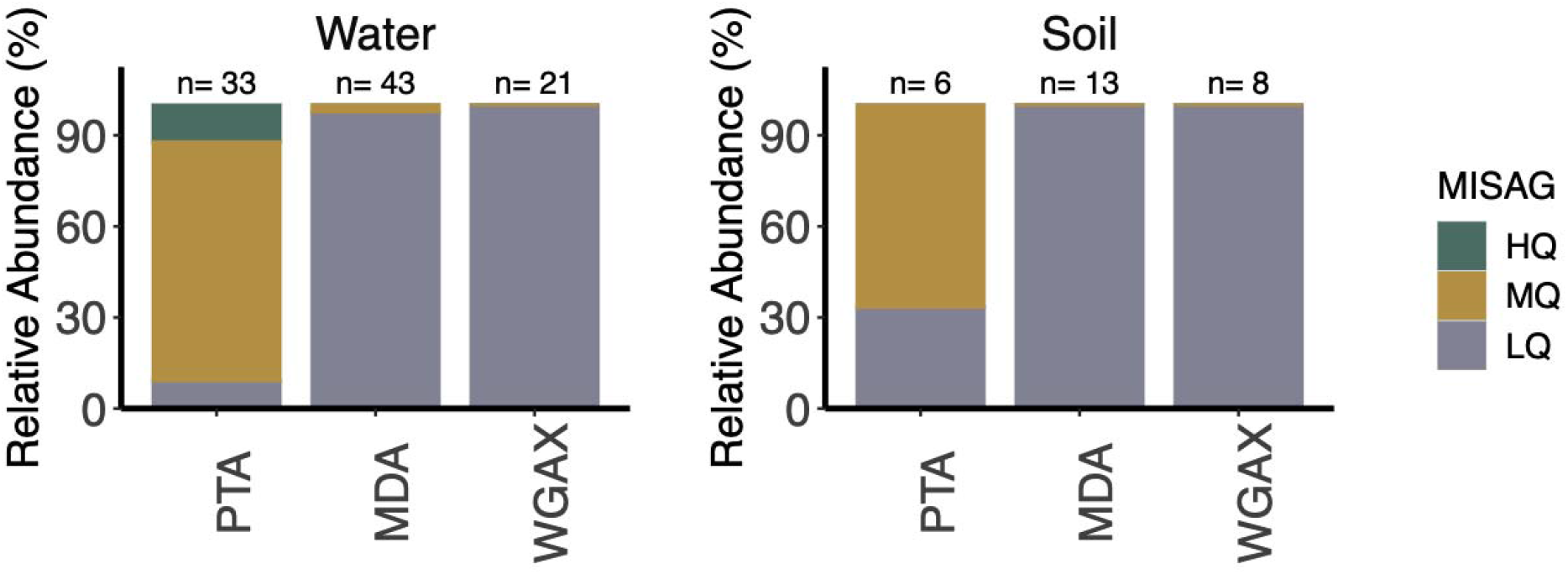
Environmental single cells from aquatic and soil samples categorized into the minimum information MISAG standards. The high-quality standard draft criterion includes a completeness score of > 90%, a contamination score of < 5%, the presence of the 23S, 16S and 5S rRNA genes and at least 18 tRNAs ^33^. The 4 genomes that made up the HQ fraction of the PTA aquatic samples satisfy these requirements, however the 16S rRNA genes were excluded from the final genomes as they were removed as a side-effect of the informatic decontamination procedure. This is a common problem when extracting MAGs from metagenomes ^42^, and for the same reasons, were removed after single cell decontamination likely due to variation in tetranucleotide frequencies.

