## Supplementary_Table_S1 for "scMicrobe PTA: Near Complete Genomes from Single Bacterial Cells"

**Supplementary Table S1. Summary statistics including mean, standard deviation and median for each of the measured genome statistics grouped by isolate species and amplification method.**

| sample_type | E. coli | E. coli | E. coli | P. putida | P. putida | P. putida | B. subtilis | B. subtilis |
|---|---|---|---|---|---|---|---|---|
| amp_method | PTA | MDA | WGAX | PTA | MDA | WGAX | PTA | MDA |
| replicate_numbers | 4 | 4 | 4 | 4 | 4 | 4 | 4 | 2 |
| Genome_fraction_mean | 91 | 62 | 39 | 58 | 33 | 13 | 94 | 60 |
| Genome_fraction_sd | 3.3 | 5.5 | 16 | 8.4 | 7.9 | 5.9 | 2.4 | 14 |
| Genome_fraction_median | 92 | 63 | 40 | 56 | 30 | 14 | 95 | 60 |
| Contig_N50_kb_mean | 41 | 45 | 44 | 9.2 | 28 | 31 | 90 | 45 |
| Contig_N50_kb_sd | 7.4 | 12 | 9.2 | 1.6 | 4.8 | 10 | 42 | 6.9 |
| Contig_N50_kb_median | 38 | 43 | 44 | 8.7 | 26 | 29 | 84 | 45 |
| MM_per_100_kb_mean | 3.2 | 3.3 | 3.9 | 11 | 11 | 12 | 6.1 | 4.8 |
| MM_per_100_kb_sd | 1.9 | 0.39 | 0.68 | 2.9 | 4.1 | 4.3 | 4 | 2.4 |
| MM_per_100_kb_median | 2.4 | 3.2 | 4.1 | 10 | 11 | 11 | 6.4 | 4.8 |
| gini_mean | 0.36 | 0.68 | 0.83 | 0.62 | 0.87 | 0.96 | 0.35 | 0.72 |
| gini_sd | 0.04 | 0.037 | 0.079 | 0.027 | 0.034 | 0.02 | 0.02 | 0.11 |
| gini_median | 0.37 | 0.68 | 0.83 | 0.62 | 0.88 | 0.96 | 0.35 | 0.72 |
| Genome_size_Mb_mean | 4.4 | 3.1 | 1.8 | 3.7 | 2 | 0.84 | 4 | 2.5 |
| Genome_size_Mb_sd | 0.33 | 0.53 | 0.74 | 0.5 | 0.49 | 0.36 | 0.096 | 0.59 |
| Genome_size_Mb_median | 4.3 | 3 | 1.9 | 3.5 | 1.8 | 0.87 | 4 | 2.5 |
| Total_aln_length_Mb_mean | 4.2 | 2.9 | 1.8 | 3.6 | 2 | 0.84 | 4 | 2.5 |
| Total_aln_length_Mb_sd | 0.15 | 0.26 | 0.73 | 0.53 | 0.49 | 0.36 | 0.094 | 0.59 |
| Total_aln_length_Mb_median | 4.2 | 2.9 | 1.9 | 3.5 | 1.9 | 0.87 | 4 | 2.5 |
| Largest_aln_Mb_mean | 0.19 | 0.16 | 0.13 | 0.08 | 0.12 | 0.1 | 0.31 | 0.13 |
| Largest_aln_Mb_sd | 0.046 | 0.013 | 0.012 | 0.019 | 0.059 | 0.016 | 0.083 | 0.019 |
| Largest_aln_Mb_median | 0.18 | 0.16 | 0.13 | 0.083 | 0.1 | 0.1 | 0.28 | 0.13 |
| Largest_contig_Mb_mean | 0.19 | 0.16 | 0.13 | 0.08 | 0.12 | 0.1 | 0.31 | 0.13 |
| Largest_contig_Mb_sd | 0.046 | 0.013 | 0.011 | 0.019 | 0.059 | 0.016 | 0.087 | 0.019 |
| Largest_contig_Mb_median | 0.18 | 0.16 | 0.13 | 0.083 | 0.1 | 0.1 | 0.28 | 0.13 |
| Contig_count_N_mean | 210 | 170 | 92 | 530 | 170 | 72 | 110 | 120 |
| Contig_count_N_sd | 64 | 61 | 41 | 16 | 47 | 39 | 37 | 24 |
| Contig_count_N_median | 200 | 160 | 100 | 540 | 150 | 77 | 100 | 120 |
| Duplication_ratio_mean | 1 | 1 | 1 | 1 | 1 | 1 | 1 | 1 |
| Duplication_ratio_sd | 5E-04 | 0 | 0 | 0.0017 | 0 | 0.00058 | 0.0025 | 0 |
| Duplication_ratio_median | 1 | 1 | 1 | 1 | 1 | 1 | 1 | 1 |
| Miss_contig_length_mean | 14 | 150 | 60 | 120 | 80 | 42 | 380 | 250 |
| Miss_contig_length_sd | 22 | 62 | 44 | 26 | 61 | 35 | 250 | 25 |
| Miss_contig_length_median | 4.3 | 130 | 70 | 130 | 53 | 42 | 320 | 250 |
| Missassembled_contigs_mean | 1.2 | 8 | 3 | 15 | 6.8 | 3.2 | 4.8 | 10 |
| Missassembled_contigs_sd | 1.3 | 2.2 | 2.2 | 0.96 | 2.4 | 2.2 | 2.1 | 0.71 |
| Missassembled_contigs_median | 1 | 7.5 | 3.5 | 16 | 6 | 3 | 4.5 | 10 |
| Indels_per_100_kb_mean | 0.7 | 0.41 | 0.48 | 1.4 | 1 | 1.6 | 11 | 11 |
| Indels_per_100_kb_sd | 0.38 | 0.069 | 0.29 | 0.32 | 0.49 | 0.92 | 0.32 | 0.7 |
| Indels_per_100_kb_median | 0.68 | 0.4 | 0.4 | 1.5 | 1 | 1.5 | 11 | 11 |
