## Supplementary_Table_S2 for "scMicrobe PTA: Near Complete Genomes from Single Bacterial Cells"

**Supplementary Table S2. One way ANOVA statistics comparing amplification methods on isolate single-cell genomes. All one-way ANOVAs yielding significant results underwent subsequent analysis using Tukey's Honestly Significant Difference (HSD) test to identify the specific pairwise comparisons contributing most significantly to the observed overall effect. Proportional metrics such as genome fraction and gini coefficient were arcsine transformed prior to analysis.**

| source | paper_source | method | parameter | comparison | meta_data | df | statistic | p.value |
|---|---|---|---|---|---|---|---|---|
| Isolate | fig1 | ANOVA | Contig_N50_kb | NA | Bsubtilis | 1 | 2.04 | 0.23 |
| Isolate | fig1 | ANOVA | Genome_fraction | NA | Bsubtilis | 1 | 37.25 | 0 |
| Isolate | fig1 | ANOVA | gini | NA | Bsubtilis | 1 | 45.52 | 0 |
| Isolate | fig1 | ANOVA | MM_per_100_kb | NA | Bsubtilis | 1 | 0.17 | 0.7 |
| Isolate | fig1 | ANOVA | Contig_N50_kb | NA | Ecoli | 2 | 0.2 | 0.82 |
| Isolate | fig1 | ANOVA | Genome_fraction | NA | Ecoli | 2 | 32.26 | 0 |
| Isolate | fig1 | ANOVA | gini | NA | Ecoli | 2 | 51.75 | 0 |
| Isolate | fig1 | ANOVA | MM_per_100_kb | NA | Ecoli | 2 | 0.45 | 0.65 |
| Isolate | fig1 | ANOVA | Contig_N50_kb | NA | Pputida | 2 | 13.01 | 0 |
| Isolate | fig1 | ANOVA | Genome_fraction | NA | Pputida | 2 | 32.81 | 0 |
| Isolate | fig1 | ANOVA | gini | NA | Pputida | 2 | 106.08 | 0 |
| Isolate | fig1 | ANOVA | MM_per_100_kb | NA | Pputida | 2 | 0.12 | 0.89 |
| Isolate | fig1 | Tukeys_HSD | Contig_N50_kb | PTA-MDA | Bsubtilis | NA | NA | 0.23 |
| Isolate | fig1 | Tukeys_HSD | Genome_fraction | PTA-MDA | Bsubtilis | NA | NA | 0 |
| Isolate | fig1 | Tukeys_HSD | gini | PTA-MDA | Bsubtilis | NA | NA | 0 |
| Isolate | fig1 | Tukeys_HSD | MM_per_100_kb | PTA-MDA | Bsubtilis | NA | NA | 0.7 |
| Isolate | fig1 | Tukeys_HSD | Contig_N50_kb | PTA-MDA | Ecoli | NA | NA | 0.82 |
| Isolate | fig1 | Tukeys_HSD | Contig_N50_kb | WGAX-MDA | Ecoli | NA | NA | 0.99 |
| Isolate | fig1 | Tukeys_HSD | Contig_N50_kb | WGAX-PTA | Ecoli | NA | NA | 0.89 |
| Isolate | fig1 | Tukeys_HSD | Genome_fraction | PTA-MDA | Ecoli | NA | NA | 0 |
| Isolate | fig1 | Tukeys_HSD | Genome_fraction | WGAX-MDA | Ecoli | NA | NA | 0.03 |
| Isolate | fig1 | Tukeys_HSD | Genome_fraction | WGAX-PTA | Ecoli | NA | NA | 0 |
| Isolate | fig1 | Tukeys_HSD | gini | PTA-MDA | Ecoli | NA | NA | 0 |
| Isolate | fig1 | Tukeys_HSD | gini | WGAX-MDA | Ecoli | NA | NA | 0.02 |
| Isolate | fig1 | Tukeys_HSD | gini | WGAX-PTA | Ecoli | NA | NA | 0 |
| Isolate | fig1 | Tukeys_HSD | MM_per_100_kb | PTA-MDA | Ecoli | NA | NA | 0.98 |
| Isolate | fig1 | Tukeys_HSD | MM_per_100_kb | WGAX-MDA | Ecoli | NA | NA | 0.77 |
| Isolate | fig1 | Tukeys_HSD | MM_per_100_kb | WGAX-PTA | Ecoli | NA | NA | 0.65 |
| Isolate | fig1 | Tukeys_HSD | Contig_N50_kb | PTA-MDA | Pputida | NA | NA | 0.01 |
| Isolate | fig1 | Tukeys_HSD | Contig_N50_kb | WGAX-MDA | Pputida | NA | NA | 0.78 |
| Isolate | fig1 | Tukeys_HSD | Contig_N50_kb | WGAX-PTA | Pputida | NA | NA | 0 |
| Isolate | fig1 | Tukeys_HSD | Genome_fraction | PTA-MDA | Pputida | NA | NA | 0.01 |
| Isolate | fig1 | Tukeys_HSD | Genome_fraction | WGAX-MDA | Pputida | NA | NA | 0.01 |
| Isolate | fig1 | Tukeys_HSD | Genome_fraction | WGAX-PTA | Pputida | NA | NA | 0 |
| Isolate | fig1 | Tukeys_HSD | gini | PTA-MDA | Pputida | NA | NA | 0 |
| Isolate | fig1 | Tukeys_HSD | gini | WGAX-MDA | Pputida | NA | NA | 0 |
| Isolate | fig1 | Tukeys_HSD | gini | WGAX-PTA | Pputida | NA | NA | 0 |
| Isolate | fig1 | Tukeys_HSD | MM_per_100_kb | PTA-MDA | Pputida | NA | NA | 1 |
| Isolate | fig1 | Tukeys_HSD | MM_per_100_kb | WGAX-MDA | Pputida | NA | NA | 0.9 |
| Isolate | fig1 | Tukeys_HSD | MM_per_100_kb | WGAX-PTA | Pputida | NA | NA | 0.93 |
| Isolate | supplemental | ANOVA | Contig_count_N | NA | Bsubtilis | 1 | 0.051 | 0.832 |
| Isolate | supplemental | ANOVA | Duplication_ratio | NA | Bsubtilis | 1 | 0.444 | 0.541 |
| Isolate | supplemental | ANOVA | Genome_size_Mb | NA | Bsubtilis | 1 | 30.297 | 0.005 |
| Isolate | supplemental | ANOVA | Indels_per_100_kb | NA | Bsubtilis | 1 | 0.089 | 0.781 |
| Isolate | supplemental | ANOVA | Largest_aln_Mb | NA | Bsubtilis | 1 | 8.16 | 0.046 |
| Isolate | supplemental | ANOVA | Largest_contig_Mb | NA | Bsubtilis | 1 | 7.557 | 0.051 |
| Isolate | supplemental | ANOVA | Miss_contig_length | NA | Bsubtilis | 1 | 0.506 | 0.516 |
| Isolate | supplemental | ANOVA | Missassembled_contigs | NA | Bsubtilis | 1 | 13.308 | 0.022 |
| Isolate | supplemental | ANOVA | Total_aln_length_Mb | NA | Bsubtilis | 1 | 30.442 | 0.005 |
| Isolate | supplemental | ANOVA | Contig_count_N | NA | Ecoli | 2 | 4.52 | 0.044 |
| Isolate | supplemental | ANOVA | Duplication_ratio | NA | Ecoli | 2 | 1 | 0.405 |
| Isolate | supplemental | ANOVA | Genome_size_Mb | NA | Ecoli | 2 | 20.585 | 0 |

**Supplementary Table S2. (Continued).**

| source | paper_source | method | parameter | comparison | meta_data | df | statistic | p.value |
|---|---|---|---|---|---|---|---|---|
| Isolate | supplemental | ANOVA | Missassembled_contigs | NA | Ecoli | 2 | 13.489 | 0.002 |
| Isolate | supplemental | ANOVA | Total_aln_length_Mb | NA | Ecoli | 2 | 28.963 | 0 |
| Isolate | supplemental | ANOVA | Contig_count_N | NA | Pputida | 2 | 175.877 | 0 |
| Isolate | supplemental | ANOVA | Duplication_ratio | NA | Pputida | 2 | 3.901 | 0.06 |
| Isolate | supplemental | ANOVA | Genome_size_Mb | NA | Pputida | 2 | 38.949 | 0 |
| Isolate | supplemental | ANOVA | Indels_per_100_kb | NA | Pputida | 2 | 0.774 | 0.49 |
| Isolate | supplemental | ANOVA | Largest_aln_Mb | NA | Pputida | 2 | 0.998 | 0.406 |
| Isolate | supplemental | ANOVA | Largest_contig_Mb | NA | Pputida | 2 | 0.998 | 0.406 |
| Isolate | supplemental | ANOVA | Miss_contig_length | NA | Pputida | 2 | 3.223 | 0.088 |
| Isolate | supplemental | ANOVA | Missassembled_contigs | NA | Pputida | 2 | 40.029 | 0 |
| Isolate | supplemental | ANOVA | Total_aln_length_Mb | NA | Pputida | 2 | 35.919 | 0 |
| Isolate | supplemental | Tukeys_HSD | Contig_count_N | MDA-PTA | Bsubtilis | NA | NA | 0.832 |
| Isolate | supplemental | Tukeys_HSD | Duplication_ratio | MDA-PTA | Bsubtilis | NA | NA | 0.541 |
| Isolate | supplemental | Tukeys_HSD | Genome_size_Mb | MDA-PTA | Bsubtilis | NA | NA | 0.005 |
| Isolate | supplemental | Tukeys_HSD | Indels_per_100_kb | MDA-PTA | Bsubtilis | NA | NA | 0.781 |
| Isolate | supplemental | Tukeys_HSD | Largest_aln_Mb | MDA-PTA | Bsubtilis | NA | NA | 0.046 |
| Isolate | supplemental | Tukeys_HSD | Largest_contig_Mb | MDA-PTA | Bsubtilis | NA | NA | 0.051 |
| Isolate | supplemental | Tukeys_HSD | Miss_contig_length | MDA-PTA | Bsubtilis | NA | NA | 0.516 |
| Isolate | supplemental | Tukeys_HSD | Missassembled_contigs | MDA-PTA | Bsubtilis | NA | NA | 0.022 |
| Isolate | supplemental | Tukeys_HSD | Total_aln_length_Mb | MDA-PTA | Bsubtilis | NA | NA | 0.005 |
| Isolate | supplemental | Tukeys_HSD | Contig_count_N | MDA-PTA | Ecoli | NA | NA | 0.678 |
| Isolate | supplemental | Tukeys_HSD | Contig_count_N | WGAX-PTA | Ecoli | NA | NA | 0.041 |
| Isolate | supplemental | Tukeys_HSD | Contig_count_N | WGAX-MDA | Ecoli | NA | NA | 0.152 |
| Isolate | supplemental | Tukeys_HSD | Duplication_ratio | MDA-PTA | Ecoli | NA | NA | 0.469 |
| Isolate | supplemental | Tukeys_HSD | Duplication_ratio | WGAX-PTA | Ecoli | NA | NA | 0.469 |
| Isolate | supplemental | Tukeys_HSD | Duplication_ratio | WGAX-MDA | Ecoli | NA | NA | 1 |
| Isolate | supplemental | Tukeys_HSD | Genome_size_Mb | MDA-PTA | Ecoli | NA | NA | 0.026 |
| Isolate | supplemental | Tukeys_HSD | Genome_size_Mb | WGAX-PTA | Ecoli | NA | NA | 0 |
| Isolate | supplemental | Tukeys_HSD | Genome_size_Mb | WGAX-MDA | Ecoli | NA | NA | 0.027 |
| Isolate | supplemental | Tukeys_HSD | Indels_per_100_kb | MDA-PTA | Ecoli | NA | NA | 0.345 |
| Isolate | supplemental | Tukeys_HSD | Indels_per_100_kb | WGAX-PTA | Ecoli | NA | NA | 0.517 |
| Isolate | supplemental | Tukeys_HSD | Indels_per_100_kb | WGAX-MDA | Ecoli | NA | NA | 0.938 |
| Isolate | supplemental | Tukeys_HSD | Largest_aln_Mb | MDA-PTA | Ecoli | NA | NA | 0.296 |
| Isolate | supplemental | Tukeys_HSD | Largest_aln_Mb | WGAX-PTA | Ecoli | NA | NA | 0.029 |
| Isolate | supplemental | Tukeys_HSD | Largest_aln_Mb | WGAX-MDA | Ecoli | NA | NA | 0.32 |
| Isolate | supplemental | Tukeys_HSD | Largest_contig_Mb | MDA-PTA | Ecoli | NA | NA | 0.3 |
| Isolate | supplemental | Tukeys_HSD | Largest_contig_Mb | WGAX-PTA | Ecoli | NA | NA | 0.03 |
| Isolate | supplemental | Tukeys_HSD | Largest_contig_Mb | WGAX-MDA | Ecoli | NA | NA | 0.318 |
| Isolate | supplemental | Tukeys_HSD | Miss_contig_length | MDA-PTA | Ecoli | NA | NA | 0.006 |
| Isolate | supplemental | Tukeys_HSD | Miss_contig_length | WGAX-PTA | Ecoli | NA | NA | 0.372 |
| Isolate | supplemental | Tukeys_HSD | Miss_contig_length | WGAX-MDA | Ecoli | NA | NA | 0.054 |
| Isolate | supplemental | Tukeys_HSD | Missassembled_contigs | MDA-PTA | Ecoli | NA | NA | 0.002 |
| Isolate | supplemental | Tukeys_HSD | Missassembled_contigs | WGAX-PTA | Ecoli | NA | NA | 0.431 |
| Isolate | supplemental | Tukeys_HSD | Missassembled_contigs | WGAX-MDA | Ecoli | NA | NA | 0.012 |
| Isolate | supplemental | Tukeys_HSD | Total_aln_length_Mb | MDA-PTA | Ecoli | NA | NA | 0.006 |
| Isolate | supplemental | Tukeys_HSD | Total_aln_length_Mb | WGAX-PTA | Ecoli | NA | NA | 0 |
| Isolate | supplemental | Tukeys_HSD | Total_aln_length_Mb | WGAX-MDA | Ecoli | NA | NA | 0.02 |
| Isolate | supplemental | Tukeys_HSD | Contig_count_N | MDA-PTA | Pputida | NA | NA | 0 |
| Isolate | supplemental | Tukeys_HSD | Contig_count_N | WGAX-PTA | Pputida | NA | NA | 0 |
| Isolate | supplemental | Tukeys_HSD | Contig_count_N | WGAX-MDA | Pputida | NA | NA | 0.014 |
| Isolate | supplemental | Tukeys_HSD | Duplication_ratio | MDA-PTA | Pputida | NA | NA | 0.165 |
| Isolate | supplemental | Tukeys_HSD | Duplication_ratio | WGAX-PTA | Pputida | NA | NA | 0.059 |
| Isolate | supplemental | Tukeys_HSD | Duplication_ratio | WGAX-MDA | Pputida | NA | NA | 0.785 |
| Isolate | supplemental | Tukeys_HSD | Genome_size_Mb | MDA-PTA | Pputida | NA | NA | 0.002 |
| Isolate | supplemental | Tukeys_HSD | Genome_size_Mb | WGAX-PTA | Pputida | NA | NA | 0 |
| Isolate | supplemental | Tukeys_HSD | Genome_size_Mb | WGAX-MDA | Pputida | NA | NA | 0.013 |
| Isolate | supplemental | Tukeys_HSD | Indels_per_100_kb | MDA-PTA | Pputida | NA | NA | 0.693 |

**Supplementary Table S2. (Continued).**

| source | paper_source | method | parameter | comparison | meta_data | df | statistic | p.value |
|---|---|---|---|---|---|---|---|---|
| Isolate | supplemental | Tukeys_HSD | Indels_per_100_kb | WGAX-PTA | Pputida | NA | NA | 0.923 |
| Isolate | supplemental | Tukeys_HSD | Indels_per_100_kb | WGAX-MDA | Pputida | NA | NA | 0.474 |
| Isolate | supplemental | Tukeys_HSD | Largest_aln_Mb | MDA-PTA | Pputida | NA | NA | 0.379 |
| Isolate | supplemental | Tukeys_HSD | Largest_aln_Mb | WGAX-PTA | Pputida | NA | NA | 0.687 |
| Isolate | supplemental | Tukeys_HSD | Largest_aln_Mb | WGAX-MDA | Pputida | NA | NA | 0.845 |
| Isolate | supplemental | Tukeys_HSD | Largest_contig_Mb | MDA-PTA | Pputida | NA | NA | 0.379 |
| Isolate | supplemental | Tukeys_HSD | Largest_contig_Mb | WGAX-PTA | Pputida | NA | NA | 0.687 |
| Isolate | supplemental | Tukeys_HSD | Largest_contig_Mb | WGAX-MDA | Pputida | NA | NA | 0.845 |
| Isolate | supplemental | Tukeys_HSD | Miss_contig_length | MDA-PTA | Pputida | NA | NA | 0.429 |
| Isolate | supplemental | Tukeys_HSD | Miss_contig_length | WGAX-PTA | Pputida | NA | NA | 0.074 |
| Isolate | supplemental | Tukeys_HSD | Miss_contig_length | WGAX-MDA | Pputida | NA | NA | 0.463 |
| Isolate | supplemental | Tukeys_HSD | Missassembled_contigs | MDA-PTA | Pputida | NA | NA | 0 |
| Isolate | supplemental | Tukeys_HSD | Missassembled_contigs | WGAX-PTA | Pputida | NA | NA | 0 |
| Isolate | supplemental | Tukeys_HSD | Missassembled_contigs | WGAX-MDA | Pputida | NA | NA | 0.074 |
| Isolate | supplemental | Tukeys_HSD | Total_aln_length_Mb | MDA-PTA | Pputida | NA | NA | 0.002 |
| Isolate | supplemental | Tukeys_HSD | Total_aln_length_Mb | WGAX-PTA | Pputida | NA | NA | 0 |
| Isolate | supplemental | Tukeys_HSD | Total_aln_length_Mb | WGAX-MDA | Pputida | NA | NA | 0.013 |
