## Supplementary_Table_S3 for "scMicrobe PTA: Near Complete Genomes from Single Bacterial Cells"

**Supplementary Table S3. Statistical analysis of environmental SAGs. One way ANOVA statistics to compare the effect of amplification method on environmental single cell genomes derived from water and soil samples. All one-way ANOVAs yielding significant results underwent subsequent analysis using Tukey's Honestly Significant Difference (HSD) test to identify the specific pairwise comparisons contributing most significantly to the observed overall effect. Proportional metrics such as genome completeness and contamination were arcsine transformed prior to analysis. The Fisher's exact test compared the effect of amplification method on the fraction of environmental single cell genomes with at least one virus (> 5 kb), plasmid (> 5 kb) or BGC, derived from water and soil samples.**

| source | paper_source | statistical_assum | method | parameter | comparison | meta_data | df | statistic | p.value |
|---|---|---|---|---|---|---|---|---|---|
| Environmental | fig2 | parametric | ANOVA | Completeness | NA | Soil | 2 | 19.59 | 0 |
| Environmental | fig2 | parametric | ANOVA | Contamination | NA | Soil | 2 | 13.21 | 0 |
| Environmental | fig2 | parametric | ANOVA | Contig_N50_kb | NA | Soil | 2 | 2.96 | 0.07 |
| Environmental | fig2 | parametric | ANOVA | Completeness | NA | Water | 2 | 246.31 | 0 |
| Environmental | fig2 | parametric | ANOVA | Contamination | NA | Water | 2 | 73.24 | 0 |
| Environmental | fig2 | parametric | ANOVA | Contig_N50_kb | NA | Water | 2 | 17.04 | 0 |
| Environmental | fig2 | parametric | Tukeys_HSD | Completeness | PTA-MDA | Soil | NA | NA | 0 |
| Environmental | fig2 | parametric | Tukeys_HSD | Completeness | WGAX-MDA | Soil | NA | NA | 0.73 |
| Environmental | fig2 | parametric | Tukeys_HSD | Completeness | WGAX-PTA | Soil | NA | NA | 0 |
| Environmental | fig2 | parametric | Tukeys_HSD | Contamination | PTA-MDA | Soil | NA | NA | 0 |
| Environmental | fig2 | parametric | Tukeys_HSD | Contamination | WGAX-MDA | Soil | NA | NA | 0.99 |
| Environmental | fig2 | parametric | Tukeys_HSD | Contamination | WGAX-PTA | Soil | NA | NA | 0 |
| Environmental | fig2 | parametric | Tukeys_HSD | Contig_N50_kb | PTA-MDA | Soil | NA | NA | 0.07 |
| Environmental | fig2 | parametric | Tukeys_HSD | Contig_N50_kb | WGAX-MDA | Soil | NA | NA | 0.96 |
| Environmental | fig2 | parametric | Tukeys_HSD | Contig_N50_kb | WGAX-PTA | Soil | NA | NA | 0.15 |
| Environmental | fig2 | parametric | Tukeys_HSD | Completeness | PTA-MDA | Water | NA | NA | 0 |
| Environmental | fig2 | parametric | Tukeys_HSD | Completeness | WGAX-MDA | Water | NA | NA | 0.14 |
| Environmental | fig2 | parametric | Tukeys_HSD | Completeness | WGAX-PTA | Water | NA | NA | 0 |
| Environmental | fig2 | parametric | Tukeys_HSD | Contamination | PTA-MDA | Water | NA | NA | 0 |
| Environmental | fig2 | parametric | Tukeys_HSD | Contamination | WGAX-MDA | Water | NA | NA | 0.75 |
| Environmental | fig2 | parametric | Tukeys_HSD | Contamination | WGAX-PTA | Water | NA | NA | 0 |
| Environmental | fig2 | parametric | Tukeys_HSD | Contig_N50_kb | PTA-MDA | Water | NA | NA | 0 |
| Environmental | fig2 | parametric | Tukeys_HSD | Contig_N50_kb | WGAX-MDA | Water | NA | NA | 0.96 |
| Environmental | fig2 | parametric | Tukeys_HSD | Contig_N50_kb | WGAX-PTA | Water | NA | NA | 0 |
| Environmental | supplemental | parametric | ANOVA | Contig count | NA | Water | 2 | 41.314 | 0 |
| Environmental | supplemental | parametric | ANOVA | GC (%) | NA | Water | 2 | 1.261 | 0.288 |
| Environmental | supplemental | parametric | ANOVA | Genome size Mb | NA | Water | 2 | 143.303 | 0 |
| Environmental | supplemental | parametric | ANOVA | Contig count | NA | Soil | 2 | 72.652 | 0 |
| Environmental | supplemental | parametric | ANOVA | GC (%) | NA | Soil | 2 | 0.002 | 0.998 |
| Environmental | supplemental | parametric | ANOVA | Genome size Mb | NA | Soil | 2 | 28.399 | 0 |
| Environmental | supplemental | parametric | Tukeys_HSD | Contig count | MDA-PTA | Water | NA | NA | 0 |
| Environmental | supplemental | parametric | Tukeys_HSD | Contig count | WGAX-PTA | Water | NA | NA | 0 |
| Environmental | supplemental | parametric | Tukeys_HSD | Contig count | WGAX-MDA | Water | NA | NA | 0.34 |
| Environmental | supplemental | parametric | Tukeys_HSD | GC (%) | MDA-PTA | Water | NA | NA | 0.98 |
| Environmental | supplemental | parametric | Tukeys_HSD | GC (%) | WGAX-PTA | Water | NA | NA | 0.39 |
| Environmental | supplemental | parametric | Tukeys_HSD | GC (%) | WGAX-MDA | Water | NA | NA | 0.28 |
| Environmental | supplemental | parametric | Tukeys_HSD | Genome size Mb | MDA-PTA | Water | NA | NA | 0 |
| Environmental | supplemental | parametric | Tukeys_HSD | Genome size Mb | WGAX-PTA | Water | NA | NA | 0 |
| Environmental | supplemental | parametric | Tukeys_HSD | Genome size Mb | WGAX-MDA | Water | NA | NA | 0.91 |
| Environmental | supplemental | parametric | Tukeys_HSD | Contig count | MDA-PTA | Soil | NA | NA | 0 |
| Environmental | supplemental | parametric | Tukeys_HSD | Contig count | WGAX-PTA | Soil | NA | NA | 0 |
| Environmental | supplemental | parametric | Tukeys_HSD | Contig count | WGAX-MDA | Soil | NA | NA | 0.72 |
| Environmental | supplemental | parametric | Tukeys_HSD | GC (%) | MDA-PTA | Soil | NA | NA | 1 |
| Environmental | supplemental | parametric | Tukeys_HSD | GC (%) | WGAX-PTA | Soil | NA | NA | 1 |
| Environmental | supplemental | parametric | Tukeys_HSD | GC (%) | WGAX-MDA | Soil | NA | NA | 1 |
| Environmental | supplemental | parametric | Tukeys_HSD | Genome size Mb | MDA-PTA | Soil | NA | NA | 0 |
| Environmental | supplemental | parametric | Tukeys_HSD | Genome size Mb | WGAX-PTA | Soil | NA | NA | 0 |
| Environmental | supplemental | parametric | Tukeys_HSD | Genome size Mb | WGAX-MDA | Soil | NA | NA | 0.73 |
| Environmental | fig2 | non_parametric | Fishers_exact | virus_contig_gtr5kb | PTA_vs_MDA | Water | NA | 23.425 | 0 |
| Environmental | fig2 | non_parametric | Fishers_exact | virus_contig_gtr5kb | PTA_vs_WGAX | Water | NA | 0.267 | 0.047 |
| Environmental | fig2 | non_parametric | Fishers_exact | virus_contig_gtr5kb | MDA_vs_WGAX | Water | NA | 6.196 | 0.034 |
| Environmental | fig2 | non_parametric | Fishers_exact | virus_contig_gtr5kb | PTA_vs_MDA | Soil | NA | 5.361 | 0.222 |
| Environmental | fig2 | non_parametric | Fishers_exact | virus_contig_gtr5kb | PTA_vs_WGAX | Soil | NA | 0 | 0.165 |
| Environmental | fig2 | non_parametric | Fishers_exact | virus_contig_gtr5kb | MDA_vs_WGAX | Soil | NA | 0 | 1 |
| Environmental | fig2 | non_parametric | Fishers_exact | plasmid_contig_gtr5kb | PTA_vs_MDA | Water | NA | 1.546 | 0.478 |
| Environmental | fig2 | non_parametric | Fishers_exact | plasmid_contig_gtr5kb | PTA_vs_WGAX | Water | NA | 0.382 | 0.151 |
| Environmental | fig2 | non_parametric | Fishers_exact | plasmid_contig_gtr5kb | MDA_vs_WGAX | Water | NA | 0.588 | 0.407 |

**Supplementary Table S3. (Continued).**

| source | paper_source | statistical_assum | method | parameter | comparison | meta_data | df | statistic | p.value |
|---|---|---|---|---|---|---|---|---|---|
| Environmental | fig2 | non_parametric | Fishers_exact | bgc_pa | MDA_vs_WGAX | Water | NA | 0.725 | 0.769 |
| Environmental | fig2 | non_parametric | Fishers_exact | bgc_pa | PTA_vs_MDA | Soil | NA | 2.231 | 0.628 |
| Environmental | fig2 | non_parametric | Fishers_exact | bgc_pa | PTA_vs_WGAX | Soil | NA | 0.091 | 0.091 |
| Environmental | fig2 | non_parametric | Fishers_exact | bgc_pa | MDA_vs_WGAX | Soil | NA | 0.181 | 0.174 |
