## Supplementary_Table_S4 for "scMicrobe PTA: Near Complete Genomes from Single Bacterial Cells"

**Supplementary Table S4. Summary statistics including mean, standard deviation and median for each of the measured genome statistics grouped by sample environment and amplification method.**

| subsampling | 1M | 1M | 1M | 1M | 1M | 1M | 20M | 20M | 20M | 20M | 20M | 20M |
|---|---|---|---|---|---|---|---|---|---|---|---|---|
| system | Water | Water | Water | Soil | Soil | Soil | Water | Water | Water | Soil | Soil | Soil |
| amp_method | PTA | MDA | WGAX | PTA | MDA | WGAX | PTA | MDA | WGAX | PTA | MDA | WGAX |
| replicate_numbers | 33 | 43 | 21 | 6 | 13 | 8 | 28 | 36 | 16 | 6 | 13 | 8 |
| Completeness_mean | 79 | 19 | 14 | 48 | 13 | 9 | 87 | 32 | 25 | 79 | 22 | 16 |
| Completeness_sd | 16 | 10 | 6.7 | 18 | 12 | 5.1 | 12 | 14 | 14 | 11 | 17 | 11 |
| Completeness_median | 83 | 17 | 11 | 57 | 8 | 8 | 89 | 30 | 23 | 83 | 15 | 15 |
| Contamination_mean | 1.6 | 0.039 | 0.0067 | 1.4 | 0.08 | 0.046 | 6.8 | 0.41 | 0.12 | 7 | 0.57 | 0.06 |
| Contamination_sd | 1.5 | 0.092 | 0.014 | 1.2 | 0.16 | 0.084 | 2.6 | 0.62 | 0.22 | 3.7 | 1.4 | 0.079 |
| Contamination_median | 1.1 | 0.01 | 0 | 1.6 | 0.01 | 0.02 | 6.7 | 0.18 | 0.03 | 8.9 | 0.08 | 0.035 |
| Contig_N50_kb_mean | 54 | 13 | 15 | 11 | 26 | 24 | 190 | 13 | 14 | 24 | 25 | 25 |
| Contig_N50_kb_sd | 55 | 5.6 | 6 | 5 | 14 | 14 | 550 | 4.9 | 6.4 | 10 | 17 | 18 |
| Contig_N50_kb_median | 34 | 12 | 15 | 11 | 26 | 23 | 48 | 12 | 13 | 25 | 18 | 19 |
| Contig.count_mean | 120 | 52 | 38 | 330 | 44 | 26 | 160 | 81 | 68 | 400 | 75 | 47 |
| Contig.count_sd | 58 | 18 | 16 | 100 | 28 | 24 | 49 | 23 | 24 | 110 | 41 | 31 |
| Contig.count_median | 120 | 54 | 35 | 310 | 38 | 20 | 160 | 83 | 69 | 360 | 81 | 41 |
| Genome.size.Mb_mean | 1.9 | 0.45 | 0.41 | 2.4 | 0.52 | 0.33 | 2.2 | 0.69 | 0.71 | 4.6 | 1 | 0.66 |
| Genome.size.Mb_sd | 0.61 | 0.18 | 0.25 | 0.96 | 0.44 | 0.31 | 0.67 | 0.2 | 0.41 | 1.2 | 0.78 | 0.57 |
| Genome.size.Mb_median | 1.9 | 0.42 | 0.34 | 2.3 | 0.37 | 0.23 | 2.2 | 0.66 | 0.68 | 4.6 | 0.98 | 0.45 |
| GC_mean | 0.45 | 0.44 | 0.48 | 0.57 | 0.58 | 0.58 | 0.44 | 0.43 | 0.47 | 0.58 | 0.58 | 0.58 |
| GC_sd | 0.095 | 0.079 | 0.047 | 0.11 | 0.097 | 0.12 | 0.086 | 0.079 | 0.049 | 0.11 | 0.098 | 0.12 |
| GC_median | 0.44 | 0.45 | 0.48 | 0.6 | 0.63 | 0.63 | 0.42 | 0.43 | 0.46 | 0.62 | 0.64 | 0.64 |
