## Supplementary_Table_S5 for "scMicrobe PTA: Near Complete Genomes from Single Bacterial Cells"

**Supplementary Table S5: Metadata for individual environmental SAG sequencing and overview of de novo assembly results.**

| genome_id | subsampling | seq_platform | BioProject | BioSample | IMG Genome ID | amp_method | system | library_name | gtdb_taxonomy | bgc_pa | Enzymatic Lysis | misag | rrna_16S_hit |
|---|---|---|---|---|---|---|---|---|---|---|---|---|---|
| scBac_PTA_EnviBac_sc_plate3well3A_S9 | 1M | novseq 2000 | PRJNA1067728 | SAMN39550262 | 3300071324 | PTA | Water | not_assigned | p__Bacteroidota;c__Bacteroidia;o__Flavobacteriales;f__Schleiferiaceae | present | TRUE | HQ | absent |
| scBac_PTA_EnviBac_sc_plate3well3B_S10 | 1M | novseq 2000 | PRJNA1067728 | SAMN39550263 | 3300071323 | PTA | Water | not_assigned | p__Bacteroidota;c__Bacteroidia;o__Flavobacteriales;f__Schleiferiaceae | present | TRUE | MQ | absent |
| scBac_PTA_EnviBac_sc_plate3well3C_S11 | 1M | novseq 2000 | PRJNA1067728 | SAMN39550264 | 3300071322 | PTA | Water | not_assigned | p__Proteobacteria;c__Gammaproteobacteria;o__Pseudomonadales;f__Nitrincolaceae | present | TRUE | MQ | absent |
| scBac_PTA_EnviBac_sc_plate3well3D_S12 | 1M | novseq 2000 | PRJNA1067728 | SAMN39550265 | 3300071321 | PTA | Water | not_assigned | p__Actinobacteriota;c__Acidimicrobiia;o__Acidimicrobiales;f__UBA11606 | present | TRUE | LQ | present |
| scBac_PTA_EnviBac_sc_plate3well3E_S13 | 1M | novseq 2000 | PRJNA1067728 | SAMN39550266 | 3300071320 | PTA | Water | not_assigned | p__Proteobacteria;c__Alphaproteobacteria;o__Rhodobacterales;f__Rhodobacteraceae | present | TRUE | MQ | absent |
| scBac_WGAX_EnviBac_sc_well3C_S95 | 1M | novseq 2000 | PRJNA1067728 | SAMN39550297 | 3300071319 | WGAX | Water | not_assigned | p__Proteobacteria;c__Gammaproteobacteria;o__Pseudomonadales;f__Halieaceae | present | FALSE | LQ | absent |
| scBac_WGAX_EnviBac_sc_well4H_S96 | 1M | novseq 2000 | PRJNA1067728 | SAMN39550298 | 3300071318 | WGAX | Water | not_assigned | unassigned | absent | FALSE | LQ | absent |
| scBac_WGAX_EnviBac_sc_well5B_S97 | 1M | novseq 2000 | PRJNA1067728 | SAMN39550299 | 3300071317 | WGAX | Water | not_assigned | p__Proteobacteria;c__Gammaproteobacteria;o__Pseudomonadales;f__Nitrincolaceae | absent | FALSE | LQ | absent |
| scBac_WGAX_EnviBac_sc_well6D_S99 | 1M | novseq 2000 | PRJNA1067728 | SAMN39550301 | 3300071316 | WGAX | Water | not_assigned | p__Proteobacteria;c__Gammaproteobacteria;o__Pseudomonadales;f__Halieaceae | present | FALSE | LQ | present |
| scBac_WGAX_EnviBac_sc_well7D_S100 | 1M | novseq 2000 | PRJNA1067728 | SAMN39550302 | 3300071315 | WGAX | Water | not_assigned | unassigned | absent | FALSE | LQ | absent |
| scBac_Watchmaker_EnviBac_sc_well3B_S79 | 1M | novseq 2000 | PRJNA1067728 | SAMN39550273 | 3300071314 | MDA | Water | not_assigned | p__Proteobacteria;c__Gammaproteobacteria;o__Pseudomonadales;f__Nitrincolaceae | present | FALSE | LQ | present |
| scBac_Watchmaker_EnviBac_sc_well3C_S80 | 1M | novseq 2000 | PRJNA1067728 | SAMN39550274 | 3300071313 | MDA | Water | not_assigned | p__Proteobacteria;c__Gammaproteobacteria;o__Pseudomonadales;f__Nitrincolaceae | absent | FALSE | LQ | present |
| scBac_Watchmaker_EnviBac_sc_well4C_S81 | 1M | novseq 2000 | PRJNA1067728 | SAMN39550275 | 3300071312 | MDA | Water | not_assigned | unassigned | absent | FALSE | LQ | present |
| scBac_Watchmaker_EnviBac_sc_well5A_S83 | 1M | novseq 2000 | PRJNA1067728 | SAMN39550277 | 3300071311 | MDA | Water | not_assigned | p__Proteobacteria;c__Alphaproteobacteria;o__Rhodobacterales;f__Rhodobacteraceae | absent | FALSE | LQ | absent |
| scBac_Watchmaker_EnviBac_sc_well5E_S84 | 1M | novseq 2000 | PRJNA1067728 | SAMN39550278 | 3300071310 | MDA | Water | not_assigned | p__Proteobacteria;c__Alphaproteobacteria;o__Rhodobacterales;f__Rhodobacteraceae | absent | FALSE | LQ | absent |
| scBac_Watchmaker_EnviBac_sc_well6E_S85 | 1M | novseq 2000 | PRJNA1067728 | SAMN39550279 | 3300071309 | MDA | Water | not_assigned | p__Proteobacteria;c__Alphaproteobacteria;o__Rhodobacterales;f__Rhodobacteraceae | absent | FALSE | LQ | present |
| scBac_Watchmaker_EnviBac_sc_well7E_S86 | 1M | novseq 2000 | PRJNA1067728 | SAMN39550280 | 3300071308 | MDA | Water | not_assigned | p__Bacteroidota;c__Bacteroidia;o__Flavobacteriales;f__Schleiferiaceae | absent | FALSE | LQ | present |
| EnvBac_Water_PTA_D06_scBac_plate.B | 1M | novseq 6000 | PRJNA1052789 | SAMN38845241 | 3300071202 | PTA | Water | OAHZZ | p__Proteobacteria;c__Gammaproteobacteria;o__Pseudomonadales;f__Nitrincolaceae | present | TRUE | MQ | absent |
| EnvBac_Water_WGAX_E12_scBac_plate.B | 1M | novseq 6000 | PRJNA1052790 | SAMN38845225 | 3300071203 | WGAX | Water | OANAA | p__Bacteroidota;c__Bacteroidia;o__Flavobacteriales;f__Schleiferiaceae | present | FALSE | LQ | absent |
| EnvBac_Water_PTA_A03_scBac_plate.B | 1M | novseq 6000 | PRJNA1052791 | SAMN38845175 | 3300071204 | PTA | Water | OANAB | p__Proteobacteria;c__Gammaproteobacteria;o__Pseudomonadales;f__HTCC2089 | present | TRUE | MQ | absent |
| EnvBac_Water_PTA_B03_scBac_plate.B | 1M | novseq 6000 | PRJNA1052792 | SAMN38845155 | 3300071245 | PTA | Water | OANAC | p__Bacteroidota;c__Bacteroidia;o__Flavobacteriales;f__Flavobacteriaceae | present | TRUE | MQ | absent |
| EnvBac_Water_PTA_C03_scBac_plate.B | 1M | novseq 6000 | PRJNA1052793 | SAMN38845156 | 3300071205 | PTA | Water | OANAG | p__Actinobacteriota;c__Actinomycetia;o__Actinomycetales;f__Microbacteriaceae | present | TRUE | MQ | absent |
| EnvBac_Water_PTA_D03_scBac_plate.B | 1M | novseq 6000 | PRJNA1052794 | SAMN38845251 | 3300071206 | PTA | Water | OANAH | p__Proteobacteria;c__Gammaproteobacteria;o__Enterobacterales;f__Alteromonadaceae | present | TRUE | MQ | absent |
| EnvBac_Water_PTA_E03_scBac_plate.B | 1M | novseq 6000 | PRJNA1052795 | SAMN38845210 | 3300071246 | PTA | Water | OANAN | p__Actinobacteriota;c__Acidimicrobiia;o__Acidimicrobiales;f__UBA11606 | absent | TRUE | LQ | absent |
| EnvBac_Water_PTA_F03_scBac_plate.B | 1M | novseq 6000 | PRJNA1052796 | SAMN38845157 | 3300071247 | PTA | Water | OANAO | p__Proteobacteria;c__Alphaproteobacteria;o__Pelagibacterales;f__Pelagibacteraceae | present | TRUE | HQ | absent |
| EnvBac_Water_PTA_G03_scBac_plate.B | 1M | novseq 6000 | PRJNA1052797 | SAMN38845159 | 3300071207 | PTA | Water | OANAP | p__Proteobacteria;c__Gammaproteobacteria;o__UBA4575;f__UBA4575 | present | TRUE | MQ | absent |
| EnvBac_Water_PTA_H03_scBac_plate.B | 1M | novseq 6000 | PRJNA1052798 | SAMN38845243 | 3300071282 | PTA | Water | OANAS | p__Proteobacteria;c__Gammaproteobacteria;o__Burkholderiales;f__Methylophilaceae | present | TRUE | MQ | absent |
| EnvBac_Water_PTA_A04_scBac_plate.B | 1M | novseq 6000 | PRJNA1052799 | SAMN38845163 | 3300071248 | PTA | Water | OANAT | p__Proteobacteria;c__Alphaproteobacteria;o__Pelagibacterales;f__Pelagibacteraceae | present | TRUE | MQ | absent |
| EnvBac_Water_PTA_B04_scBac_plate.B | 1M | novseq 6000 | PRJNA1052800 | SAMN38845160 | 3300071208 | PTA | Water | OANAU | p__Proteobacteria;c__Alphaproteobacteria;o__HIMB59;f__HIMB59 | present | TRUE | MQ | absent |
| EnvBac_Water_PTA_C04_scBac_plate.B | 1M | novseq 6000 | PRJNA1052801 | SAMN38845194 | 3300071249 | PTA | Water | OANAW | p__Proteobacteria;c__Gammaproteobacteria;o__UBA4575;f__UBA4575 | absent | TRUE | LQ | absent |
| EnvBac_Water_PTA_D04_scBac_plate.B | 1M | novseq 6000 | PRJNA1052802 | SAMN38845211 | 3300071209 | PTA | Water | OANAX | p__Bacteroidota;c__Bacteroidia;o__Flavobacteriales;f__Flavobacteriaceae | present | TRUE | MQ | absent |
| EnvBac_Water_PTA_E04_scBac_plate.B | 1M | novseq 6000 | PRJNA1052803 | SAMN38845212 | 3300071210 | PTA | Water | OANAY | p__Bacteroidota;c__Bacteroidia;o__Flavobacteriales;f__Flavobacteriaceae | present | TRUE | MQ | absent |
| EnvBac_Water_PTA_F04_scBac_plate.B | 1M | novseq 6000 | PRJNA1052804 | SAMN38845178 | 3300071250 | PTA | Water | OANAZ | p__Proteobacteria;c__Alphaproteobacteria;o__Pelagibacterales;f__Pelagibacteraceae | absent | TRUE | MQ | absent |
| EnvBac_Water_PTA_G04_scBac_plate.B | 1M | novseq 6000 | PRJNA1052805 | SAMN38845179 | 3300071251 | PTA | Water | OANBA | p__Proteobacteria;c__Gammaproteobacteria;o__Chromatiales;f__Sedimenticolaceae | absent | TRUE | MQ | absent |
| EnvBac_Water_PTA_H04_scBac_plate.B | 1M | novseq 6000 | PRJNA1052806 | SAMN38845176 | 3300071211 | PTA | Water | OANBB | p__Proteobacteria;c__Gammaproteobacteria;o__Burkholderiales;f__Methylophilaceae | present | TRUE | HQ | absent |
| EnvBac_Water_PTA_A05_scBac_plate.B | 1M | novseq 6000 | PRJNA1052807 | SAMN38845215 | 3300071212 | PTA | Water | OANBC | p__Actinobacteriota;c__Actinomycetia;o__Actinomycetales;f__Microbacteriaceae | present | TRUE | MQ | absent |
| EnvBac_Water_PTA_B05_scBac_plate.B | 1M | novseq 6000 | PRJNA1052808 | SAMN38845205 | 3300071213 | PTA | Water | OANBG | p__Proteobacteria;c__Gammaproteobacteria;o__Pseudomonadales;f__Halieaceae | present | TRUE | MQ | absent |
| EnvBac_Water_PTA_C05_scBac_plate.B | 1M | novseq 6000 | PRJNA1052809 | SAMN38845237 | 3300071214 | PTA | Water | OANBH | p__Proteobacteria;c__Alphaproteobacteria;o__Rhodobacterales;f__Rhodobacteraceae | present | TRUE | MQ | absent |
| EnvBac_Water_PTA_D05_scBac_plate.B | 1M | novseq 6000 | PRJNA1052810 | SAMN38845247 | 3300071252 | PTA | Water | OANBN | p__Proteobacteria;c__Gammaproteobacteria;o__Enterobacterales;f__Alteromonadaceae | present | TRUE | MQ | absent |
| EnvBac_Water_PTA_E05_scBac_plate.B | 1M | novseq 6000 | PRJNA1052811 | SAMN38845242 | 3300071215 | PTA | Water | OANBO | p__Proteobacteria;c__Alphaproteobacteria;o__Rhodobacterales;f__Rhodobacteraceae | present | TRUE | MQ | absent |
| EnvBac_Water_PTA_F05_scBac_plate.B | 1M | novseq 6000 | PRJNA1052812 | SAMN38845168 | 3300071253 | PTA | Water | OANBP | p__Bacteroidota;c__Bacteroidia;o__Flavobacteriales;f__Schleiferiaceae | present | TRUE | HQ | absent |
| EnvBac_Water_PTA_G05_scBac_plate.B | 1M | novseq 6000 | PRJNA1052813 | SAMN38845219 | 3300071216 | PTA | Water | OANBS | p__Proteobacteria;c__Alphaproteobacteria;o__Rhodobacterales;f__Rhodobacteraceae | present | TRUE | MQ | absent |
| EnvBac_Water_PTA_H05_scBac_plate.B | 1M | novseq 6000 | PRJNA1052814 | SAMN38845221 | 3300071217 | PTA | Water | OANBT | p__Proteobacteria;c__Alphaproteobacteria;o__Rickettsiales;f__ | present | TRUE | MQ | absent |
| EnvBac_Water_PTA_A06_scBac_plate.B | 1M | novseq 6000 | PRJNA1052815 | SAMN38845170 | 3300071254 | PTA | Water | OANBU | p__Bacteroidota;c__Bacteroidia;o__Flavobacteriales;f__Schleiferiaceae | present | TRUE | MQ | absent |
| EnvBac_Water_PTA_B06_scBac_plate.B | 1M | novseq 6000 | PRJNA1052816 | SAMN38845224 | 3300071283 | PTA | Water | OANBW | p__Proteobacteria;c__Alphaproteobacteria;o__Pelagibacterales;f__Pelagibacteraceae | present | TRUE | MQ | absent |
| EnvBac_Water_PTA_C06_scBac_plate.B | 1M | novseq 6000 | PRJNA1052817 | SAMN38845140 | 3300071218 | PTA | Water | OANBX | p__Bacteroidota;c__Rhodothermia;o__Balneolales;f__Balneolaceae | present | TRUE | MQ | absent |
| EnvBac_Water_Wchmk_A09_scBac_plate.B | 1M | novseq 6000 | PRJNA1052818 | SAMN38845172 | 3300071274 | MDA | Water | OANCC | p__Proteobacteria;c__Alphaproteobacteria;o__Pelagibacterales;f__Pelagibacteraceae | absent | FALSE | LQ | absent |
| EnvBac_Water_Wchmk_B09_scBac_plate.B | 1M | novseq 6000 | PRJNA1052819 | SAMN38845152 | 3300071284 | MDA | Water | OANCG | p__Bacteroidota;c__Bacteroidia;o__Flavobacteriales;f__Flavobacteriaceae | absent | FALSE | LQ | absent |
| EnvBac_Water_Wchmk_C09_scBac_plate.B | 1M | novseq 6000 | PRJNA1052820 | SAMN38845238 | 3300071255 | MDA | Water | OANCH | unassigned | absent | FALSE | LQ | absent |
| EnvBac_Water_Wchmk_D09_scBac_plate.B | 1M | novseq 6000 | PRJNA1052821 | SAMN38845249 | 3300071219 | MDA | Water | OANCN | p__Proteobacteria;c__Gammaproteobacteria;o__PS1;f__Thioglobaceae | absent | FALSE | LQ | absent |
| EnvBac_Water_Wchmk_E09_scBac_plate.B | 1M | novseq 6000 | PRJNA1052852 | SAMN38845174 | 3300071285 | MDA | Water | OANCO | p__Bacteroidota;c__Bacteroidia;o__Flavobacteriales;f__Flavobacteriaceae | present | FALSE | LQ | absent |
| EnvBac_Water_Wchmk_F09_scBac_plate.B | 1M | novseq 6000 | PRJNA1052853 | SAMN38845190 | 3300071286 | MDA | Water | OANCP | unassigned | absent | FALSE | LQ | absent |
| EnvBac_Water_Wchmk_G09_scBac_plate.B | 1M | novseq 6000 | PRJNA1052854 | SAMN38845161 | 3300071287 | MDA | Water | OANCS | p__Bacteroidota;c__Bacteroidia;o__Flavobacteriales;f__Flavobacteriaceae | present | FALSE | LQ | absent |
| EnvBac_Water_Wchmk_H09_scBac_plate.B | 1M | novseq 6000 | PRJNA1052855 | SAMN38845162 | 3300071220 | MDA | Water | OANCT | p__Bacteroidota;c__Bacteroidia;o__Flavobacteriales;f__Schleiferiaceae | absent | FALSE | LQ | absent |
| EnvBac_Water_Wchmk_A10_scBac_plate.B | 1M | novseq 6000 | PRJNA1052856 | SAMN38845180 | 3300071256 | MDA | Water | OANCU | p__Actinobacteriota;c__Actinomycetia;o__Actinomycetales;f__Microbacteriaceae | absent | FALSE | LQ | absent |
| EnvBac_Water_Wchmk_B10_scBac_plate.B | 1M | novseq 6000 | PRJNA1052857 | SAMN38845192 | 3300071221 | MDA | Water | OANCW | p__Bacteroidota;c__Bacteroidia;o__Flavobacteriales;f__Flavobacteriaceae | absent | FALSE | LQ | absent |
| EnvBac_Water_Wchmk_C10_scBac_plate.B | 1M | novseq 6000 | PRJNA1052858 | SAMN38845181 | 3300071257 | MDA | Water | OANCX | p__Proteobacteria;c__Alphaproteobacteria;o__Rhodobacterales;f__Rhodobacteraceae | present | FALSE | LQ | absent |
| EnvBac_Water_Wchmk_D10_scBac_plate.B | 1M | novseq 6000 | PRJNA1052859 | SAMN38845186 | 3300071222 | MDA | Water | OANCY | p__Bacteroidota;c__Bacteroidia;o__Flavobacteriales;f__Flavobacteriaceae | present | FALSE | LQ | absent |
| EnvBac_Water_Wchmk_E10_scBac_plate.B | 1M | novseq 6000 | PRJNA1052860 | SAMN38845234 | 3300071258 | MDA | Water | OANCZ | p__Proteobacteria;c__Alphaproteobacteria;o__Rhodobacterales;f__Rhodobacteraceae | absent | FALSE | LQ | absent |
| EnvBac_Water_Wchmk_F10_scBac_plate.B | 1M | novseq 6000 | PRJNA1052861 | SAMN38845166 | 3300071259 | MDA | Water | OANGA | p__Proteobacteria;c__Gammaproteobacteria;o__Chromatiales;f__Sedimenticolaceae | absent | FALSE | LQ | absent |
| EnvBac_Water_Wchmk_G10_scBac_plate.B | 1M | novseq 6000 | PRJNA1052862 | SAMN38845187 | 3300071223 | MDA | Water | OANGB | unassigned | absent | FALSE | LQ | absent |
| EnvBac_Water_Wchmk_H10_scBac_plate.B | 1M | novseq 6000 | PRJNA1052863 | SAMN38845223 | 3300071275 | MDA | Water | OANGC | p__Proteobacteria;c__Gammaproteobacteria;o__Burkholderiales;f__Methylophilaceae | present | FALSE | LQ | absent |
| EnvBac_Water_WGAX_A11_scBac_plate.B | 1M | novseq 6000 | PRJNA1052864 | SAMN38845169 | 3300071260 | WGAX | Water | OANGG | p__Proteobacteria;c__Gammaproteobacteria;o__Pseudomonadales;f__Nitrincolaceae | absent | FALSE | LQ | absent |
| EnvBac_Water_WGAX_B11_scBac_plate.B | 1M | novseq 6000 | PRJNA1052865 | SAMN38845220 | 3300071224 | WGAX | Water | OANGH | unassigned | absent | FALSE | LQ | absent |
| EnvBac_Water_WGAX_C11_scBac_plate.B | 1M | novseq 6000 | PRJNA1052866 | SAMN38845208 | 3300071225 | WGAX | Water | OANGN | unassigned | present | FALSE | LQ | absent |
| EnvBac_Water_WGAX_D11_scBac_plate.B | 1M | novseq 6000 | PRJNA1052867 | SAMN38845171 | 3300071276 | WGAX | Water | OANGO | unassigned | absent | FALSE | LQ | absent |
| EnvBac_Water_WGAX_E11_scBac_plate.B | 1M | novseq 6000 | PRJNA1052868 | SAMN38845191 | 3300071261 | WGAX | Water | OANGP | unassigned | absent | FALSE | LQ | absent |
| EnvBac_Water_WGAX_F11_scBac_plate.B | 1M | novseq 6000 | PRJNA1052869 | SAMN38845151 | 3300071262 | WGAX | Water | OANGS | unassigned | absent | FALSE | LQ | absent |
| EnvBac_Water_WGAX_G11_scBac_plate.B | 1M | novseq 6000 | PRJNA1052870 | SAMN38845141 | 3300071226 | WGAX | Water | OANGT | unassigned | absent | FALSE | LQ | absent |
| EnvBac_Water_WGAX_H11_scBac_plate.B | 1M | novseq 6000 | PRJNA1052871 | SAMN38845189 | 3300071263 | WGAX | Water | OANGU | unassigned | absent | FALSE | LQ | absent |
| EnvBac_Water_WGAX_A12_scBac_plate.B | 1M | novseq 6000 | PRJNA1052872 | SAMN38845153 | 3300071277 | WGAX | Water | OANGW | p__Bacteroidota;c__Bacteroidia;o__Flavobacteriales;f__Flavobacteriaceae | absent | FALSE | LQ | absent |
| EnvBac_Water_WGAX_B12_scBac_plate.B | 1M | novseq 6000 | PRJNA1052873 | SAMN38845144 | 3300071227 | WGAX | Water | OANGX | p__Proteobacteria;c__Gammaproteobacteria;o__Pseudomonadales;f__ | absent | FALSE | LQ | absent |
| EnvBac_Water_WGAX_C12_scBac_plate.B | 1M | novseq 6000 | PRJNA1052874 | SAMN38845213 | 3300071228 | WGAX | Water | OANGY | p__Proteobacteria;c__Gammaproteobacteria;o__Pseudomonadales;f__Pseudohongiellaceae | absent | FALSE | LQ | absent |
| EnvBac_Water_WGAX_D12_scBac_plate.B | 1M | novseq 6000 | PRJNA1052875 | SAMN38845203 | 3300071229 | WGAX | Water | OANGZ | unassigned | absent | FALSE | LQ | absent |
| EnvBac_Water_WGAX_F12_scBac_plate.B | 1M | novseq 6000 | PRJNA1052876 | SAMN38845231 | 3300071230 | WGAX | Water | OANHA | p__Bacteroidota;c__Bacteroidia;o__Flavobacteriales;f__Schleiferiaceae | present | FALSE | LQ | absent |
| EnvBac_Water_WGAX_G12_scBac_plate.B | 1M | novseq 6000 | PRJNA1052877 | SAMN38845145 | 3300071264 | WGAX | Water | OANHB | unassigned | absent | FALSE | LQ | absent |
| EnvBac_Water_WGAX_H12_scBac_plate.B | 1M | novseq 6000 | PRJNA1052878 | SAMN38845232 | 3300071265 | WGAX | Water | OANHC | unassigned | absent | FALSE | LQ | absent |

**Supplementary Table S5: (Continued).**

| genome_id | subsampling | seq_platform | BioProject | BioSample | IMG Genome ID | amp_method | system | library_name | gtdb_taxonomy | bgc_pa | Enzymatic Lysis | misag | rrna_16S_hit |
|---|---|---|---|---|---|---|---|---|---|---|---|---|---|
| EnvBac_Soil_PTA_F04_scBac_plate.C | 1M | novseq 6000 | PRJNA1052773 | SAMN38845148 | 3300071231 | PTA | Soil | OANNT | p__Proteobacteria;c__Gammaproteobacteria;o__Burkholderiales;f__UKL13-2 | present | TRUE | LQ | absent |
| EnvBac_Soil_PTA_A05_scBac_plate.C | 1M | novseq 6000 | PRJNA1052774 | SAMN38845149 | 3300071290 | PTA | Soil | OANNX | p__Proteobacteria;c__Gammaproteobacteria;o__Nevskiales;f__Nevskiaceae | present | TRUE | MQ | absent |
| EnvBac_Soil_PTA_B05_scBac_plate.C | 1M | novseq 6000 | PRJNA1052775 | SAMN38845185 | 3300071232 | PTA | Soil | OANNY | p__Proteobacteria;c__Gammaproteobacteria;o__Burkholderiales;f__SG8-39 | absent | TRUE | MQ | absent |
| EnvBac_Soil_PTA_C05_scBac_plate.C | 1M | novseq 6000 | PRJNA1052776 | SAMN38845233 | 3300071233 | PTA | Soil | OANNZ | p__Gemmatimonadota;c__Gemmatimonadetes;o__Gemmatimonadales;f__Gemmatimonadaceae | present | TRUE | MQ | absent |
| EnvBac_Soil_Wchmk_E07_scBac_plate.C | 1M | novseq 6000 | PRJNA1052777 | SAMN38845204 | 3300071266 | MDA | Soil | OANOT | p__Verrucomicrobiota;c__Verrucomicrobiae;o__Opitutales;f__Opitutaceae | absent | FALSE | LQ | absent |
| EnvBac_Soil_Wchmk_B08_scBac_plate.C | 1M | novseq 6000 | PRJNA1052778 | SAMN38845182 | 3300071267 | MDA | Soil | OANOZ | unassigned | present | FALSE | LQ | absent |
| EnvBac_Soil_WGAX_A10_scBac_plate.C | 1M | novseq 6000 | PRJNA1052779 | SAMN38845230 | 3300071278 | WGAX | Soil | OANPO | p__Bacteroidota;c__Bacteroidia;o__NS11-12g;f__ | absent | FALSE | LQ | absent |
| EnvBac_Soil_WGAX_B10_scBac_plate.C | 1M | novseq 6000 | PRJNA1052780 | SAMN38845183 | 3300071234 | WGAX | Soil | OANPP | unassigned | absent | FALSE | LQ | absent |
| EnvBac_Soil_WGAX_C10_scBac_plate.C | 1M | novseq 6000 | PRJNA1052781 | SAMN38845147 | 3300071235 | WGAX | Soil | OANPS | unassigned | present | FALSE | LQ | absent |
| EnvBac_Soil_WGAX_D10_scBac_plate.C | 1M | novseq 6000 | PRJNA1052782 | SAMN38845253 | 3300071236 | WGAX | Soil | OANPT | unassigned | absent | FALSE | LQ | absent |
| EnvBac_Soil_WGAX_E10_scBac_plate.C | 1M | novseq 6000 | PRJNA1052783 | SAMN38845184 | 3300071291 | WGAX | Soil | OANPU | unassigned | absent | FALSE | LQ | absent |
| EnvBac_Soil_WGAX_F10_scBac_plate.C | 1M | novseq 6000 | PRJNA1052784 | SAMN38845244 | 3300071237 | WGAX | Soil | OANPW | p__Actinobacteriota;c__Thermoleophilia;o__Gaiellales;f__Gaiellaceae | absent | FALSE | LQ | absent |
| EnvBac_Soil_WGAX_C11_scBac_plate.C | 1M | novseq 6000 | PRJNA1052785 | SAMN38845164 | 3300071293 | WGAX | Soil | OANSB | unassigned | absent | FALSE | LQ | absent |
| EnvBac_Soil_WGAX_D11_scBac_plate.C | 1M | novseq 6000 | PRJNA1052786 | SAMN38845214 | 3300071294 | WGAX | Soil | OANSC | unassigned | absent | FALSE | LQ | absent |
| EnvBac_Water_edMDA_A04_scBac_plate.A | 1M | novseq 6000 | PRJNA1052822 | SAMN38845250 | 3300071190 | MDA | Water | OANSP | unassigned | absent | TRUE | LQ | absent |
| EnvBac_Water_edMDA_B04_scBac_plate.A | 1M | novseq 6000 | PRJNA1052823 | SAMN38845158 | 3300071292 | MDA | Water | OANSS | p__Bacteroidota;c__Bacteroidia;o__Flavobacteriales;f__Flavobacteriaceae | absent | TRUE | LQ | absent |
| EnvBac_Water_edMDA_C04_scBac_plate.A | 1M | novseq 6000 | PRJNA1052824 | SAMN38845226 | 3300071191 | MDA | Water | OANST | p__Proteobacteria;c__Gammaproteobacteria;o__Chromatiales;f__Sedimenticolaceae | present | TRUE | LQ | absent |
| EnvBac_Water_edMDA_D04_scBac_plate.A | 1M | novseq 6000 | PRJNA1052825 | SAMN38845217 | 3300071192 | MDA | Water | OANSU | p__Bacteroidota;c__Rhodothermia;o__Balneolales;f__Balneolaceae | present | TRUE | LQ | absent |
| EnvBac_Water_edMDA_F04_scBac_plate.A | 1M | novseq 6000 | PRJNA1052826 | SAMN38845193 | 3300071193 | MDA | Water | OANSX | unassigned | absent | TRUE | LQ | absent |
| EnvBac_Water_edMDA_G04_scBac_plate.A | 1M | novseq 6000 | PRJNA1052827 | SAMN38845227 | 3300071280 | MDA | Water | OANSY | p__Proteobacteria;c__Gammaproteobacteria;o__Burkholderiales;f__Methylophilaceae | present | TRUE | LQ | absent |
| EnvBac_Water_edMDA_H04_scBac_plate.A | 1M | novseq 6000 | PRJNA1052828 | SAMN38845177 | 3300071281 | MDA | Water | OANSZ | p__Proteobacteria;c__Gammaproteobacteria;o__Burkholderiales;f__Methylophilaceae | present | TRUE | LQ | absent |
| EnvBac_Water_edMDA_A05_scBac_plate.A | 1M | novseq 6000 | PRJNA1052829 | SAMN38845252 | 3300071194 | MDA | Water | OANTA | unassigned | absent | TRUE | LQ | absent |
| EnvBac_Water_edMDA_C05_scBac_plate.A | 1M | novseq 6000 | PRJNA1052830 | SAMN38845167 | 3300071195 | MDA | Water | OANTC | p__Proteobacteria;c__Alphaproteobacteria;o__Puniceispirillales;f__UBA1172 | absent | TRUE | LQ | absent |
| EnvBac_Water_edMDA_E05_scBac_plate.A | 1M | novseq 6000 | PRJNA1052831 | SAMN38845228 | 3300071196 | MDA | Water | OANTH | p__Bacteroidota;c__Bacteroidia;o__Flavobacteriales;f__Schleiferiaceae | absent | TRUE | LQ | absent |
| EnvBac_Water_edMDA_F05_scBac_plate.A | 1M | novseq 6000 | PRJNA1052832 | SAMN38845246 | 3300071238 | MDA | Water | OANTN | p__Proteobacteria;c__Gammaproteobacteria;o__Pseudomonadales;f__Litoricolaceae | absent | TRUE | LQ | absent |
| EnvBac_Water_edMDA_G05_scBac_plate.A | 1M | novseq 6000 | PRJNA1052833 | SAMN38845165 | 3300071197 | MDA | Water | OANTO | unassigned | absent | TRUE | LQ | absent |
| EnvBac_Water_edMDA_H05_scBac_plate.A | 1M | novseq 6000 | PRJNA1052834 | SAMN38845206 | 3300071239 | MDA | Water | OANTP | p__Bacteroidota;c__Bacteroidia;o__Flavobacteriales;f__Schleiferiaceae | absent | TRUE | LQ | absent |
| EnvBac_Water_edMDA_A06_scBac_plate.A | 1M | novseq 6000 | PRJNA1052835 | SAMN38845236 | 3300071181 | MDA | Water | OANTS | p__Proteobacteria;c__Alphaproteobacteria;o__Pelagibacterales;f__Pelagibacteraceae | present | TRUE | MQ | absent |
| EnvBac_Water_edMDA_B06_scBac_plate.A | 1M | novseq 6000 | PRJNA1052836 | SAMN38845216 | 3300071182 | MDA | Water | OANTT | p__Bacteroidota;c__Bacteroidia;o__Flavobacteriales;f__Flavobacteriaceae | absent | TRUE | LQ | absent |
| EnvBac_Water_edMDA_C06_scBac_plate.A | 1M | novseq 6000 | PRJNA1052837 | SAMN38845207 | 3300071183 | MDA | Water | OANTU | p__Proteobacteria;c__Gammaproteobacteria;o__Enterobacterales;f__Alteromonadaceae | absent | TRUE | LQ | absent |
| EnvBac_Water_edMDA_E06_scBac_plate.A | 1M | novseq 6000 | PRJNA1052838 | SAMN38845218 | 3300071184 | MDA | Water | OANTX | p__Bacteroidota;c__Bacteroidia;o__Flavobacteriales;f__Flavobacteriaceae | absent | TRUE | LQ | absent |
| EnvBac_Water_edMDA_F06_scBac_plate.A | 1M | novseq 6000 | PRJNA1052839 | SAMN38845150 | 3300071185 | MDA | Water | OANTY | p__Bacteroidota;c__Bacteroidia;o__Flavobacteriales;f__Flavobacteriaceae | present | TRUE | LQ | absent |
| EnvBac_Water_edMDA_G06_scBac_plate.A | 1M | novseq 6000 | PRJNA1052840 | SAMN38845248 | 3300071186 | MDA | Water | OANTZ | unassigned | present | TRUE | LQ | absent |
| EnvBac_Water_edMDA_H06_scBac_plate.A | 1M | novseq 6000 | PRJNA1052841 | SAMN38845235 | 3300071187 | MDA | Water | OANUA | p__Proteobacteria;c__Alphaproteobacteria;o__Pelagibacterales;f__Pelagibacteraceae | absent | TRUE | LQ | absent |
| EnvBac_Soil_edMDA_A08_scBac_plate.A | 1M | novseq 6000 | PRJNA1052842 | SAMN38845139 | 3300071188 | MDA | Soil | OANUB | p__Acidobacteriota;c__Vicinamibacteria;o__Vicinamibacterales;f__UBA2999 | present | TRUE | LQ | absent |
| EnvBac_Soil_edMDA_B08_scBac_plate.A | 1M | novseq 6000 | PRJNA1052843 | SAMN38845188 | 3300071189 | MDA | Soil | OANUC | unassigned | absent | TRUE | LQ | absent |
| EnvBac_Soil_edMDA_D08_scBac_plate.A | 1M | novseq 6000 | PRJNA1052844 | SAMN38845222 | 3300071240 | MDA | Soil | OANUH | unassigned | present | TRUE | LQ | absent |
| EnvBac_Soil_edMDA_E08_scBac_plate.A | 1M | novseq 6000 | PRJNA1052845 | SAMN38845209 | 3300071241 | MDA | Soil | OANUN | unassigned | present | TRUE | LQ | absent |
| EnvBac_Soil_edMDA_F08_scBac_plate.A | 1M | novseq 6000 | PRJNA1052846 | SAMN38845239 | 3300071242 | MDA | Soil | OANUO | p__Proteobacteria;c__Alphaproteobacteria;o__Sphingomonadales;f__Sphingomonadaceae | present | TRUE | LQ | absent |
| EnvBac_Soil_edMDA_B09_scBac_plate.A | 1M | novseq 6000 | PRJNA1052847 | SAMN38845143 | 3300071198 | MDA | Soil | OANUW | p__Gemmatimonadota;c__Gemmatimonadetes;o__Gemmatimonadales;f__Gemmatimonadaceae | absent | TRUE | LQ | absent |
| EnvBac_Soil_edMDA_C09_scBac_plate.A | 1M | novseq 6000 | PRJNA1052848 | SAMN38845173 | 3300071199 | MDA | Soil | OANUX | unassigned | absent | TRUE | LQ | absent |
| EnvBac_Soil_edMDA_D09_scBac_plate.A | 1M | novseq 6000 | PRJNA1052849 | SAMN38845142 | 3300071200 | MDA | Soil | OANUY | unassigned | present | TRUE | LQ | absent |
| EnvBac_Soil_edMDA_E09_scBac_plate.A | 1M | novseq 6000 | PRJNA1052850 | SAMN38845240 | 3300071201 | MDA | Soil | OANUZ | unassigned | absent | TRUE | LQ | absent |
| EnvBac_Soil_edMDA_F09_scBac_plate.A | 1M | novseq 6000 | PRJNA1052787 | SAMN38845154 | 3300071243 | MDA | Soil | OANWA | unassigned | absent | TRUE | LQ | absent |
| EnvBac_Soil_edMDA_G09_scBac_plate.A | 1M | novseq 6000 | PRJNA1052788 | SAMN38845229 | 3300071244 | MDA | Soil | OANWB | unassigned | absent | TRUE | LQ | absent |
| EnvBac_Water_PTA_D06_scBac_plate.B | 20M | novseq 6000 | PRJNA1052789 | SAMN38845241 | 3300071202 | PTA | Water | OAHZZ | p__Proteobacteria;c__Gammaproteobacteria;o__Pseudomonadales;f__Nitrincolaceae | present | TRUE | HQ | absent |
| EnvBac_Water_WGAX_E12_scBac_plate.B | 20M | novseq 6000 | PRJNA1052790 | SAMN38845225 | 3300071203 | WGAX | Water | OANAA | p__Bacteroidota;c__Bacteroidia;o__Flavobacteriales;f__Schleiferiaceae | present | FALSE | LQ | absent |
| EnvBac_Water_PTA_A03_scBac_plate.B | 20M | novseq 6000 | PRJNA1052791 | SAMN38845175 | 3300071204 | PTA | Water | OANAB | p__Proteobacteria;c__Gammaproteobacteria;o__Pseudomonadales;f__HTCC2089 | present | TRUE | MQ | absent |
| EnvBac_Water_PTA_B03_scBac_plate.B | 20M | novseq 6000 | PRJNA1052792 | SAMN38845155 | 3300071245 | PTA | Water | OANAC | p__Bacteroidota;c__Bacteroidia;o__Flavobacteriales;f__Flavobacteriaceae | present | TRUE | MQ | absent |
| EnvBac_Water_PTA_C03_scBac_plate.B | 20M | novseq 6000 | PRJNA1052793 | SAMN38845156 | 3300071205 | PTA | Water | OANAG | p__Actinobacteriota;c__Actinomycetia;o__Actinomycetales;f__Microbacteriaceae | present | TRUE | MQ | absent |
| EnvBac_Water_PTA_D03_scBac_plate.B | 20M | novseq 6000 | PRJNA1052794 | SAMN38845251 | 3300071206 | PTA | Water | OANAH | p__Proteobacteria;c__Gammaproteobacteria;o__Enterobacterales;f__Alteromonadaceae | present | TRUE | MQ | absent |
| EnvBac_Water_PTA_E03_scBac_plate.B | 20M | novseq 6000 | PRJNA1052795 | SAMN38845210 | 3300071246 | PTA | Water | OANAN | p__Actinobacteriota;c__Acidimicrobiia;o__Acidimicrobiales;f__UBA11606 | absent | TRUE | MQ | absent |
| EnvBac_Water_PTA_F03_scBac_plate.B | 20M | novseq 6000 | PRJNA1052796 | SAMN38845157 | 3300071247 | PTA | Water | OANAO | p__Proteobacteria;c__Alphaproteobacteria;o__Pelagibacterales;f__Pelagibacteraceae | present | TRUE | MQ | absent |
| EnvBac_Water_PTA_G03_scBac_plate.B | 20M | novseq 6000 | PRJNA1052797 | SAMN38845159 | 3300071207 | PTA | Water | OANAP | p__Proteobacteria;c__Gammaproteobacteria;o__UBA4575;f__UBA4575 | present | TRUE | MQ | absent |
| EnvBac_Water_PTA_H03_scBac_plate.B | 20M | novseq 6000 | PRJNA1052798 | SAMN38845243 | 3300071282 | PTA | Water | OANAS | p__Proteobacteria;c__Gammaproteobacteria;o__Burkholderiales;f__Methylophilaceae | present | TRUE | MQ | absent |
| EnvBac_Water_PTA_A04_scBac_plate.B | 20M | novseq 6000 | PRJNA1052799 | SAMN38845163 | 3300071248 | PTA | Water | OANAT | p__Proteobacteria;c__Alphaproteobacteria;o__Pelagibacterales;f__Pelagibacteraceae | absent | TRUE | LQ | absent |
| EnvBac_Water_PTA_B04_scBac_plate.B | 20M | novseq 6000 | PRJNA1052800 | SAMN38845160 | 3300071208 | PTA | Water | OANAU | p__Proteobacteria;c__Alphaproteobacteria;o__HIMB59;f__HIMB59 | present | TRUE | MQ | absent |
| EnvBac_Water_PTA_C04_scBac_plate.B | 20M | novseq 6000 | PRJNA1052801 | SAMN38845194 | 3300071249 | PTA | Water | OANAW | p__Proteobacteria;c__Gammaproteobacteria;o__UBA4575;f__UBA4575 | absent | TRUE | LQ | absent |
| EnvBac_Water_PTA_D04_scBac_plate.B | 20M | novseq 6000 | PRJNA1052802 | SAMN38845211 | 3300071209 | PTA | Water | OANAX | p__Bacteroidota;c__Bacteroidia;o__Flavobacteriales;f__Flavobacteriaceae | present | TRUE | MQ | absent |
| EnvBac_Water_PTA_E04_scBac_plate.B | 20M | novseq 6000 | PRJNA1052803 | SAMN38845212 | 3300071210 | PTA | Water | OANAY | p__Bacteroidota;c__Bacteroidia;o__Flavobacteriales;f__Flavobacteriaceae | present | TRUE | MQ | absent |
| EnvBac_Water_PTA_F04_scBac_plate.B | 20M | novseq 6000 | PRJNA1052804 | SAMN38845178 | 3300071250 | PTA | Water | OANAZ | p__Proteobacteria;c__Alphaproteobacteria;o__Pelagibacterales;f__Pelagibacteraceae | absent | TRUE | MQ | absent |
| EnvBac_Water_PTA_G04_scBac_plate.B | 20M | novseq 6000 | PRJNA1052805 | SAMN38845179 | 3300071251 | PTA | Water | OANBA | p__Proteobacteria;c__Gammaproteobacteria;o__Chromatiales;f__Sedimenticolaceae | absent | TRUE | MQ | absent |
| EnvBac_Water_PTA_H04_scBac_plate.B | 20M | novseq 6000 | PRJNA1052806 | SAMN38845176 | 3300071211 | PTA | Water | OANBB | p__Proteobacteria;c__Gammaproteobacteria;o__Burkholderiales;f__Methylophilaceae | present | TRUE | MQ | absent |
| EnvBac_Water_PTA_A05_scBac_plate.B | 20M | novseq 6000 | PRJNA1052807 | SAMN38845215 | 3300071212 | PTA | Water | OANBC | p__Actinobacteriota;c__Actinomycetia;o__Actinomycetales;f__Microbacteriaceae | present | TRUE | HQ | absent |
| EnvBac_Water_PTA_B05_scBac_plate.B | 20M | novseq 6000 | PRJNA1052808 | SAMN38845205 | 3300071213 | PTA | Water | OANBG | p__Proteobacteria;c__Gammaproteobacteria;o__Pseudomonadales;f__Halieaceae | present | TRUE | HQ | absent |
| EnvBac_Water_PTA_C05_scBac_plate.B | 20M | novseq 6000 | PRJNA1052809 | SAMN38845237 | 3300071214 | PTA | Water | OANBH | p__Proteobacteria;c__Alphaproteobacteria;o__Rhodobacterales;f__Rhodobacteraceae | present | TRUE | HQ | absent |
| EnvBac_Water_PTA_D05_scBac_plate.B | 20M | novseq 6000 | PRJNA1052810 | SAMN38845247 | 3300071252 | PTA | Water | OANBN | p__Proteobacteria;c__Gammaproteobacteria;o__Enterobacterales;f__Alteromonadaceae | present | TRUE | MQ | absent |
| EnvBac_Water_PTA_E05_scBac_plate.B | 20M | novseq 6000 | PRJNA1052811 | SAMN38845242 | 3300071215 | PTA | Water | OANBO | p__Proteobacteria;c__Alphaproteobacteria;o__Rhodobacterales;f__Rhodobacteraceae | present | TRUE | HQ | absent |
| EnvBac_Water_PTA_F05_scBac_plate.B | 20M | novseq 6000 | PRJNA1052812 | SAMN38845168 | 3300071253 | PTA | Water | OANBP | p__Bacteroidota;c__Bacteroidia;o__Flavobacteriales;f__Schleiferiaceae | present | TRUE | MQ | absent |
| EnvBac_Water_PTA_G05_scBac_plate.B | 20M | novseq 6000 | PRJNA1052813 | SAMN38845219 | 3300071216 | PTA | Water | OANBS | p__Proteobacteria;c__Alphaproteobacteria;o__Rhodobacterales;f__Rhodobacteraceae | present | TRUE | MQ | absent |
| EnvBac_Water_PTA_H05_scBac_plate.B | 20M | novseq 6000 | PRJNA1052814 | SAMN38845221 | 3300071217 | PTA | Water | OANBT | p__Proteobacteria;c__Alphaproteobacteria;o__Rickettsiales;f__ | present | TRUE | MQ | absent |
| EnvBac_Water_PTA_A06_scBac_plate.B | 20M | novseq 6000 | PRJNA1052815 | SAMN38845170 | 3300071254 | PTA | Water | OANBU | p__Bacteroidota;c__Bacteroidia;o__Flavobacteriales;f__Schleiferiaceae | present | TRUE | LQ | absent |
| EnvBac_Water_PTA_B06_scBac_plate.B | 20M | novseq 6000 | PRJNA1052816 | SAMN38845224 | 3300071283 | PTA | Water | OANBW | p__Proteobacteria;c__Alphaproteobacteria;o__Pelagibacterales;f__Pelagibacteraceae | present | TRUE | MQ | absent |
| EnvBac_Water_PTA_C06_scBac_plate.B | 20M | novseq 6000 | PRJNA1052817 | SAMN38845140 | 3300071218 | PTA | Water | OANBX | p__Bacteroidota;c__Rhodothermia;o__Balneolales;f__Balneolaceae | present | TRUE | MQ | absent |
| EnvBac_Water_Wchmk_A09_scBac_plate.B | 20M | novseq 6000 | PRJNA1052818 | SAMN38845172 | 3300071274 | MDA | Water | OANCC | p__Proteobacteria;c__Alphaproteobacteria;o__Pelagibacterales;f__Pelagibacteraceae | absent | FALSE | MQ | absent |
| EnvBac_Water_Wchmk_B09_scBac_plate.B | 20M | novseq 6000 | PRJNA1052819 | SAMN38845152 | 3300071284 | MDA | Water | OANCG | p__Bacteroidota;c__Bacteroidia;o__Flavobacteriales;f__Flavobacteriaceae | absent | FALSE | LQ | absent |
| EnvBac_Water_Wchmk_C09_scBac_plate.B | 20M | novseq 6000 | PRJNA1052820 | SAMN38845238 | 3300071255 | MDA | Water | OANCH | p__Bacteroidota;c__Bacteroidia;o__Cytophagales;f__Cyclobacteriaceae | present | FALSE | LQ | absent |
| EnvBac_Water_Wchmk_D09_scBac_plate.B | 20M | novseq 6000 | PRJNA1052821 | SAMN38845249 | 3300071219 | MDA | Water | OANCN | p__Proteobacteria;c__Gammaproteobacteria;o__PS1;f__Thioglobaceae | absent | FALSE | LQ | absent |
| EnvBac_Water_Wchmk_E09_scBac_plate.B | 20M | novseq 6000 | PRJNA1052852 | SAMN38845174 | 3300071285 | MDA | Water | OANCO | p__Bacteroidota;c__Bacteroidia;o__Flavobacteriales;f__Flavobacteriaceae | present | FALSE | LQ | absent |
| EnvBac_Water_Wchmk_F09_scBac_plate.B | 20M | novseq 6000 | PRJNA1052853 | SAMN38845190 | 3300071286 | MDA | Water | OANCP | p__Actinobacteriota;c__Actinomycetia;o__Actinomycetales;f__Microbacteriaceae | absent | FALSE | LQ | absent |

**Supplementary Table S5: (Continued).**

| genome_id | subsampling | seq_platform | BioProject | BioSample | IMG Genome ID | amp_method | system | library_name | gtdb_taxonomy | bgc_pa | Enzymatic Lysis | misag | rrna_16S_hit |
|---|---|---|---|---|---|---|---|---|---|---|---|---|---|
| EnvBac_Water_Wchmk_G09_scBac_plate.B | 20M | novseq 6000 | PRJNA1052854 | SAMN38845161 | 3300071287 | MDA | Water | OANCS | p__Bacteroidota;c__Bacteroidia;o__Flavobacteriales;f__Flavobacteriaceae | present | FALSE | LQ | absent |
| EnvBac_Water_Wchmk_H09_scBac_plate.B | 20M | novseq 6000 | PRJNA1052855 | SAMN38845162 | 3300071220 | MDA | Water | OANCT | p__Bacteroidota;c__Bacteroidia;o__Flavobacteriales;f__Schleiferiaceae | present | FALSE | LQ | absent |
| EnvBac_Water_Wchmk_A10_scBac_plate.B | 20M | novseq 6000 | PRJNA1052856 | SAMN38845180 | 3300071256 | MDA | Water | OANCU | p__Actinobacteriota;c__Actinomycetia;o__Actinomycetales;f__Microbacteriaceae | absent | FALSE | LQ | absent |
| EnvBac_Water_Wchmk_B10_scBac_plate.B | 20M | novseq 6000 | PRJNA1052857 | SAMN38845192 | 3300071221 | MDA | Water | OANCW | p__Bacteroidota;c__Bacteroidia;o__Flavobacteriales;f__Flavobacteriaceae | absent | FALSE | LQ | absent |
| EnvBac_Water_Wchmk_C10_scBac_plate.B | 20M | novseq 6000 | PRJNA1052858 | SAMN38845181 | 3300071257 | MDA | Water | OANCX | p__Proteobacteria;c__Alphaproteobacteria;o__Rhodobacterales;f__Rhodobacteraceae | present | FALSE | LQ | absent |
| EnvBac_Water_Wchmk_D10_scBac_plate.B | 20M | novseq 6000 | PRJNA1052859 | SAMN38845186 | 3300071222 | MDA | Water | OANCY | p__Bacteroidota;c__Bacteroidia;o__Flavobacteriales;f__Flavobacteriaceae | present | FALSE | MQ | absent |
| EnvBac_Water_Wchmk_E10_scBac_plate.B | 20M | novseq 6000 | PRJNA1052860 | SAMN38845234 | 3300071258 | MDA | Water | OANCZ | p__Proteobacteria;c__Alphaproteobacteria;o__Rhodobacterales;f__Rhodobacteraceae | absent | FALSE | LQ | absent |
| EnvBac_Water_Wchmk_F10_scBac_plate.B | 20M | novseq 6000 | PRJNA1052861 | SAMN38845166 | 3300071259 | MDA | Water | OANGA | p__Proteobacteria;c__Gammaproteobacteria;o__Chromatiales;f__Sedimenticolaceae | absent | FALSE | LQ | absent |
| EnvBac_Water_Wchmk_G10_scBac_plate.B | 20M | novseq 6000 | PRJNA1052862 | SAMN38845187 | 3300071223 | MDA | Water | OANGB | p__Proteobacteria;c__Alphaproteobacteria;o__Rhodobacterales;f__Rhodobacteraceae | absent | FALSE | LQ | absent |
| EnvBac_Water_Wchmk_H10_scBac_plate.B | 20M | novseq 6000 | PRJNA1052863 | SAMN38845223 | 3300071275 | MDA | Water | OANGC | p__Proteobacteria;c__Gammaproteobacteria;o__Burkholderiales;f__Methylophilaceae | present | FALSE | LQ | absent |
| EnvBac_Water_WGAX_A11_scBac_plate.B | 20M | novseq 6000 | PRJNA1052864 | SAMN38845169 | 3300071260 | WGAX | Water | OANGG | p__Proteobacteria;c__Gammaproteobacteria;o__Pseudomonadales;f__Nitrincolaceae | present | FALSE | LQ | absent |
| EnvBac_Water_WGAX_B11_scBac_plate.B | 20M | novseq 6000 | PRJNA1052865 | SAMN38845220 | 3300071224 | WGAX | Water | OANGH | p__Bacteroidota;c__Bacteroidia;o__Cytophagales;f__Cyclobacteriaceae | absent | FALSE | LQ | absent |
| EnvBac_Water_WGAX_C11_scBac_plate.B | 20M | novseq 6000 | PRJNA1052866 | SAMN38845208 | 3300071225 | WGAX | Water | OANGN | p__Proteobacteria;c__Alphaproteobacteria;o__Rhodobacterales;f__Rhodobacteraceae | present | FALSE | LQ | absent |
| EnvBac_Water_WGAX_D11_scBac_plate.B | 20M | novseq 6000 | PRJNA1052867 | SAMN38845171 | 3300071276 | WGAX | Water | OANGO | p__Proteobacteria;c__Alphaproteobacteria;o__HIMB59;f__HIMB59 | absent | FALSE | LQ | absent |
| EnvBac_Water_WGAX_E11_scBac_plate.B | 20M | novseq 6000 | PRJNA1052868 | SAMN38845191 | 3300071261 | WGAX | Water | OANGP | p__Proteobacteria;c__Gammaproteobacteria;o__Pseudomonadales;f__DSM-100316 | absent | FALSE | LQ | absent |
| EnvBac_Water_WGAX_F11_scBac_plate.B | 20M | novseq 6000 | PRJNA1052869 | SAMN38845151 | 3300071262 | WGAX | Water | OANGS | unassigned | absent | FALSE | LQ | absent |
| EnvBac_Water_WGAX_G11_scBac_plate.B | 20M | novseq 6000 | PRJNA1052870 | SAMN38845141 | 3300071226 | WGAX | Water | OANGT | p__Bacteroidota;c__Bacteroidia;o__Flavobacteriales;f__Schleiferiaceae | absent | FALSE | LQ | absent |
| EnvBac_Water_WGAX_H11_scBac_plate.B | 20M | novseq 6000 | PRJNA1052871 | SAMN38845189 | 3300071263 | WGAX | Water | OANGU | p__Actinobacteriota;c__Actinomycetia;o__Actinomycetales;f__Microbacteriaceae | absent | FALSE | LQ | absent |
| EnvBac_Water_WGAX_A12_scBac_plate.B | 20M | novseq 6000 | PRJNA1052872 | SAMN38845153 | 3300071277 | WGAX | Water | OANGW | p__Bacteroidota;c__Bacteroidia;o__Flavobacteriales;f__Flavobacteriaceae | absent | FALSE | LQ | absent |
| EnvBac_Water_WGAX_B12_scBac_plate.B | 20M | novseq 6000 | PRJNA1052873 | SAMN38845144 | 3300071227 | WGAX | Water | OANGX | p__Proteobacteria;c__Gammaproteobacteria;o__Pseudomonadales;f__ | present | FALSE | MQ | absent |
| EnvBac_Water_WGAX_C12_scBac_plate.B | 20M | novseq 6000 | PRJNA1052874 | SAMN38845213 | 3300071228 | WGAX | Water | OANGY | p__Proteobacteria;c__Gammaproteobacteria;o__Pseudomonadales;f__Pseudohongiellaceae | present | FALSE | LQ | absent |
| EnvBac_Water_WGAX_D12_scBac_plate.B | 20M | novseq 6000 | PRJNA1052875 | SAMN38845203 | 3300071229 | WGAX | Water | OANGZ | p__Proteobacteria;c__Alphaproteobacteria;o__Rhodobacterales;f__Rhodobacteraceae | present | FALSE | LQ | absent |
| EnvBac_Water_WGAX_F12_scBac_plate.B | 20M | novseq 6000 | PRJNA1052876 | SAMN38845231 | 3300071230 | WGAX | Water | OANHA | p__Bacteroidota;c__Bacteroidia;o__Flavobacteriales;f__Schleiferiaceae | present | FALSE | LQ | absent |
| EnvBac_Water_WGAX_G12_scBac_plate.B | 20M | novseq 6000 | PRJNA1052877 | SAMN38845145 | 3300071264 | WGAX | Water | OANHB | unassigned | absent | FALSE | LQ | absent |
| EnvBac_Water_WGAX_H12_scBac_plate.B | 20M | novseq 6000 | PRJNA1052878 | SAMN38845232 | 3300071265 | WGAX | Water | OANHC | p__Proteobacteria;c__Gammaproteobacteria;o__Chromatiales;f__Sedimenticolaceae | absent | FALSE | LQ | absent |
| EnvBac_Soil_PTA_G02_scBac_plate.C | 20M | novseq 6000 | PRJNA1052771 | SAMN38845245 | 3300071288 | PTA | Soil | OANHT | p__Acidobacteriota;c__Vicinamibacteria;o__Vicinamibacterales;f__UBA2999 | absent | TRUE | MQ | absent |
| EnvBac_Soil_PTA_B04_scBac_plate.C | 20M | novseq 6000 | PRJNA1052772 | SAMN38845146 | 3300071289 | PTA | Soil | OANNN | p__Bacteroidota;c__Bacteroidia;o__NS11-12g;f__ | present | TRUE | MQ | absent |
| EnvBac_Soil_PTA_F04_scBac_plate.C | 20M | novseq 6000 | PRJNA1052773 | SAMN38845148 | 3300071231 | PTA | Soil | OANNT | p__Proteobacteria;c__Gammaproteobacteria;o__Burkholderiales;f__UKL13-2 | present | TRUE | MQ | absent |
| EnvBac_Soil_PTA_A05_scBac_plate.C | 20M | novseq 6000 | PRJNA1052774 | SAMN38845149 | 3300071290 | PTA | Soil | OANNX | p__Proteobacteria;c__Gammaproteobacteria;o__Nevskiales;f__Nevskiaceae | present | TRUE | LQ | absent |
| EnvBac_Soil_PTA_B05_scBac_plate.C | 20M | novseq 6000 | PRJNA1052775 | SAMN38845185 | 3300071232 | PTA | Soil | OANNY | p__Proteobacteria;c__Gammaproteobacteria;o__Burkholderiales;f__SG8-39 | present | TRUE | MQ | absent |
| EnvBac_Soil_PTA_C05_scBac_plate.C | 20M | novseq 6000 | PRJNA1052776 | SAMN38845233 | 3300071233 | PTA | Soil | OANNZ | p__Gemmatimonadota;c__Gemmatimonadetes;o__Gemmatimonadales;f__Gemmatimonadaceae | present | TRUE | MQ | absent |
| EnvBac_Soil_Wchmk_E07_scBac_plate.C | 20M | novseq 6000 | PRJNA1052777 | SAMN38845204 | 3300071266 | MDA | Soil | OANOT | p__Verrucomicrobiota;c__Verrucomicrobiae;o__Opitutales;f__Opitutaceae | present | FALSE | LQ | absent |
| EnvBac_Soil_Wchmk_B08_scBac_plate.C | 20M | novseq 6000 | PRJNA1052778 | SAMN38845182 | 3300071267 | MDA | Soil | OANOZ | p__Bacteroidota;c__Bacteroidia;o__Sphingobacteriales;f__Sphingobacteriaceae | absent | FALSE | LQ | absent |
| EnvBac_Soil_WGAX_A10_scBac_plate.C | 20M | novseq 6000 | PRJNA1052779 | SAMN38845230 | 3300071278 | WGAX | Soil | OANPO | p__Bacteroidota;c__Bacteroidia;o__NS11-12g;f__ | absent | FALSE | LQ | absent |
| EnvBac_Soil_WGAX_B10_scBac_plate.C | 20M | novseq 6000 | PRJNA1052780 | SAMN38845183 | 3300071234 | WGAX | Soil | OANPP | unassigned | absent | FALSE | LQ | absent |
| EnvBac_Soil_WGAX_C10_scBac_plate.C | 20M | novseq 6000 | PRJNA1052781 | SAMN38845147 | 3300071235 | WGAX | Soil | OANPS | p__Acidobacteriota;c__Vicinamibacteria;o__Vicinamibacterales;f__UBA2999 | present | FALSE | LQ | absent |
| EnvBac_Soil_WGAX_D10_scBac_plate.C | 20M | novseq 6000 | PRJNA1052782 | SAMN38845253 | 3300071236 | WGAX | Soil | OANPT | unassigned | absent | FALSE | LQ | absent |
| EnvBac_Soil_WGAX_E10_scBac_plate.C | 20M | novseq 6000 | PRJNA1052783 | SAMN38845184 | 3300071291 | WGAX | Soil | OANPU | unassigned | present | FALSE | LQ | absent |
| EnvBac_Soil_WGAX_F10_scBac_plate.C | 20M | novseq 6000 | PRJNA1052784 | SAMN38845244 | 3300071237 | WGAX | Soil | OANPW | p__Actinobacteriota;c__Thermoleophilia;o__Gaiellales;f__Gaiellaceae | absent | FALSE | LQ | absent |
| EnvBac_Soil_WGAX_C11_scBac_plate.C | 20M | novseq 6000 | PRJNA1052785 | SAMN38845164 | 3300071293 | WGAX | Soil | OANSB | p__Firmicutes;c__Bacilli;o__Lactobacillales;f__Streptococcaceae | absent | FALSE | LQ | absent |
| EnvBac_Soil_WGAX_D11_scBac_plate.C | 20M | novseq 6000 | PRJNA1052786 | SAMN38845214 | 3300071294 | WGAX | Soil | OANSC | unassigned | absent | FALSE | LQ | absent |
| EnvBac_Water_edMDA_A04_scBac_plate.A | 20M | novseq 6000 | PRJNA1052822 | SAMN38845250 | 3300071190 | MDA | Water | OANSP | p__Bacteroidota;c__Bacteroidia;o__Flavobacteriales;f__Flavobacteriaceae | absent | TRUE | LQ | absent |
| EnvBac_Water_edMDA_B04_scBac_plate.A | 20M | novseq 6000 | PRJNA1052823 | SAMN38845158 | 3300071292 | MDA | Water | OANSS | p__Bacteroidota;c__Bacteroidia;o__Flavobacteriales;f__Flavobacteriaceae | present | TRUE | LQ | absent |
| EnvBac_Water_edMDA_C04_scBac_plate.A | 20M | novseq 6000 | PRJNA1052824 | SAMN38845226 | 3300071191 | MDA | Water | OANST | p__Proteobacteria;c__Gammaproteobacteria;o__Chromatiales;f__Sedimenticolaceae | present | TRUE | LQ | absent |
| EnvBac_Water_edMDA_D04_scBac_plate.A | 20M | novseq 6000 | PRJNA1052825 | SAMN38845217 | 3300071192 | MDA | Water | OANSU | p__Bacteroidota;c__Rhodothermia;o__Balneolales;f__Balneolaceae | present | TRUE | LQ | absent |
| EnvBac_Water_edMDA_F04_scBac_plate.A | 20M | novseq 6000 | PRJNA1052826 | SAMN38845193 | 3300071193 | MDA | Water | OANSX | p__Proteobacteria;c__Gammaproteobacteria;o__Pseudomonadales;f__Litoricolaceae | absent | TRUE | LQ | absent |
| EnvBac_Water_edMDA_G04_scBac_plate.A | 20M | novseq 6000 | PRJNA1052827 | SAMN38845227 | 3300071280 | MDA | Water | OANSY | p__Proteobacteria;c__Gammaproteobacteria;o__Burkholderiales;f__Methylophilaceae | present | TRUE | LQ | absent |
| EnvBac_Water_edMDA_H04_scBac_plate.A | 20M | novseq 6000 | PRJNA1052828 | SAMN38845177 | 3300071281 | MDA | Water | OANSZ | p__Proteobacteria;c__Gammaproteobacteria;o__Burkholderiales;f__Methylophilaceae | present | TRUE | LQ | absent |
| EnvBac_Water_edMDA_A05_scBac_plate.A | 20M | novseq 6000 | PRJNA1052829 | SAMN38845252 | 3300071194 | MDA | Water | OANTA | p__Bacteroidota;c__Bacteroidia;o__Flavobacteriales;f__CAIKAF01 | present | TRUE | LQ | absent |
| EnvBac_Water_edMDA_C05_scBac_plate.A | 20M | novseq 6000 | PRJNA1052830 | SAMN38845167 | 3300071195 | MDA | Water | OANTC | p__Proteobacteria;c__Alphaproteobacteria;o__Puniceispirillales;f__UBA1172 | present | TRUE | LQ | absent |
| EnvBac_Water_edMDA_E05_scBac_plate.A | 20M | novseq 6000 | PRJNA1052831 | SAMN38845228 | 3300071196 | MDA | Water | OANTH | p__Bacteroidota;c__Bacteroidia;o__Flavobacteriales;f__Schleiferiaceae | absent | TRUE | LQ | absent |
| EnvBac_Water_edMDA_F05_scBac_plate.A | 20M | novseq 6000 | PRJNA1052832 | SAMN38845246 | 3300071238 | MDA | Water | OANTN | p__Proteobacteria;c__Gammaproteobacteria;o__Pseudomonadales;f__Litoricolaceae | present | TRUE | LQ | absent |
| EnvBac_Water_edMDA_G05_scBac_plate.A | 20M | novseq 6000 | PRJNA1052833 | SAMN38845165 | 3300071197 | MDA | Water | OANTO | unassigned | absent | TRUE | LQ | absent |
| EnvBac_Water_edMDA_H05_scBac_plate.A | 20M | novseq 6000 | PRJNA1052834 | SAMN38845206 | 3300071239 | MDA | Water | OANTP | p__Bacteroidota;c__Bacteroidia;o__Flavobacteriales;f__Schleiferiaceae | absent | TRUE | LQ | absent |
| EnvBac_Water_edMDA_A06_scBac_plate.A | 20M | novseq 6000 | PRJNA1052835 | SAMN38845236 | 3300071181 | MDA | Water | OANTS | p__Proteobacteria;c__Alphaproteobacteria;o__Pelagibacterales;f__Pelagibacteraceae | present | TRUE | MQ | absent |
| EnvBac_Water_edMDA_B06_scBac_plate.A | 20M | novseq 6000 | PRJNA1052836 | SAMN38845216 | 3300071182 | MDA | Water | OANTT | p__Bacteroidota;c__Bacteroidia;o__Flavobacteriales;f__Flavobacteriaceae | absent | TRUE | LQ | absent |
| EnvBac_Water_edMDA_C06_scBac_plate.A | 20M | novseq 6000 | PRJNA1052837 | SAMN38845207 | 3300071183 | MDA | Water | OANTU | p__Proteobacteria;c__Gammaproteobacteria;o__Enterobacterales;f__Alteromonadaceae | absent | TRUE | LQ | absent |
| EnvBac_Water_edMDA_E06_scBac_plate.A | 20M | novseq 6000 | PRJNA1052838 | SAMN38845218 | 3300071184 | MDA | Water | OANTX | p__Bacteroidota;c__Bacteroidia;o__Flavobacteriales;f__Flavobacteriaceae | absent | TRUE | LQ | absent |
| EnvBac_Water_edMDA_F06_scBac_plate.A | 20M | novseq 6000 | PRJNA1052839 | SAMN38845150 | 3300071185 | MDA | Water | OANTY | p__Bacteroidota;c__Bacteroidia;o__Flavobacteriales;f__Flavobacteriaceae | present | TRUE | LQ | absent |
| EnvBac_Water_edMDA_G06_scBac_plate.A | 20M | novseq 6000 | PRJNA1052840 | SAMN38845248 | 3300071186 | MDA | Water | OANTZ | unassigned | present | TRUE | LQ | absent |
| EnvBac_Water_edMDA_H06_scBac_plate.A | 20M | novseq 6000 | PRJNA1052841 | SAMN38845235 | 3300071187 | MDA | Water | OANUA | p__Proteobacteria;c__Alphaproteobacteria;o__Pelagibacterales;f__Pelagibacteraceae | absent | TRUE | MQ | present |
| EnvBac_Soil_edMDA_A08_scBac_plate.A | 20M | novseq 6000 | PRJNA1052842 | SAMN38845139 | 3300071188 | MDA | Soil | OANUB | p__Acidobacteriota;c__Vicinamibacteria;o__Vicinamibacterales;f__UBA2999 | present | TRUE | MQ | absent |
| EnvBac_Soil_edMDA_B08_scBac_plate.A | 20M | novseq 6000 | PRJNA1052843 | SAMN38845188 | 3300071189 | MDA | Soil | OANUC | p__Proteobacteria;c__Alphaproteobacteria;o__Dongiales;f__Dongiaceae | present | TRUE | LQ | absent |
| EnvBac_Soil_edMDA_D08_scBac_plate.A | 20M | novseq 6000 | PRJNA1052844 | SAMN38845222 | 3300071240 | MDA | Soil | OANUH | p__Proteobacteria;c__Gammaproteobacteria;o__Burkholderiales;f__UKL13-2 | present | TRUE | LQ | absent |
| EnvBac_Soil_edMDA_E08_scBac_plate.A | 20M | novseq 6000 | PRJNA1052845 | SAMN38845209 | 3300071241 | MDA | Soil | OANUN | unassigned | present | TRUE | LQ | absent |
| EnvBac_Soil_edMDA_F08_scBac_plate.A | 20M | novseq 6000 | PRJNA1052846 | SAMN38845239 | 3300071242 | MDA | Soil | OANUO | p__Proteobacteria;c__Alphaproteobacteria;o__Sphingomonadales;f__Sphingomonadaceae | present | TRUE | LQ | absent |
| EnvBac_Soil_edMDA_B09_scBac_plate.A | 20M | novseq 6000 | PRJNA1052847 | SAMN38845143 | 3300071198 | MDA | Soil | OANUW | p__Gemmatimonadota;c__Gemmatimonadetes;o__Gemmatimonadales;f__Gemmatimonadaceae | present | TRUE | MQ | absent |
| EnvBac_Soil_edMDA_C09_scBac_plate.A | 20M | novseq 6000 | PRJNA1052848 | SAMN38845173 | 3300071199 | MDA | Soil | OANUX | unassigned | absent | TRUE | LQ | absent |
| EnvBac_Soil_edMDA_D09_scBac_plate.A | 20M | novseq 6000 | PRJNA1052849 | SAMN38845142 | 3300071200 | MDA | Soil | OANUY | p__Proteobacteria;c__Gammaproteobacteria;o__Burkholderiales;f__SG8-39 | present | TRUE | LQ | absent |
| EnvBac_Soil_edMDA_E09_scBac_plate.A | 20M | novseq 6000 | PRJNA1052850 | SAMN38845240 | 3300071201 | MDA | Soil | OANUZ | unassigned | absent | TRUE | LQ | absent |
| EnvBac_Soil_edMDA_F09_scBac_plate.A | 20M | novseq 6000 | PRJNA1052787 | SAMN38845154 | 3300071243 | MDA | Soil | OANWA | p__Bacteroidota;c__Bacteroidia;o__NS11-12g;f__UKL13-3 | absent | TRUE | LQ | absent |
| EnvBac_Soil_edMDA_G09_scBac_plate.A | 20M | novseq 6000 | PRJNA1052788 | SAMN38845229 | 3300071244 | MDA | Soil | OANWB | p__Bdellovibrionota;c__Bdellovibrionia;o__Bdellovibrionales;f__Bdellovibrionaceae | absent | TRUE | LQ | absent |

**Supplementary Table S5: (Continued).**

| genome_id | plasmid_gtr5kb | virus_gtr5kb | Completeness_(%) | Contamination_(%) | contig_count | genome_size_Mbp | Contig_N50 | GC_Content |
|---|---|---|---|---|---|---|---|---|
| scBac_PTA_EnviBac_sc_plate3well3A_S9 | absent | present | 91 | 0.03 | 76 | 1.9 | 52000 | 0.45 |
| scBac_PTA_EnviBac_sc_plate3well3B_S10 | absent | absent | 81 | 1.4 | 110 | 2.1 | 39000 | 0.44 |
| scBac_PTA_EnviBac_sc_plate3well3C_S11 | present | present | 86 | 0.98 | 210 | 2.8 | 19000 | 0.51 |
| scBac_PTA_EnviBac_sc_plate3well3D_S12 | present | absent | 20 | 1.6 | 220 | 1 | 5800 | 0.6 |
| scBac_PTA_EnviBac_sc_plate3well3E_S13 | present | present | 87 | 1 | 150 | 3 | 31000 | 0.49 |
| scBac_WGAX_EnviBac_sc_well3C_S95 | absent | absent | 15 | 0 | 35 | 0.44 | 20000 | 0.55 |
| scBac_WGAX_EnviBac_sc_well4H_S96 | present | absent | 11 | 0 | 31 | 0.36 | 15000 | 0.48 |
| scBac_WGAX_EnviBac_sc_well5B_S97 | absent | absent | 9.6 | 0.01 | 41 | 0.31 | 9700 | 0.52 |
| scBac_WGAX_EnviBac_sc_well6D_S99 | absent | absent | 26 | 0 | 74 | 0.86 | 20000 | 0.51 |
| scBac_WGAX_EnviBac_sc_well7D_S100 | absent | absent | 10 | 0.05 | 24 | 0.18 | 9800 | 0.49 |
| scBac_Watchmaker_EnviBac_sc_well3B_S79 | present | absent | 23 | 0.02 | 97 | 0.74 | 9700 | 0.51 |
| scBac_Watchmaker_EnviBac_sc_well3C_S80 | present | absent | 17 | 0.03 | 86 | 0.62 | 10000 | 0.51 |
| scBac_Watchmaker_EnviBac_sc_well4C_S81 | absent | absent | 12 | 0.07 | 68 | 0.49 | 9000 | 0.47 |
| scBac_Watchmaker_EnviBac_sc_well5A_S83 | present | absent | 36 | 0 | 64 | 1.1 | 37000 | 0.52 |
| scBac_Watchmaker_EnviBac_sc_well5E_S84 | present | absent | 10 | 0.01 | 47 | 0.45 | 15000 | 0.59 |
| scBac_Watchmaker_EnviBac_sc_well6E_S85 | present | absent | 20 | 0.04 | 66 | 0.62 | 13000 | 0.49 |
| scBac_Watchmaker_EnviBac_sc_well7E_S86 | absent | absent | 19 | 0.45 | 63 | 0.51 | 12000 | 0.45 |
| EnvBac_Water_PTA_D06_scBac_plate.B | present | present | 87 | 0.23 | 120 | 2.9 | 39000 | 0.52 |
| EnvBac_Water_WGAX_E12_scBac_plate.B | absent | absent | 17 | 0 | 35 | 0.47 | 18000 | 0.44 |
| EnvBac_Water_PTA_A03_scBac_plate.B | absent | present | 86 | 4.8 | 240 | 2.7 | 17000 | 0.63 |
| EnvBac_Water_PTA_B03_scBac_plate.B | absent | present | 79 | 2.7 | 120 | 1.9 | 31000 | 0.38 |
| EnvBac_Water_PTA_C03_scBac_plate.B | present | present | 74 | 4 | 110 | 1.4 | 39000 | 0.56 |
| EnvBac_Water_PTA_D03_scBac_plate.B | present | present | 77 | 2.6 | 210 | 2.2 | 16000 | 0.4 |
| EnvBac_Water_PTA_E03_scBac_plate.B | absent | absent | 39 | 1.1 | 190 | 1.1 | 7000 | 0.63 |
| EnvBac_Water_PTA_F03_scBac_plate.B | present | present | 91 | 2.4 | 87 | 1.4 | 51000 | 0.31 |
| EnvBac_Water_PTA_G03_scBac_plate.B | absent | absent | 84 | 0.02 | 26 | 1.4 | 1e+05 | 0.42 |
| EnvBac_Water_PTA_H03_scBac_plate.B | absent | present | 85 | 2.4 | 52 | 1.3 | 140000 | 0.35 |
| EnvBac_Water_PTA_A04_scBac_plate.B | absent | present | 71 | 5.1 | 160 | 1.1 | 9700 | 0.31 |
| EnvBac_Water_PTA_B04_scBac_plate.B | absent | absent | 73 | 0.13 | 50 | 1.2 | 230000 | 0.34 |
| EnvBac_Water_PTA_C04_scBac_plate.B | absent | present | 48 | 4.5 | 78 | 0.94 | 19000 | 0.42 |
| EnvBac_Water_PTA_D04_scBac_plate.B | absent | absent | 88 | 0.98 | 120 | 2.1 | 32000 | 0.4 |
| EnvBac_Water_PTA_E04_scBac_plate.B | absent | present | 89 | 0.55 | 95 | 2.1 | 45000 | 0.36 |
| EnvBac_Water_PTA_F04_scBac_plate.B | absent | absent | 83 | 1 | 120 | 1.2 | 17000 | 0.3 |
| EnvBac_Water_PTA_G04_scBac_plate.B | present | absent | 74 | 2.2 | 140 | 1.6 | 16000 | 0.47 |
| EnvBac_Water_PTA_H04_scBac_plate.B | absent | present | 98 | 2.5 | 44 | 1.4 | 91000 | 0.35 |
| EnvBac_Water_PTA_A05_scBac_plate.B | present | absent | 77 | 0.13 | 37 | 1.1 | 190000 | 0.54 |
| EnvBac_Water_PTA_B05_scBac_plate.B | absent | present | 81 | 0 | 41 | 2.5 | 130000 | 0.58 |
| EnvBac_Water_PTA_C05_scBac_plate.B | present | absent | 84 | 0.16 | 140 | 2.5 | 38000 | 0.5 |
| EnvBac_Water_PTA_D05_scBac_plate.B | present | absent | 69 | 1.4 | 140 | 2 | 25000 | 0.4 |
| EnvBac_Water_PTA_E05_scBac_plate.B | present | present | 88 | 1.5 | 180 | 2.6 | 22000 | 0.49 |
| EnvBac_Water_PTA_F05_scBac_plate.B | absent | absent | 99 | 1 | 100 | 2.1 | 34000 | 0.44 |
| EnvBac_Water_PTA_G05_scBac_plate.B | present | absent | 81 | 0.06 | 77 | 2.3 | 70000 | 0.55 |
| EnvBac_Water_PTA_H05_scBac_plate.B | present | present | 90 | 0.72 | 100 | 2.4 | 43000 | 0.42 |
| EnvBac_Water_PTA_A06_scBac_plate.B | present | absent | 79 | 3.2 | 150 | 2 | 22000 | 0.44 |
| EnvBac_Water_PTA_B06_scBac_plate.B | absent | present | 85 | 3.1 | 130 | 1.4 | 16000 | 0.3 |
| EnvBac_Water_PTA_C06_scBac_plate.B | absent | absent | 80 | 0.11 | 40 | 1.5 | 140000 | 0.45 |
| EnvBac_Water_Wchmk_A09_scBac_plate.B | present | absent | 37 | 0.01 | 58 | 0.51 | 15000 | 0.3 |
| EnvBac_Water_Wchmk_B09_scBac_plate.B | absent | absent | 31 | 0.03 | 61 | 0.59 | 18000 | 0.41 |
| EnvBac_Water_Wchmk_C09_scBac_plate.B | absent | absent | 11 | 0.14 | 68 | 0.42 | 7200 | 0.42 |
| EnvBac_Water_Wchmk_D09_scBac_plate.B | absent | absent | 31 | 0 | 59 | 0.41 | 8300 | 0.36 |
| EnvBac_Water_Wchmk_E09_scBac_plate.B | absent | absent | 26 | 0.01 | 73 | 0.6 | 10000 | 0.36 |
| EnvBac_Water_Wchmk_F09_scBac_plate.B | absent | absent | 9.9 | 0 | 41 | 0.22 | 6700 | 0.53 |
| EnvBac_Water_Wchmk_G09_scBac_plate.B | absent | absent | 23 | 0.01 | 55 | 0.49 | 14000 | 0.38 |
| EnvBac_Water_Wchmk_H09_scBac_plate.B | absent | absent | 18 | 0.01 | 57 | 0.43 | 9500 | 0.45 |
| EnvBac_Water_Wchmk_A10_scBac_plate.B | absent | absent | 9.1 | 0 | 25 | 0.24 | 13000 | 0.57 |
| EnvBac_Water_Wchmk_B10_scBac_plate.B | present | absent | 16 | 0 | 45 | 0.31 | 8800 | 0.46 |
| EnvBac_Water_Wchmk_C10_scBac_plate.B | present | absent | 12 | 0 | 68 | 0.45 | 8200 | 0.49 |
| EnvBac_Water_Wchmk_D10_scBac_plate.B | absent | absent | 39 | 0.38 | 74 | 0.86 | 16000 | 0.41 |
| EnvBac_Water_Wchmk_E10_scBac_plate.B | present | absent | 9.7 | 0 | 33 | 0.3 | 15000 | 0.57 |
| EnvBac_Water_Wchmk_F10_scBac_plate.B | absent | absent | 20 | 0.09 | 45 | 0.34 | 12000 | 0.47 |
| EnvBac_Water_Wchmk_G10_scBac_plate.B | present | absent | 14 | 0 | 59 | 0.44 | 9200 | 0.47 |
| EnvBac_Water_Wchmk_H10_scBac_plate.B | absent | absent | 24 | 0.01 | 67 | 0.41 | 7500 | 0.34 |
| EnvBac_Water_WGAX_A11_scBac_plate.B | present | present | 18 | 0 | 69 | 0.8 | 22000 | 0.51 |
| EnvBac_Water_WGAX_B11_scBac_plate.B | absent | present | 15 | 0 | 52 | 0.49 | 12000 | 0.41 |
| EnvBac_Water_WGAX_C11_scBac_plate.B | present | absent | 11 | 0 | 45 | 0.38 | 13000 | 0.5 |
| EnvBac_Water_WGAX_D11_scBac_plate.B | absent | present | 11 | 0 | 28 | 0.25 | 13000 | 0.41 |
| EnvBac_Water_WGAX_E11_scBac_plate.B | absent | absent | 11 | 0 | 31 | 0.32 | 13000 | 0.46 |
| EnvBac_Water_WGAX_F11_scBac_plate.B | absent | absent | 4 | 0 | 8 | 0.07 | 15000 | 0.49 |
| EnvBac_Water_WGAX_G11_scBac_plate.B | absent | present | 7.6 | 0.04 | 34 | 0.18 | 5600 | 0.45 |
| EnvBac_Water_WGAX_H11_scBac_plate.B | absent | absent | 6.4 | 0 | 17 | 0.1 | 7600 | 0.54 |
| EnvBac_Water_WGAX_A12_scBac_plate.B | absent | absent | 17 | 0.01 | 41 | 0.46 | 15000 | 0.36 |
| EnvBac_Water_WGAX_B12_scBac_plate.B | absent | absent | 31 | 0.02 | 59 | 1 | 33000 | 0.46 |
| EnvBac_Water_WGAX_C12_scBac_plate.B | absent | absent | 24 | 0.01 | 53 | 0.66 | 21000 | 0.52 |
| EnvBac_Water_WGAX_D12_scBac_plate.B | present | absent | 12 | 0 | 35 | 0.32 | 16000 | 0.52 |
| EnvBac_Water_WGAX_F12_scBac_plate.B | absent | absent | 13 | 0 | 28 | 0.34 | 15000 | 0.45 |
| EnvBac_Water_WGAX_G12_scBac_plate.B | present | absent | 9.5 | 0 | 37 | 0.33 | 14000 | 0.46 |
| EnvBac_Water_WGAX_H12_scBac_plate.B | absent | present | 9.3 | 0 | 27 | 0.2 | 9700 | 0.46 |
| EnvBac_Soil_PTA_G02_scBac_plate.C | present | present | 22 | 3.2 | 320 | 1.4 | 5400 | 0.6 |
| EnvBac_Soil_PTA_B04_scBac_plate.C | present | absent | 54 | 1.7 | 220 | 2.2 | 16000 | 0.36 |

**Supplementary Table S5: (Continued).**

| genome_id | plasmid_gtr5kb | virus_gtr5kb | Completeness_(%) | Contamination_(%) | contig_count | genome_size_Mbp | Contig_N50 | GC_Content |
|---|---|---|---|---|---|---|---|---|
| EnvBac_Soil_PTA_F04_scBac_plate.C | present | absent | 28 | 0.02 | 300 | 1.3 | 5100 | 0.65 |
| EnvBac_Soil_PTA_A05_scBac_plate.C | present | present | 60 | 1.5 | 500 | 3.8 | 11000 | 0.63 |
| EnvBac_Soil_PTA_B05_scBac_plate.C | present | absent | 64 | 2.2 | 400 | 3.1 | 11000 | 0.6 |
| EnvBac_Soil_PTA_C05_scBac_plate.C | present | absent | 60 | 0.05 | 240 | 2.4 | 17000 | 0.6 |
| EnvBac_Soil_Wchmk_E07_scBac_plate.C | absent | absent | 12 | 0.56 | 59 | 0.53 | 12000 | 0.63 |
| EnvBac_Soil_Wchmk_B08_scBac_plate.C | present | absent | 8 | 0 | 78 | 0.54 | 8300 | 0.42 |
| EnvBac_Soil_WGAX_A10_scBac_plate.C | absent | absent | 9.5 | 0.02 | 28 | 0.37 | 28000 | 0.37 |
| EnvBac_Soil_WGAX_B10_scBac_plate.C | absent | absent | 5.2 | 0 | 1 | 0.01 | 7700 | 0.59 |
| EnvBac_Soil_WGAX_C10_scBac_plate.C | present | absent | 15 | 0.04 | 71 | 0.75 | 18000 | 0.65 |
| EnvBac_Soil_WGAX_D10_scBac_plate.C | absent | absent | 5.5 | 0.04 | 13 | 0.19 | 35000 | 0.66 |
| EnvBac_Soil_WGAX_E10_scBac_plate.C | absent | absent | 6.6 | 0 | 20 | 0.18 | 12000 | 0.64 |
| EnvBac_Soil_WGAX_F10_scBac_plate.C | absent | absent | 17 | 0.25 | 49 | 0.84 | 48000 | 0.68 |
| EnvBac_Soil_WGAX_C11_scBac_plate.C | absent | absent | 11 | 0 | 21 | 0.27 | 31000 | 0.4 |
| EnvBac_Soil_WGAX_D11_scBac_plate.C | absent | absent | 2.4 | 0.02 | 2 | 0.02 | 14000 | 0.62 |
| EnvBac_Water_edMDA_A04_scBac_plate.A | absent | absent | 15 | 0 | 54 | 0.42 | 10000 | 0.38 |
| EnvBac_Water_edMDA_B04_scBac_plate.A | absent | absent | 11 | 0 | 49 | 0.4 | 12000 | 0.38 |
| EnvBac_Water_edMDA_C04_scBac_plate.A | absent | present | 22 | 0.14 | 48 | 0.46 | 14000 | 0.47 |
| EnvBac_Water_edMDA_D04_scBac_plate.A | absent | absent | 19 | 0 | 70 | 0.66 | 15000 | 0.41 |
| EnvBac_Water_edMDA_F04_scBac_plate.A | present | absent | 10 | 0.01 | 30 | 0.28 | 13000 | 0.5 |
| EnvBac_Water_edMDA_G04_scBac_plate.A | absent | absent | 26 | 0.03 | 23 | 0.26 | 15000 | 0.35 |
| EnvBac_Water_edMDA_H04_scBac_plate.A | absent | absent | 17 | 0 | 27 | 0.28 | 13000 | 0.34 |
| EnvBac_Water_edMDA_A05_scBac_plate.A | absent | absent | 8.6 | 0 | 30 | 0.22 | 10000 | 0.45 |
| EnvBac_Water_edMDA_C05_scBac_plate.A | present | absent | 25 | 0.01 | 54 | 0.58 | 16000 | 0.47 |
| EnvBac_Water_edMDA_E05_scBac_plate.A | absent | absent | 13 | 0 | 38 | 0.39 | 14000 | 0.47 |
| EnvBac_Water_edMDA_F05_scBac_plate.A | absent | absent | 20 | 0 | 54 | 0.4 | 11000 | 0.49 |
| EnvBac_Water_edMDA_G05_scBac_plate.A | present | absent | 8.4 | 0 | 32 | 0.23 | 10000 | 0.57 |
| EnvBac_Water_edMDA_H05_scBac_plate.A | absent | absent | 8.6 | 0.01 | 33 | 0.28 | 12000 | 0.45 |
| EnvBac_Water_edMDA_A06_scBac_plate.A | absent | absent | 53 | 0.09 | 35 | 0.65 | 33000 | 0.3 |
| EnvBac_Water_edMDA_B06_scBac_plate.A | absent | absent | 13 | 0 | 35 | 0.26 | 10000 | 0.43 |
| EnvBac_Water_edMDA_C06_scBac_plate.A | present | absent | 14 | 0 | 55 | 0.35 | 7400 | 0.4 |
| EnvBac_Water_edMDA_E06_scBac_plate.A | absent | absent | 16 | 0.01 | 43 | 0.31 | 9500 | 0.4 |
| EnvBac_Water_edMDA_F06_scBac_plate.A | present | present | 22 | 0 | 60 | 0.49 | 12000 | 0.39 |
| EnvBac_Water_edMDA_G06_scBac_plate.A | absent | absent | 7.1 | 0 | 18 | 0.2 | 14000 | 0.6 |
| EnvBac_Water_edMDA_H06_scBac_plate.A | absent | absent | 42 | 0.06 | 65 | 0.62 | 13000 | 0.29 |
| EnvBac_Soil_edMDA_A08_scBac_plate.A | present | absent | 25 | 0.13 | 89 | 1.3 | 32000 | 0.62 |
| EnvBac_Soil_edMDA_B08_scBac_plate.A | present | absent | 7.9 | 0.04 | 25 | 0.37 | 27000 | 0.63 |
| EnvBac_Soil_edMDA_D08_scBac_plate.A | absent | absent | 4.6 | 0 | 12 | 0.19 | 46000 | 0.65 |
| EnvBac_Soil_edMDA_E08_scBac_plate.A | absent | absent | 6.9 | 0.01 | 26 | 0.3 | 18000 | 0.42 |
| EnvBac_Soil_edMDA_F08_scBac_plate.A | present | absent | 23 | 0.02 | 53 | 0.8 | 38000 | 0.66 |
| EnvBac_Soil_edMDA_B09_scBac_plate.A | present | absent | 47 | 0 | 90 | 1.5 | 35000 | 0.61 |
| EnvBac_Soil_edMDA_C09_scBac_plate.A | absent | absent | 6.4 | 0.02 | 23 | 0.22 | 20000 | 0.65 |
| EnvBac_Soil_edMDA_D09_scBac_plate.A | absent | absent | 8.9 | 0 | 40 | 0.43 | 26000 | 0.64 |
| EnvBac_Soil_edMDA_E09_scBac_plate.A | absent | absent | 3 | 0 | 1 | 0.05 | 53000 | 0.64 |
| EnvBac_Soil_edMDA_F09_scBac_plate.A | absent | present | 11 | 0 | 36 | 0.35 | 14000 | 0.47 |
| EnvBac_Soil_edMDA_G09_scBac_plate.A | absent | absent | 6.2 | 0.26 | 38 | 0.2 | 6100 | 0.44 |
| EnvBac_Water_PTA_D06_scBac_plate.B | present | present | 100 | 3.4 | 100 | 3.3 | 150000 | 0.51 |
| EnvBac_Water_WGAX_E12_scBac_plate.B | present | absent | 34 | 0.03 | 72 | 0.9 | 18000 | 0.45 |
| EnvBac_Water_PTA_A03_scBac_plate.B | present | present | 95 | 8.3 | 210 | 3.3 | 43000 | 0.62 |
| EnvBac_Water_PTA_B03_scBac_plate.B | present | present | 81 | 9.8 | 200 | 2.3 | 29000 | 0.38 |
| EnvBac_Water_PTA_C03_scBac_plate.B | present | present | 74 | 7.3 | 160 | 1.6 | 49000 | 0.55 |
| EnvBac_Water_PTA_D03_scBac_plate.B | present | present | 89 | 6.5 | 260 | 2.7 | 18000 | 0.41 |
| EnvBac_Water_PTA_E03_scBac_plate.B | absent | present | 73 | 6.7 | 250 | 2.2 | 14000 | 0.63 |
| EnvBac_Water_PTA_F03_scBac_plate.B | present | absent | 77 | 6.7 | 150 | 1.6 | 47000 | 0.34 |
| EnvBac_Water_PTA_G03_scBac_plate.B | present | absent | 65 | 2.1 | 99 | 1.2 | 89000 | 0.43 |
| EnvBac_Water_PTA_H03_scBac_plate.B | present | present | 77 | 6 | 160 | 1.6 | 57000 | 0.38 |
| EnvBac_Water_PTA_A04_scBac_plate.B | present | absent | 85 | 10 | 220 | 1.5 | 11000 | 0.32 |
| EnvBac_Water_PTA_B04_scBac_plate.B | present | absent | 85 | 6.4 | 110 | 1.7 | 2e+05 | 0.35 |
| EnvBac_Water_PTA_C04_scBac_plate.B | present | present | 55 | 11 | 150 | 1.2 | 12000 | 0.42 |
| EnvBac_Water_PTA_D04_scBac_plate.B | present | absent | 97 | 8.4 | 160 | 2.6 | 57000 | 0.4 |
| EnvBac_Water_PTA_E04_scBac_plate.B | present | absent | 100 | 6.4 | 130 | 2.6 | 72000 | 0.37 |
| EnvBac_Water_PTA_F04_scBac_plate.B | present | present | 97 | 7.1 | 160 | 1.6 | 32000 | 0.33 |
| EnvBac_Water_PTA_G04_scBac_plate.B | present | present | 85 | 8.3 | 200 | 2 | 17000 | 0.47 |
| EnvBac_Water_PTA_H04_scBac_plate.B | present | absent | 89 | 8.2 | 130 | 1.7 | 280000 | 0.37 |
| EnvBac_Water_PTA_A05_scBac_plate.B | present | present | 93 | 3.6 | 93 | 1.5 | 940000 | 0.52 |
| EnvBac_Water_PTA_B05_scBac_plate.B | present | present | 100 | 4.1 | 100 | 3.3 | 2800000 | 0.56 |
| EnvBac_Water_PTA_C05_scBac_plate.B | present | present | 96 | 4.8 | 190 | 3.1 | 58000 | 0.49 |
| EnvBac_Water_PTA_D05_scBac_plate.B | present | absent | 82 | 1.9 | 150 | 2.4 | 28000 | 0.4 |
| EnvBac_Water_PTA_E05_scBac_plate.B | present | present | 100 | 3 | 180 | 3 | 40000 | 0.49 |
| EnvBac_Water_PTA_F05_scBac_plate.B | present | present | 100 | 6.7 | 160 | 2.5 | 47000 | 0.45 |
| EnvBac_Water_PTA_G05_scBac_plate.B | present | present | 100 | 8.3 | 140 | 2.9 | 130000 | 0.53 |
| EnvBac_Water_PTA_H05_scBac_plate.B | present | absent | 94 | 6.3 | 140 | 2.8 | 51000 | 0.42 |
| EnvBac_Water_PTA_A06_scBac_plate.B | present | present | 82 | 12 | 270 | 2.6 | 20000 | 0.44 |
| EnvBac_Water_PTA_B06_scBac_plate.B | absent | absent | 89 | 9.1 | 190 | 1.8 | 27000 | 0.32 |
| EnvBac_Water_PTA_C06_scBac_plate.B | absent | absent | 76 | 6.1 | 120 | 1.8 | 77000 | 0.45 |
| EnvBac_Water_Wchmk_A09_scBac_plate.B | present | absent | 56 | 0.02 | 85 | 0.73 | 13000 | 0.3 |
| EnvBac_Water_Wchmk_B09_scBac_plate.B | absent | absent | 41 | 0.08 | 100 | 0.93 | 14000 | 0.4 |
| EnvBac_Water_Wchmk_C09_scBac_plate.B | absent | present | 23 | 0.23 | 140 | 0.79 | 7100 | 0.42 |
| EnvBac_Water_Wchmk_D09_scBac_plate.B | absent | absent | 48 | 0.2 | 79 | 0.6 | 9500 | 0.36 |
| EnvBac_Water_Wchmk_E09_scBac_plate.B | absent | absent | 40 | 0.21 | 120 | 0.96 | 11000 | 0.35 |
| EnvBac_Water_Wchmk_F09_scBac_plate.B | present | present | 30 | 0.33 | 89 | 0.6 | 8700 | 0.49 |

**Supplementary Table S5: (Continued).**

| genome_id | plasmid_gtr5kb | virus_gtr5kb | Completeness_(%) | Contamination_(%) | contig_count | genome_size_Mbp | Contig_N50 | GC_Content |
|---|---|---|---|---|---|---|---|---|
| EnvBac_Water_Wchmk_G09_scBac_plate.B | absent | absent | 34 | 0.54 | 83 | 0.76 | 14000 | 0.38 |
| EnvBac_Water_Wchmk_H09_scBac_plate.B | absent | present | 30 | 0.18 | 97 | 0.78 | 10000 | 0.45 |
| EnvBac_Water_Wchmk_A10_scBac_plate.B | present | absent | 16 | 0.03 | 49 | 0.4 | 12000 | 0.58 |
| EnvBac_Water_Wchmk_B10_scBac_plate.B | present | absent | 19 | 0.44 | 69 | 0.48 | 9400 | 0.46 |
| EnvBac_Water_Wchmk_C10_scBac_plate.B | present | absent | 23 | 0.03 | 110 | 0.83 | 10000 | 0.49 |
| EnvBac_Water_Wchmk_D10_scBac_plate.B | absent | absent | 51 | 0.04 | 96 | 1.1 | 18000 | 0.41 |
| EnvBac_Water_Wchmk_E10_scBac_plate.B | present | absent | 21 | 0.06 | 73 | 0.64 | 16000 | 0.56 |
| EnvBac_Water_Wchmk_F10_scBac_plate.B | absent | absent | 26 | 0.11 | 75 | 0.53 | 8900 | 0.47 |
| EnvBac_Water_Wchmk_G10_scBac_plate.B | present | absent | 20 | 0.17 | 100 | 0.77 | 9600 | 0.48 |
| EnvBac_Water_Wchmk_H10_scBac_plate.B | absent | absent | 38 | 0.76 | 89 | 0.57 | 9100 | 0.35 |
| EnvBac_Water_WGAX_A11_scBac_plate.B | present | present | 49 | 0.72 | 110 | 1.5 | 22000 | 0.51 |
| EnvBac_Water_WGAX_B11_scBac_plate.B | absent | present | 27 | 0 | 110 | 0.83 | 9000 | 0.41 |
| EnvBac_Water_WGAX_C11_scBac_plate.B | present | absent | 24 | 0 | 88 | 0.76 | 11000 | 0.5 |
| EnvBac_Water_WGAX_D11_scBac_plate.B | absent | present | 19 | 0.52 | 60 | 0.49 | 12000 | 0.42 |
| EnvBac_Water_WGAX_E11_scBac_plate.B | absent | absent | 21 | 0.05 | 72 | 0.64 | 13000 | 0.46 |
| EnvBac_Water_WGAX_F11_scBac_plate.B | absent | absent | 8.8 | 0 | 37 | 0.21 | 6800 | 0.49 |
| EnvBac_Water_WGAX_G11_scBac_plate.B | absent | absent | 10 | 0.43 | 59 | 0.31 | 5500 | 0.46 |
| EnvBac_Water_WGAX_H11_scBac_plate.B | absent | absent | 10 | 0 | 25 | 0.14 | 7600 | 0.54 |
| EnvBac_Water_WGAX_A12_scBac_plate.B | absent | absent | 31 | 0 | 71 | 0.81 | 17000 | 0.36 |
| EnvBac_Water_WGAX_B12_scBac_plate.B | absent | present | 58 | 0.09 | 79 | 1.6 | 30000 | 0.46 |
| EnvBac_Water_WGAX_C12_scBac_plate.B | present | absent | 35 | 0.04 | 89 | 1.1 | 20000 | 0.53 |
| EnvBac_Water_WGAX_D12_scBac_plate.B | present | absent | 25 | 0.05 | 55 | 0.72 | 19000 | 0.54 |
| EnvBac_Water_WGAX_F12_scBac_plate.B | absent | absent | 19 | 0.01 | 51 | 0.53 | 13000 | 0.45 |
| EnvBac_Water_WGAX_G12_scBac_plate.B | present | absent | 16 | 0.03 | 67 | 0.57 | 13000 | 0.46 |
| EnvBac_Water_WGAX_H12_scBac_plate.B | present | present | 15 | 0 | 47 | 0.34 | 9700 | 0.46 |
| EnvBac_Soil_PTA_G02_scBac_plate.C | present | present | 58 | 9.6 | 610 | 4.2 | 9400 | 0.63 |
| EnvBac_Soil_PTA_B04_scBac_plate.C | present | present | 76 | 9.5 | 340 | 3.3 | 15000 | 0.37 |
| EnvBac_Soil_PTA_F04_scBac_plate.C | present | absent | 81 | 1.6 | 340 | 5.2 | 31000 | 0.67 |
| EnvBac_Soil_PTA_A05_scBac_plate.C | present | absent | 88 | 10 | 390 | 6.4 | 36000 | 0.63 |
| EnvBac_Soil_PTA_B05_scBac_plate.C | present | present | 84 | 8.4 | 400 | 5 | 22000 | 0.61 |
| EnvBac_Soil_PTA_C05_scBac_plate.C | present | present | 85 | 3 | 300 | 3.6 | 27000 | 0.6 |
| EnvBac_Soil_Wchmk_E07_scBac_plate.C | absent | absent | 19 | 0.79 | 100 | 0.98 | 13000 | 0.64 |
| EnvBac_Soil_Wchmk_B08_scBac_plate.C | present | absent | 14 | 0.02 | 150 | 1 | 8500 | 0.43 |
| EnvBac_Soil_WGAX_A10_scBac_plate.C | absent | present | 20 | 0.22 | 52 | 0.68 | 19000 | 0.37 |
| EnvBac_Soil_WGAX_B10_scBac_plate.C | absent | absent | 4.9 | 0.03 | 17 | 0.14 | 10000 | 0.66 |
| EnvBac_Soil_WGAX_C10_scBac_plate.C | present | absent | 26 | 0.14 | 110 | 1.4 | 22000 | 0.65 |
| EnvBac_Soil_WGAX_D10_scBac_plate.C | absent | absent | 12 | 0 | 40 | 0.51 | 36000 | 0.66 |
| EnvBac_Soil_WGAX_E10_scBac_plate.C | present | absent | 6.5 | 0.04 | 42 | 0.35 | 13000 | 0.64 |
| EnvBac_Soil_WGAX_F10_scBac_plate.C | absent | absent | 33 | 0.04 | 65 | 1.6 | 65000 | 0.68 |
| EnvBac_Soil_WGAX_C11_scBac_plate.C | present | absent | 17 | 0 | 33 | 0.39 | 19000 | 0.4 |
| EnvBac_Soil_WGAX_D11_scBac_plate.C | present | absent | 4.6 | 0.01 | 15 | 0.16 | 14000 | 0.62 |
| EnvBac_Water_edMDA_A04_scBac_plate.A | absent | absent | 36 | 0.06 | 99 | 0.72 | 9800 | 0.38 |
| EnvBac_Water_edMDA_B04_scBac_plate.A | absent | absent | 24 | 0.18 | 83 | 0.7 | 12000 | 0.38 |
| EnvBac_Water_edMDA_C04_scBac_plate.A | absent | present | 38 | 0.23 | 78 | 0.76 | 14000 | 0.47 |
| EnvBac_Water_edMDA_D04_scBac_plate.A | absent | present | 37 | 0.02 | 98 | 1.1 | 16000 | 0.41 |
| EnvBac_Water_edMDA_F04_scBac_plate.A | present | absent | 23 | 2.4 | 55 | 0.58 | 14000 | 0.5 |
| EnvBac_Water_edMDA_G04_scBac_plate.A | absent | absent | 29 | 0.01 | 36 | 0.39 | 19000 | 0.35 |
| EnvBac_Water_edMDA_H04_scBac_plate.A | absent | absent | 31 | 0.27 | 49 | 0.44 | 14000 | 0.34 |
| EnvBac_Water_edMDA_A05_scBac_plate.A | absent | absent | 16 | 0.05 | 72 | 0.53 | 8900 | 0.45 |
| EnvBac_Water_edMDA_C05_scBac_plate.A | present | absent | 44 | 0.01 | 100 | 0.97 | 15000 | 0.47 |
| EnvBac_Water_edMDA_E05_scBac_plate.A | absent | present | 23 | 0.03 | 75 | 0.65 | 12000 | 0.46 |
| EnvBac_Water_edMDA_F05_scBac_plate.A | present | absent | 30 | 0.01 | 91 | 0.66 | 9400 | 0.5 |
| EnvBac_Water_edMDA_G05_scBac_plate.A | present | absent | 11 | 0.01 | 60 | 0.41 | 8700 | 0.58 |
| EnvBac_Water_edMDA_H05_scBac_plate.A | absent | absent | 24 | 1.8 | 58 | 0.6 | 13000 | 0.45 |
| EnvBac_Water_edMDA_A06_scBac_plate.A | absent | absent | 75 | 0.43 | 47 | 0.93 | 34000 | 0.3 |
| EnvBac_Water_edMDA_B06_scBac_plate.A | absent | absent | 22 | 0.07 | 65 | 0.43 | 9000 | 0.44 |
| EnvBac_Water_edMDA_C06_scBac_plate.A | absent | absent | 23 | 1 | 92 | 0.64 | 8800 | 0.4 |
| EnvBac_Water_edMDA_E06_scBac_plate.A | absent | absent | 30 | 2 | 88 | 0.58 | 9000 | 0.4 |
| EnvBac_Water_edMDA_F06_scBac_plate.A | present | present | 41 | 1.6 | 99 | 0.83 | 12000 | 0.39 |
| EnvBac_Water_edMDA_G06_scBac_plate.A | absent | absent | 11 | 0 | 39 | 0.39 | 13000 | 0.61 |
| EnvBac_Water_edMDA_H06_scBac_plate.A | absent | absent | 68 | 1.2 | 74 | 0.96 | 22000 | 0.29 |
| EnvBac_Soil_edMDA_A08_scBac_plate.A | present | absent | 52 | 0.62 | 140 | 2.6 | 40000 | 0.63 |
| EnvBac_Soil_edMDA_B08_scBac_plate.A | present | absent | 19 | 0.08 | 81 | 1.1 | 26000 | 0.64 |
| EnvBac_Soil_edMDA_D08_scBac_plate.A | absent | absent | 12 | 0 | 48 | 0.59 | 29000 | 0.65 |
| EnvBac_Soil_edMDA_E08_scBac_plate.A | absent | absent | 7.6 | 0 | 34 | 0.37 | 18000 | 0.42 |
| EnvBac_Soil_edMDA_F08_scBac_plate.A | present | present | 37 | 0.32 | 84 | 1.4 | 43000 | 0.66 |
| EnvBac_Soil_edMDA_B09_scBac_plate.A | present | absent | 60 | 5 | 81 | 2.5 | 67000 | 0.61 |
| EnvBac_Soil_edMDA_C09_scBac_plate.A | absent | absent | 7 | 0.02 | 25 | 0.22 | 14000 | 0.65 |
| EnvBac_Soil_edMDA_D09_scBac_plate.A | absent | absent | 24 | 0.05 | 87 | 1.2 | 35000 | 0.65 |
| EnvBac_Soil_edMDA_E09_scBac_plate.A | present | absent | 8.2 | 0 | 22 | 0.22 | 15000 | 0.65 |
| EnvBac_Soil_edMDA_F09_scBac_plate.A | absent | present | 13 | 0.24 | 62 | 0.53 | 12000 | 0.47 |
| EnvBac_Soil_edMDA_G09_scBac_plate.A | absent | absent | 15 | 0.27 | 56 | 0.33 | 7600 | 0.44 |
