## Supplementary_Table_S6 for "scMicrobe PTA: Near Complete Genomes from Single Bacterial Cells"

**Supplementary Table S6. Metadata for individual isolate SAG sequencing and overview of de novo assembly results.**

| genome_id | seq_platform | BioProject | BioSample | amp_method | sample_type | library_name | Enzymatic Lysis | misag | rrna_16S_hit | plasmid_gtr5kb |
|---|---|---|---|---|---|---|---|---|---|---|
| scBac_PTA_Bsubtilis_sc_plate2well5A_S66 | novseq 2000 | PRJNA1067728 | SAMN39550258 | PTA | Bsubtilis | not_assigned | TRUE | HQ | absent | present |
| scBac_PTA_Bsubtilis_sc_plate2well5B_S67 | novseq 2000 | PRJNA1067728 | SAMN39550259 | PTA | Bsubtilis | not_assigned | TRUE | HQ | absent | present |
| scBac_PTA_Bsubtilis_sc_plate2well5D_S68 | novseq 2000 | PRJNA1067728 | SAMN39550260 | PTA | Bsubtilis | not_assigned | TRUE | HQ | absent | present |
| scBac_PTA_Bsubtilis_sc_plate2well5G_S69 | novseq 2000 | PRJNA1067728 | SAMN39550261 | PTA | Bsubtilis | not_assigned | TRUE | HQ | absent | present |
| scBac_PTA_Ecoli_sc_plate2well4A_S62 | novseq 2000 | PRJNA1067728 | SAMN39550250 | PTA | Ecoli | not_assigned | TRUE | HQ | absent | present |
| scBac_PTA_Ecoli_sc_plate2well4B_S63 | novseq 2000 | PRJNA1067728 | SAMN39550251 | PTA | Ecoli | not_assigned | TRUE | MQ | absent | present |
| scBac_PTA_Ecoli_sc_plate2well4D_S64 | novseq 2000 | PRJNA1067728 | SAMN39550252 | PTA | Ecoli | not_assigned | TRUE | HQ | absent | present |
| scBac_PTA_Ecoli_sc_plate2well4E_S65 | novseq 2000 | PRJNA1067728 | SAMN39550253 | PTA | Ecoli | not_assigned | TRUE | HQ | absent | present |
| scBac_PTA_Pputida_sc_plate2well3A_S58 | novseq 2000 | PRJNA1067728 | SAMN39550254 | PTA | Pputida | not_assigned | TRUE | MQ | present | present |
| scBac_PTA_Pputida_sc_plate2well3B_S59 | novseq 2000 | PRJNA1067728 | SAMN39550255 | PTA | Pputida | not_assigned | TRUE | MQ | absent | present |
| scBac_PTA_Pputida_sc_plate2well3D_S60 | novseq 2000 | PRJNA1067728 | SAMN39550256 | PTA | Pputida | not_assigned | TRUE | MQ | absent | present |
| scBac_PTA_Pputida_sc_plate2well3E_S61 | novseq 2000 | PRJNA1067728 | SAMN39550257 | PTA | Pputida | not_assigned | TRUE | MQ | absent | present |
| scBac_WGAX_Ecoli_sc_well7B_S42 | novseq 2000 | PRJNA1067728 | SAMN39550285 | WGAX | Ecoli | not_assigned | FALSE | LQ | absent | present |
| scBac_WGAX_Ecoli_sc_well7C_S43 | novseq 2000 | PRJNA1067728 | SAMN39550286 | WGAX | Ecoli | not_assigned | FALSE | MQ | absent | present |
| scBac_WGAX_Ecoli_sc_well7F_S44 | novseq 2000 | PRJNA1067728 | SAMN39550287 | WGAX | Ecoli | not_assigned | FALSE | LQ | absent | present |
| scBac_WGAX_Ecoli_sc_well7G_S45 | novseq 2000 | PRJNA1067728 | SAMN39550288 | WGAX | Ecoli | not_assigned | FALSE | LQ | absent | present |
| scBac_WGAX_Pputida_sc_well3D_S38 | novseq 2000 | PRJNA1067728 | SAMN39550289 | WGAX | Pputida | not_assigned | FALSE | LQ | absent | present |
| scBac_WGAX_Pputida_sc_well3F_S39 | novseq 2000 | PRJNA1067728 | SAMN39550290 | WGAX | Pputida | not_assigned | FALSE | LQ | absent | present |
| scBac_WGAX_Pputida_sc_well3G_S40 | novseq 2000 | PRJNA1067728 | SAMN39550291 | WGAX | Pputida | not_assigned | FALSE | LQ | absent | present |
| scBac_WGAX_Pputida_sc_well3H_S41 | novseq 2000 | PRJNA1067728 | SAMN39550292 | WGAX | Pputida | not_assigned | FALSE | LQ | present | present |
| scBac_Watchmaker_Bsubtilis_sc_well7C_S32 | novseq 2000 | PRJNA1067728 | SAMN39550271 | MDA | Bsubtilis | not_assigned | FALSE | LQ | absent | present |
| scBac_Watchmaker_Bsubtilis_sc_well7H_S33 | novseq 2000 | PRJNA1067728 | SAMN39550272 | MDA | Bsubtilis | not_assigned | FALSE | MQ | absent | present |
| scBac_Watchmaker_Ecoli_sc_well6A_S28 | novseq 2000 | PRJNA1067728 | SAMN39550267 | MDA | Ecoli | not_assigned | FALSE | MQ | absent | present |
| scBac_Watchmaker_Ecoli_sc_well6C_S29 | novseq 2000 | PRJNA1067728 | SAMN39550268 | MDA | Ecoli | not_assigned | FALSE | MQ | absent | present |
| scBac_Watchmaker_Ecoli_sc_well6E_S30 | novseq 2000 | PRJNA1067728 | SAMN39550269 | MDA | Ecoli | not_assigned | FALSE | MQ | absent | present |
| scBac_Watchmaker_Ecoli_sc_well6H_S31 | novseq 2000 | PRJNA1067728 | SAMN39550270 | MDA | Ecoli | not_assigned | FALSE | MQ | absent | present |
| scBac_Watchmaker_Pputida_sc_well3A_S24 | novseq 2000 | PRJNA1067728 | SAMN39550281 | MDA | Pputida | not_assigned | FALSE | LQ | absent | present |
| scBac_Watchmaker_Pputida_sc_well3C_S25 | novseq 2000 | PRJNA1067728 | SAMN39550282 | MDA | Pputida | not_assigned | FALSE | LQ | absent | present |
| scBac_Watchmaker_Pputida_sc_well3D_S26 | novseq 2000 | PRJNA1067728 | SAMN39550283 | MDA | Pputida | not_assigned | FALSE | LQ | present | present |
| scBac_Watchmaker_Pputida_sc_well3E_S27 | novseq 2000 | PRJNA1067728 | SAMN39550284 | MDA | Pputida | not_assigned | FALSE | LQ | absent | present |

**Supplementary Table S6. (Continued).**

| genome_id | virus_gtr5kb | Genome_fraction | Completeness | Contamination | gini | contig_count | genome_size_Mbp | Contig_N50 | Largest_contig | missassem |
|---|---|---|---|---|---|---|---|---|---|---|
| scBac_PTA_Bsubtilis_sc_plate2well5A_S66 | present | 96 | 100 | 0.11 | 0.33 | 92 | 4.1 | 78000 | 250000 | 3 |
| scBac_PTA_Bsubtilis_sc_plate2well5B_S67 | present | 93 | 92 | 1.2 | 0.36 | 110 | 3.9 | 91000 | 430000 | 6 |
| scBac_PTA_Bsubtilis_sc_plate2well5D_S68 | present | 96 | 98 | 0.17 | 0.34 | 83 | 4 | 150000 | 320000 | 3 |
| scBac_PTA_Bsubtilis_sc_plate2well5G_S69 | present | 91 | 92 | 0.24 | 0.38 | 170 | 3.9 | 45000 | 230000 | 7 |
| scBac_PTA_Ecoli_sc_plate2well4A_S62 | present | 93 | 100 | 3.6 | 0.36 | 290 | 4.8 | 36000 | 160000 | 1 |
| scBac_PTA_Ecoli_sc_plate2well4B_S63 | present | 87 | 87 | 0.33 | 0.4 | 220 | 4.1 | 39000 | 180000 | 3 |
| scBac_PTA_Ecoli_sc_plate2well4D_S64 | present | 95 | 100 | 0.07 | 0.31 | 140 | 4.4 | 52000 | 260000 | 0 |
| scBac_PTA_Ecoli_sc_plate2well4E_S65 | present | 90 | 94 | 0.71 | 0.37 | 190 | 4.2 | 37000 | 180000 | 1 |
| scBac_PTA_Pputida_sc_plate2well3A_S58 | present | 58 | 65 | 0.24 | 0.62 | 540 | 3.7 | 9000 | 94000 | 15 |
| scBac_PTA_Pputida_sc_plate2well3B_S59 | present | 70 | 77 | 0.24 | 0.59 | 540 | 4.4 | 12000 | 97000 | 16 |
| scBac_PTA_Pputida_sc_plate2well3D_S60 | present | 53 | 54 | 3.2 | 0.65 | 540 | 3.4 | 7800 | 57000 | 16 |
| scBac_PTA_Pputida_sc_plate2well3E_S61 | present | 51 | 60 | 0.28 | 0.63 | 510 | 3.2 | 8400 | 71000 | 14 |
| scBac_WGAX_Ecoli_sc_well7B_S42 | present | 41 | 42 | 0.01 | 0.81 | 120 | 2.1 | 38000 | 130000 | 5 |
| scBac_WGAX_Ecoli_sc_well7C_S43 | present | 57 | 56 | 0 | 0.74 | 120 | 2.6 | 50000 | 140000 | 3 |
| scBac_WGAX_Ecoli_sc_well7F_S44 | absent | 39 | 42 | 0.05 | 0.84 | 94 | 1.8 | 35000 | 140000 | 4 |
| scBac_WGAX_Ecoli_sc_well7G_S45 | present | 19 | 19 | 0 | 0.93 | 33 | 0.87 | 54000 | 120000 | 0 |
| scBac_WGAX_Pputida_sc_well3D_S38 | present | 15 | 18 | 0.04 | 0.95 | 94 | 0.92 | 24000 | 110000 | 1 |
| scBac_WGAX_Pputida_sc_well3F_S39 | present | 20 | 23 | 0.02 | 0.94 | 110 | 1.2 | 23000 | 120000 | 6 |
| scBac_WGAX_Pputida_sc_well3G_S40 | present | 13 | 11 | 0 | 0.96 | 60 | 0.82 | 33000 | 99000 | 4 |
| scBac_WGAX_Pputida_sc_well3H_S41 | absent | 5.9 | 5.3 | 0 | 0.99 | 23 | 0.37 | 45000 | 83000 | 2 |
| scBac_Watchmaker_Bsubtilis_sc_well7C_S32 | present | 50 | 49 | 0.02 | 0.8 | 100 | 2.1 | 40000 | 110000 | 11 |
| scBac_Watchmaker_Bsubtilis_sc_well7H_S33 | present | 69 | 74 | 0.06 | 0.64 | 140 | 2.9 | 50000 | 140000 | 10 |
| scBac_Watchmaker_Ecoli_sc_well6A_S28 | present | 64 | 71 | 3 | 0.65 | 260 | 3.8 | 34000 | 180000 | 8 |
| scBac_Watchmaker_Ecoli_sc_well6C_S29 | present | 55 | 53 | 0.09 | 0.73 | 120 | 2.5 | 48000 | 140000 | 11 |
| scBac_Watchmaker_Ecoli_sc_well6E_S30 | present | 68 | 69 | 0 | 0.65 | 150 | 3.2 | 61000 | 160000 | 7 |
| scBac_Watchmaker_Ecoli_sc_well6H_S31 | present | 63 | 66 | 0 | 0.7 | 160 | 2.9 | 39000 | 170000 | 6 |
| scBac_Watchmaker_Pputida_sc_well3A_S24 | present | 44 | 48 | 0.09 | 0.83 | 230 | 2.7 | 27000 | 2e+05 | 5 |
| scBac_Watchmaker_Pputida_sc_well3C_S25 | present | 27 | 31 | 0.41 | 0.9 | 140 | 1.7 | 26000 | 73000 | 10 |
| scBac_Watchmaker_Pputida_sc_well3D_S26 | present | 32 | 34 | 0.03 | 0.87 | 160 | 2 | 24000 | 71000 | 7 |
| scBac_Watchmaker_Pputida_sc_well3E_S27 | absent | 27 | 31 | 0.04 | 0.89 | 120 | 1.7 | 35000 | 130000 | 5 |

**Supplementary Table S6. (Continued).**

| genome_id | missassem_length | Unaligned_length | Duplication_ratio | N_s_per_100_kbp | mm_per_100_kbp | indels_per_100_kbp |
|---|---|---|---|---|---|---|
| scBac_PTA_Bsubtilis_sc_plate2well5A_S66 | 330000 | 5300 | 1 | 0 | 2 | 11 |
| scBac_PTA_Bsubtilis_sc_plate2well5B_S67 | 730000 | 8600 | 1 | 0 | 3.3 | 11 |
| scBac_PTA_Bsubtilis_sc_plate2well5D_S68 | 320000 | 4900 | 1 | 0 | 9.6 | 11 |
| scBac_PTA_Bsubtilis_sc_plate2well5G_S69 | 140000 | 4300 | 1 | 0 | 9.5 | 12 |
| scBac_PTA_Ecoli_sc_plate2well4A_S62 | 3200 | 5e+05 | 1 | 0 | 2.1 | 0.42 |
| scBac_PTA_Ecoli_sc_plate2well4B_S63 | 47000 | 8400 | 1 | 0 | 6 | 0.94 |
| scBac_PTA_Ecoli_sc_plate2well4D_S64 | 0 | 0 | 1 | 0 | 1.9 | 0.34 |
| scBac_PTA_Ecoli_sc_plate2well4E_S65 | 5300 | 2100 | 1 | 0 | 2.6 | 1.1 |
| scBac_PTA_Pputida_sc_plate2well3A_S58 | 120000 | 62000 | 1 | 0 | 15 | 1.4 |
| scBac_PTA_Pputida_sc_plate2well3B_S59 | 130000 | 29000 | 1 | 0 | 8 | 0.94 |
| scBac_PTA_Pputida_sc_plate2well3D_S60 | 84000 | 1e+05 | 1 | 0 | 10 | 1.7 |
| scBac_PTA_Pputida_sc_plate2well3E_S61 | 140000 | 51000 | 1 | 0 | 11 | 1.6 |
| scBac_WGAX_Ecoli_sc_well7B_S42 | 98000 | 170000 | 1 | 0 | 4 | 0.32 |
| scBac_WGAX_Ecoli_sc_well7C_S43 | 86000 | 0 | 1 | 0 | 3 | 0.49 |
| scBac_WGAX_Ecoli_sc_well7F_S44 | 55000 | 0 | 1 | 0 | 4.3 | 0.88 |
| scBac_WGAX_Ecoli_sc_well7G_S45 | 0 | 0 | 1 | 0 | 4.5 | 0.23 |
| scBac_WGAX_Pputida_sc_well3D_S38 | 3000 | 0 | 1 | 0 | 12 | 0.65 |
| scBac_WGAX_Pputida_sc_well3F_S39 | 83000 | 0 | 1 | 0 | 8.1 | 1 |
| scBac_WGAX_Pputida_sc_well3G_S40 | 55000 | 0 | 1 | 0 | 9.4 | 2.7 |
| scBac_WGAX_Pputida_sc_well3H_S41 | 28000 | 0 | 1 | 0 | 18 | 1.9 |
| scBac_Watchmaker_Bsubtilis_sc_well7C_S32 | 230000 | 0 | 1 | 0 | 6.4 | 12 |
| scBac_Watchmaker_Bsubtilis_sc_well7H_S33 | 260000 | 0 | 1 | 0 | 3.1 | 11 |
| scBac_Watchmaker_Ecoli_sc_well6A_S28 | 140000 | 840000 | 1 | 0 | 3.9 | 0.51 |
| scBac_Watchmaker_Ecoli_sc_well6C_S29 | 230000 | 0 | 1 | 0 | 3.3 | 0.35 |
| scBac_Watchmaker_Ecoli_sc_well6E_S30 | 130000 | 0 | 1 | 0 | 3.2 | 0.38 |
| scBac_Watchmaker_Ecoli_sc_well6H_S31 | 89000 | 0 | 1 | 0 | 3 | 0.41 |
| scBac_Watchmaker_Pputida_sc_well3A_S24 | 43000 | 0 | 1 | 0 | 12 | 1.2 |
| scBac_Watchmaker_Pputida_sc_well3C_S25 | 170000 | 0 | 1 | 0 | 15 | 1.6 |
| scBac_Watchmaker_Pputida_sc_well3D_S26 | 52000 | 0 | 1 | 0 | 5.7 | 0.5 |
| scBac_Watchmaker_Pputida_sc_well3E_S27 | 54000 | 0 | 1 | 0 | 9.6 | 0.83 |
